# FACS-Sortable Triple Emulsion Picoreactors for Screening Reactions in Biphasic Environments

**DOI:** 10.1101/2024.03.01.583031

**Authors:** Samuel Thompson, Yanrong Zhang, Zijian Yang, Lisa A. Nichols, Polly M. Fordyce

**Author notes:** These authors contributed equally (Yanrong Zhang, Zijian Yang, Lisa A. Nichols).

## Abstract

Biphasic environments can enable successful chemical reactions where any single solvent results in poor substrate solubility or poor catalyst reactivity. For screening biphasic reactions at high-throughput, a platform based on microfluidic double emulsions could use widely available FACS (Fluorescence Activated Cell Sorting) machines to screen millions of picoliter reactors in a few hours. However, encapsulating biphasic reactions within double emulsions to form FACS-sortable droplet picoreactors requires optimized solvent phases and surfactants to produce triple emulsion droplets that are stable over multi-hour assays and compatible with desired reaction conditions. This work demonstrates such FACS-sortable triple emulsion picoreactors with a fluorocarbon shell and biphasic octanol-in-water core. First, surfactants were screened to stabilize octanol-in-water emulsions for the picoreactor core. With these optimized conditions, stable triple emulsion picoreactors were produced (>70% of droplets survived to 24 hours), and the ability to produce protein in the biphasic core was demonstrated via cell-free protein synthesis. Finally, triple emulsion picoreactors were sorted based on fluorescence using commercial FACS sorters at >100 Hz with 75-80% of droplets recovered. These triple emulsion picoreactors have potential for future screening bead-encoded catalyst libraries, including enzymes such as lipases for biofuel production.

---

Catalysts are essential to the modern chemical industry, and novel catalysts open up avenues to new medicines^1^, materials with novel properties^2^, and more sustainable production methods^3^. To optimize reaction properties such as yield and enantiomeric excess, catalysts must be screened for compatibility and performance with each desired substrate. However, identifying an effective catalyst can be challenging when the substrate and catalyst are differentially soluble, as low effective substrate/catalyst concentrations and catalyst inactivation can limit product formation. Examples of such differentially soluble reactions include reactions with polar/hydrophilic catalysts and hydrophobic substrates (*e*.*g*. enzymatic reactions for lipase catalyzed biofuel production^4^ or PETase mediated plastic recycling^5^, and for biphasic variations of nitration reactions^6^ and organometallic CC bond formations via olefin metathesis^7,8^ or Suzuki-Miyura coupling^9^).

This challenge can be circumvented using a biphasic reaction system with two immiscible phases (e.g. an aqueous phase containing most of the catalyst and a hydrocarbon phase containing most of the substrate and product) (**Fig. 1A**). Despite catalyst and substrate preferring opposite solvent phases, substrate, product, and catalyst can still exchange between the two phases such that overall product formation rates depend on the rates of catalysis within each phase, the equilibrium concentrations, and kinetics of partitioning (**Fig. 1A**). Hydrocarbon solvents with a variety of functional groups can form biphasic systems with polar and aqueous solvents, including alkanes, alcohols, ketones, and esters (**Fig. 1B**). This wide variety of functional groups enables many useful processes in biphasic reactions, including petrochemical oxidation^10^ and bioremediation^11^, plastic recycling^12,13^, biofuel production^4,14^, and natural product synthesis/extraction^15,16^. Critically, the presence of an aqueous phase allows the application and benefits of enzymatic processes: 1) high activity at ambient temperature and pressure, 3) generally higher turnover, 3) biodegradability, and 4) production costs that can fall below $10/kg of catalyst^17–19^.

**Figure 1:**
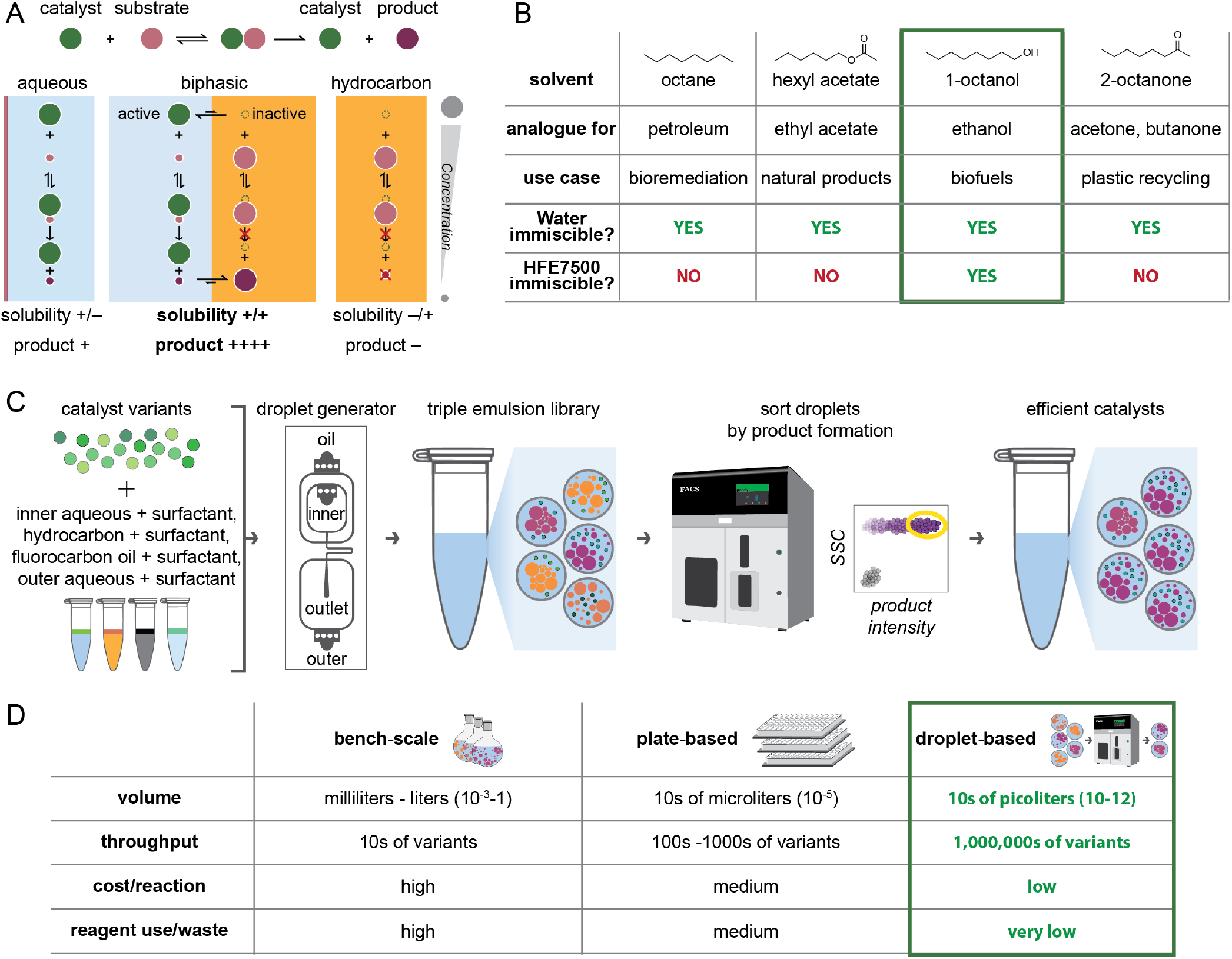
Triple emulsion picoreactors for optimizing biphasic reactions with wide-ranging applications. **A)** Schema illustrating how biphasic reactions use immiscible solvent systems to maintain high concentrations of substrate/product and active catalysts in separate phases. Rapid exchange between phases allows for turnover and accumulation of product in the hydrocarbon phase. B) Example water-immiscible solvents. Solvents must be immiscible with both water and HFE7500 to be compatible with FACS sorting. Longer alkyl chains reduce solvent miscibility with water relative to more standard industrial solvents. C) Pipeline for screening biphasic reactions in FACS sortable triple emulsion picoreactors. D) Droplet microfluidics (green outline) increases throughput and reduces both cost and waste relative to traditional techniques.

Nevertheless, the optimization of biphasic reactions remains challenging. Beyond challenges common to all chemical reactions (e.g. yield, side product contamination, enantiomeric excess, etc.), optimizing biphasic reactions requires maximizing interfacial surface area over the duration of an experiment^20^ (e.g. via mixing^21^ or continuous and segmented flow in microfluidic reactors^22–24^) and identifying reaction-compatible solvents with low water-miscibility. Common hydrocarbon solvents contain few atoms and are often polar and miscible with water (e.g. fully: methanol, ethanol, acetone; partially: ethyl acetate). Solvents with the same polar functional groups and longer alkyl chains (e.g. hexyl acetate, 1-octanol, 2-octanone) have lower miscibility with water (**Supplementary Table 1**) but may retain desirable solvent properties (e.g. ketone solvation of plastic polymers). While these more hydrophobic solvents have promise for potential use within biphasic systems, experimental pipelines capable of systematically testing and optimizing their use within high-throughput screening platforms have been lacking.

Microfluidics provides a particularly promising method for high-throughput screening of biphasic reactions, as the relatively small volumes and large interfacial surface areas can dramatically enhance small molecule transfer. Reflecting this, microfluidic slug/plug flow devices have been used to screen 10– 100 biphasic reaction conditions at a time using in-line monitoring^22,25,26^(**Supplementary Table 2**). Droplet microfluidics provides a potential way to scale these screens via highthroughput compartmentalization and sorting to recover and identify catalysts or conditions favorable for product formation. To date, single-phase droplet microfluidics has been used to screen for and isolate promising candidates from libraries of variants for organic^27^, metallic^28^, organometallic^29^, and enzymatic catalysts^30–33^ via FADS (Fluorescence Activated Droplet Screening) and MADS (Mass spectrometry Activated Droplet Screening), both of which require custom device fabrication and equipment. More recently, double emulsion droplets have enabled ultra-high-throughput encapsulation, screening, and isolation of pL-volume reactions using only simple microfluidic devices and commercially-available equipment^34–37^. In double emulsion droplets, reactions of interest are encapsulated within a thin fluorocarbon oil shell such that they can be loaded into and sorted by commercially available FACS instruments at rates of up to 1–5 kHz (9 million droplets/hour). Double emulsion droplet screening platforms compatible with biphasic reactions – triple emulsions – would allow for efficient search through complex combinatorial chemical spaces for desirable catalysts (*e*.*g*. DNA-encoded small molecule catalysts or directed evolution of enzymes that can be generated via cell-free synthesis). However, the lack of methods for generating stable FACS-sortable triple emulsion picoreactors is a significant technical barrier to realizing such a high-throughput screening platform. This method development requires (1) selecting 3 mutually immiscible phases (aqueous, hydrocarbon, and fluorocarbon) (**Figure 1B, Supplementary Figure 1**), (2) optimizing surfactants to stabilize those interfaces, (3) producing triple emulsion picoreactors, (4) characterizing triple emulsion stability, and (5) confirming triple emulsion compatibility with both the desired reaction and flow cytometry/cell sorting instruments.

Here, we present the generation of novel triple emulsion picoreactors that encapsulate a biphasic reaction environment and have been optimized for high-throughput screening using FACS. Hydrocarbon/aqueous biphasic solutions are encapsulated within fluorocarbon oil shells to form triple emulsions that can be screened using commercially available FACS instruments, with applications to high-throughput screening of enzyme variants produced via cell-free protein synthesis (**Figure 1C**). To achieve this, we first developed and deployed a novel plate-based screening pipeline to identify combinations of solvents and surfactants capable of forming stable hydrocarbon/aqueous emulsions (e.g. octanol with aqueous buffer). We then loaded stable hydrocarbon/aqueous emulsions into droplet generators to yield triple emulsion picoreactors (**Supplementary Figure 1**) that remained stable over 10s of hours. We demonstrated cell-free protein synthesis within the aqueous phase of triple emulsion picoreactors, which will enable highthroughput screening of libraries of enzyme catalysts. Finally, we demonstrated the ability to sort triple emulsion picoreactors without custom equipment, which will make it possible to screen libraries of >10^6^ variants while drastically reducing the reagent volumes required (**Figure 1D**).

## RESULTS

An optimized experimental pipeline to identify hydrocarbon solvent/aqueous buffer/surfactant combinations. Successful FACS-based screening of biphasic solvent/water reactions requires: (1) that hydrocarbon-in-water droplets can be encapsulated within a fluorocarbon shell, (2) that these triple emulsions remain stable over time, and (3) that the overall dimensions of the triple emulsion are sufficiently small to maximize interfacial surface area and pass through FACS nozzles without disrupting stable water-in-air droplet breakoff (Supplementary figures 1,2). Maintaining triple emulsion stability depends critically on the use of surfactants to tune interfacial surface tensions at each interface (inner aqueous/hydrocarbon, inner aqueous/oil, and oil/outer aqueous). Consistent with this, initial naïve attempts to generate triple emulsions in the absence of aqueous/solvent surfactants destabilized oil shells, preventing successful sorting (Supplementary Figure 3). To systematically and efficiently identify promising hydrocarbon/aqueous buffer/surfactant combinations, we leveraged the fact that changes in the number and size of emulsified droplets for two fluids with different refractive indices alter light transmission. Thus, optical properties (e.g. optical density, absorbance, turbidity) can be used to monitor emulsion stabilities^38,39^. Specifically, we: (1) tested surfactant solubilities in aqueous buffer and hydrocarbon solvents, (2) vortexed mixtures of aqueous buffer, hydrocarbon solvents, and aqueous- and hydrocarbon-compatible surfactants to emulsify them, and then (3) assessed droplet formation and stability via plate-based light transmission assays and microscopy; after identifying promising combinations, we performed an additional screen to optimize surfactant concentrations (Figure 2A). While we focus here on octanol because of its compatibility with fluorinated oils required to create FACS-sortable microfluidic droplets, this general approach could be used to optimize reaction conditions for a wide range of alternative screens.

As a first step in screening for promising fluid/surfactant combinations, we assessed the solubility of 15 readily available commercial surfactants in a standard aqueous buffer (phosphate buffered saline (PBS)) pH 7.4, a model enzyme reaction buffer) and four 8-carbon solvents (octanol, octane, 2-octanone, and hexyl acetate) (**Figure 2B, Supplementary Table 1**). All 4 solvents are known to form a biphasic system with water, with reported solubilities in water of 0.00066 g/L for octane and 0.3– 0.9 g/L for the remaining solvents. The 15 surfactants included 9 non-ionic surfactants, 2 cationic surfactants, 2 anionic surfactants, and 2 zwitterionic surfactants spanning a broad range of hydrophilic-lipophilic balance (HLB) values (see **Methods** for further discussion). Nearly all ionic surfactants (CHAPS, SB3-10, Sarkosyl, SDS) were soluble only in PBS pH 7.4, except for benzalkonium chloride (which was also soluble in 1-octanol and 2-octanone); two surfactants were provided as aqueous solutions (NP-10, CTAB), precluding solvent solubility tests. While surfactants with the lowest HLB values (e.g. EM-90, SPAN) were soluble only in organic solvents (**Supplementary Table 3**), other non-ionic surfactants were generally soluble in PBS pH 7.4 (e.g. Triton X-100, Tween-80, and Tween-20). While solubility generally trended with surfactant HLB values and solvent miscibility values (**Figure 2B, Supplementary Table 3, Supplementary Figure 4**), the relationship was not fully predictive (e.g. for Triton CG110 and for Span 20), establishing a need for direct empirical testing.

**Figure 2:**
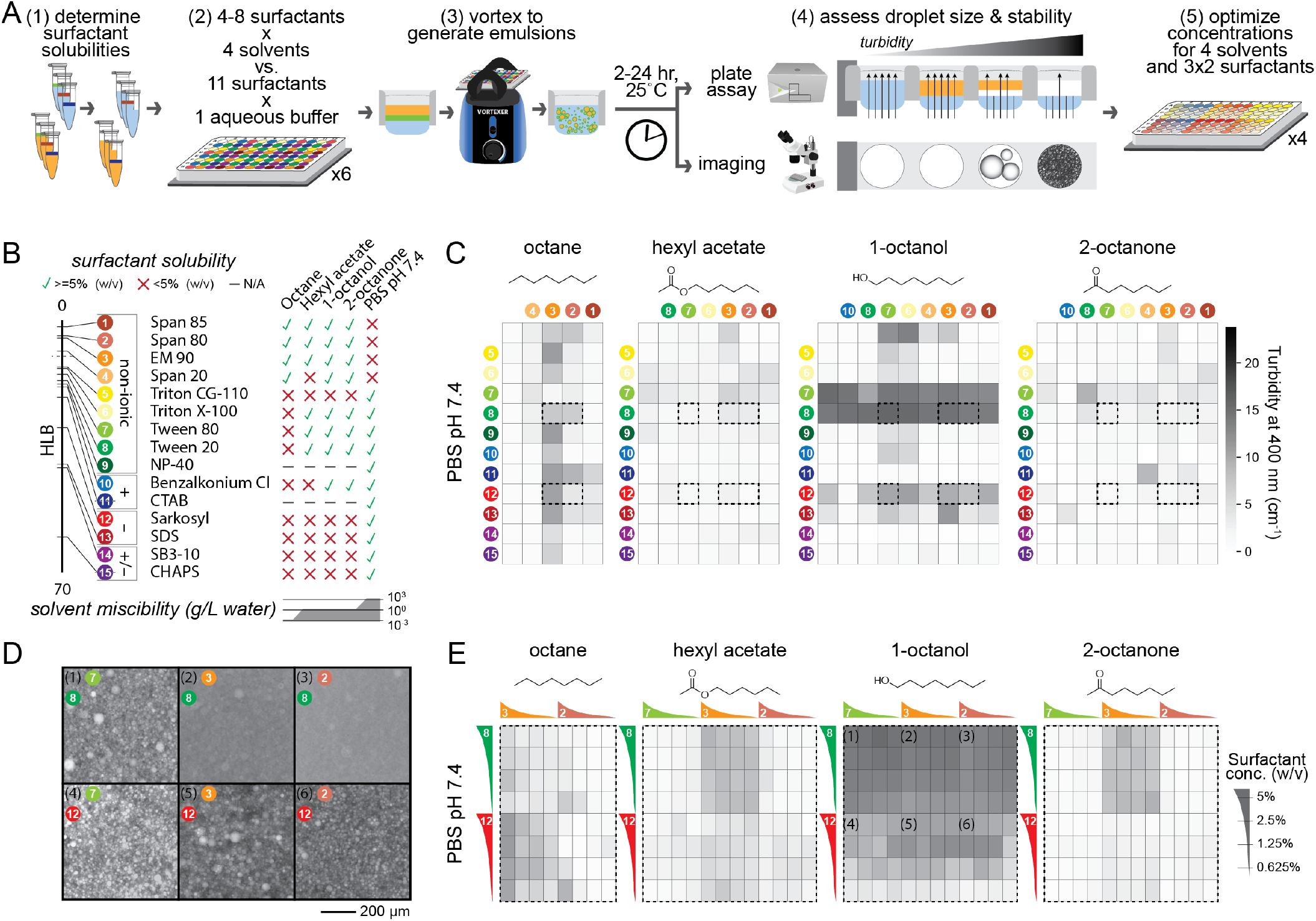
Turbidityand microscopy-based screen identifies surfactants for stabilizing hydrocarbon/aqueous single emulsions. **A)** Workflow for rapid assay to screen optimal surfactant conditions by (1) testing if surfactants are soluble in aqueous buffer and/or hydrocarbon solvents, (2) vortexing and screening for droplet formation, (3) validating and characterizing candidate droplet-forming combinations by microscopy, and (4) optimizing concentrations for droplet size and stability. **B)** Solubility for selected surfactants arranged by hydrophiliclipophilic balance (HLB) and categorized by charge as non-ionic, cationic (+), anionic (), or zwitterionic (+/−). Green checks and red Xs indicate solubility at > or < 5% (w/v), respectively. **C)** Plate reader turbidity measurements after 24 hours for one aqueous buffer (PBS pH 7.4) and 4 hydrocarbon solvents with surfactant combinations. Dashed black outlines indicate selected conditions used in D and E. **D)** Fluorescence microscopy images of octanol/aqueous emulsions stabilized by the selected surfactants from C); octanol is fluorescently labeled with Nile Red. Scale bar: 200 µm. **E)** Turbidity measurements as in C from screening selected surfactants over concentrations from 0.625-5% (w/v).

### Plate-based turbidity assay can screen for surfactants that stabilize hydrocarbon/aqueous emulsions

Next, we tested all possible combinations of aqueous buffer (1), hydrocarbon solvent (4), aqueous-soluble surfactant (8), and hydrocarbon-soluble surfactants (11) for their ability to form emulsions (8 plates in total) by including each surfactant at a single high concentration (5% w/v) and quantifying emulsion stability at 2 hrs and 24 hrs via a plate-based turbidity screen and microscopy (see **Supporting Information**). Plate-based screening revealed differences in turbidities as a function of surfactant conditions and over time, providing measurements of droplet formation and stability (**Figure 2C, Supplementary Figures 5-11**). Fine emulsions formed for each solvent, with some wells appearing visibly white and opaque (in contrast to the optically clear appearance of conditions without surfactant). After 2 hrs, maximum turbidities for each solvent ranged from 10.6–19.3 cm^-1^, consistent with optically dense solutions of fine emulsions; a higher fraction of octane conditions formed fine, optically dense emulsions. After 24 hrs, median turbidity values dropped but maximum turbidities remained high, consistent with a general demulsification across conditions but with the finest emulsions being most stable. As expected, turbidity values at 24 hrs negatively correlated with mean intensities of microscopy images (**Supplementary Figure 12**); hexyl acetate and 2-octanone emulsions clustered at well edges (**Supplementary Figure 9**,**11**) but octanol emulsions appeared well-dispersed (**Supplementary Figure 10**). Overall, 24 surfactant conditions with octanol were somewhat stable (turbidity >9.8 (50% of maximum) after 24 hrs); 11 conditions were highly stable (turbidity >17.6 (90% of maximum)) and frequently containing Tween, Span, and EM90 surfactants. To more carefully examine impacts of surfactants on emulsion formation and stability, we regenerated octanol/aqueous emulsions for a subset of highly stable conditions (Tween 80, EM90, and Span 80 hydrocarbon solvent surfactants; Tween 20 and Sarkosyl aqueous surfactants) and imaged the resulting emulsions via microscopy (**Figure 2D, Supplementary Figure 13**). Droplets generated with Tween 20 had smaller oil droplet radii (median=1.0–1.75 µm) than those generated with Sarkosyl (median=2.7–4.5) (Supplementary Figure 14), confirming that Tween 20 and Sarkosyl stabilize octanol/aqueous emulsions. Tween 20 yields slightly finer emulsions, consistent with higher measured turbidities.

To assess the concentration-dependence of surfactant stabilization, we repeated screens with surfactant concentrations from 0.625–5% (**Figure 2E, Supplementary Figures 15-21**). Droplet stabilities (turbidity values) generally increased with increasing Sarkosyl in the aqueous phase regardless of octanol surfactant concentrations while Tween-20 impacts were more concentration-independent (**Supplementary Figure 16**). These results establish that increasing surfactant concentration does not always increase emulsion stability and again highlight a need for direct experimental screening. Ultimately, we identified many surfactant conditions that can stabilize micronscale hydrocarbon/aqueous emulsions for 10s of hours, providing a starting point for further optimization to specific assays and applications.

### Triple emulsion picoreactors can be formed by encapsulating octanol/aqueous emulsions in fluorocarbon oil shells

After identifying multiple reagent combinations capable of forming stable octanol/aqueous emulsions, we tested whether emulsions containing these surfactants could be successfully and stably encapsulated within an oil shell to yield FACS-sortable triple emulsions (**Figure 3A**). Specifically, we: (1) labeled octanol with Nile Red, a lipophilic dye that preferentially partitions into octanol rather than aqueous buffer or fluorinated oil (**Supplementary Figure 22**), (2) generated octanol/aqueous droplets with relatively small median radii via vortex emulsification using Tween-20 and Span-80 as the aqueous and octanol surfactants, respectively (**Figure 2D, Supplementary Figure 13**), (3) introduced this octanol/aqueous emulsion as the inner aqueous phase within a double emulsion droplet generator (**Figure 3B**), and then (4) collected and imaged the resultant output droplets via brightfield and fluorescence microscopy (see **Supporting Information**). The octanol/aqueous emulsion appeared opaque within the droplet generator inlets and yielded droplets comprised of an oil shell and an opaque core, many of which contained one or more smaller inner droplets. As expected, the lipophilic Nile Red dye colocalized with the small droplets within the inner core, confirming successful formation of FACSsortable triple emulsion droplets (**Figure 3C,D**). Overall, nearly 50% of output droplets displayed the desired triple emulsion architecture (44.5±2.5%; n=1087), with a mean triple emulsion radius of 32.0±4.1 µm (**Figure 3 E, Supplementary Figures 23**,**24**). Triple emulsions were stable over 10s of hours, with 63.3±10.5% of droplets surviving to 24 hrs (**Figure 3E**).

**Figure 3:**
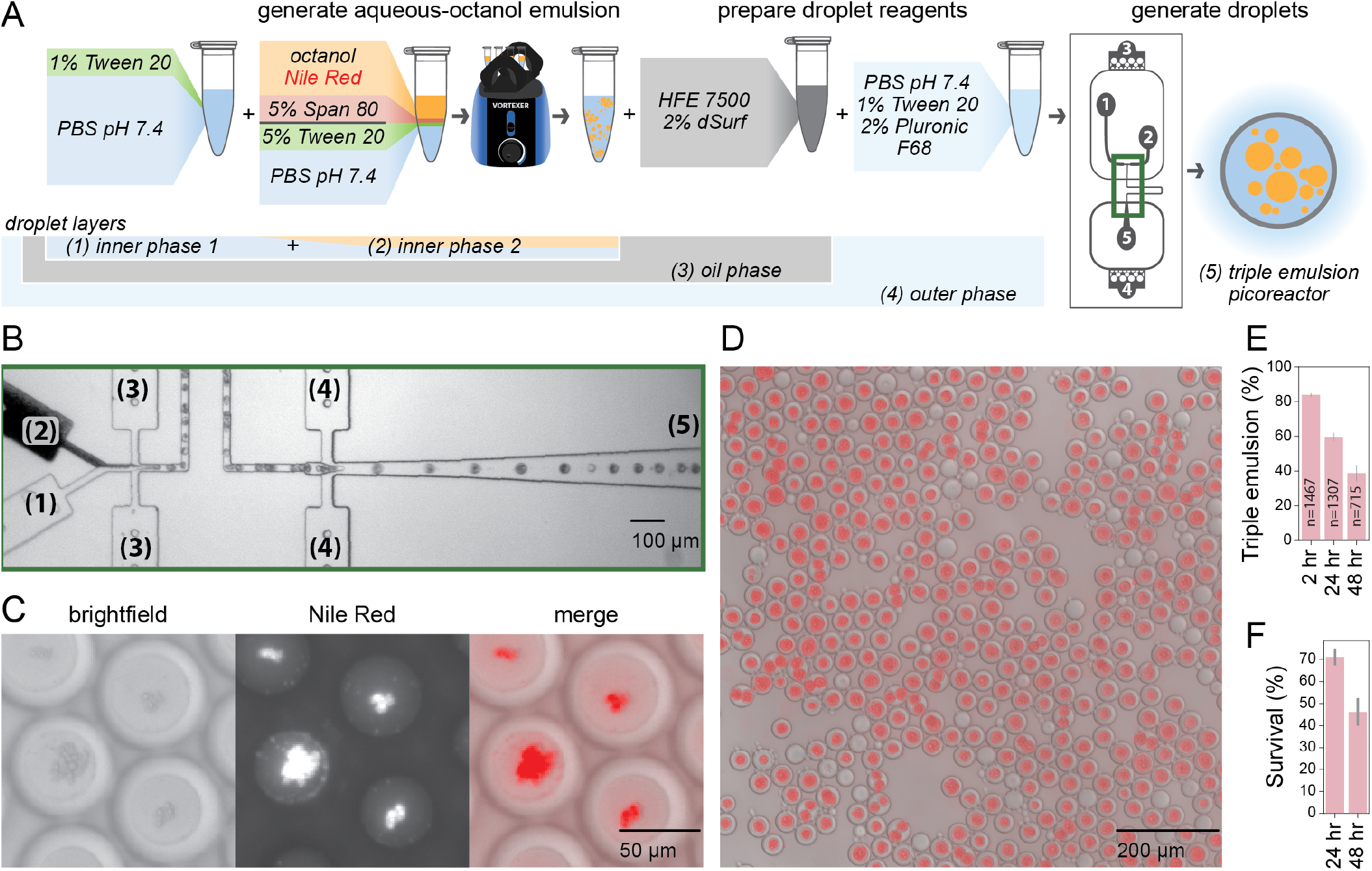
Encapsulating hydrocarbon/aqueous single emulsions in double emulsions produces triple emulsion picoreactors. **A)** Workflow for generating triple emulsions by encapsulating pre-emulsified octanol/aqueous emulsions in the core of aqueous/fluorocarbon/aqueous double emulsions. Numbered inputs and outputs match the ports/channel on the droplet generator schematic. **B)** Microscopy images of droplet generation at the region of the droplet generator outlined in green in A. Channels numbered as in A. Scale bar: 100 µm. **C)** Brightfield and fluorescence microscopy images of triple emulsions; octanol is fluorescently labeled with Nile Red. **D)** Merged brightfield and fluorescence image of produced triple emulsion droplets. **E)** Percentage of triple emulsions within produced droplet population over time. Error bars represent standard deviation in percentages across 3 images. Number of total droplets indicated on each bar. F) Survival rates reported as the ratio of triple emulsion percentages at 24 hrs versus at 2 hrs.

### *Proteins can be expressed* in situ *within triple emulsion picoreactors*

Enzymes catalysts are particularly amenable to high-throughput screening strategies when produced via *in vitro* cell-free protein synthesis within an aqueous phase^40,41^. To test whether our triple emulsion picoreactors can be used to express and screen enzyme catalysts, we first tested compatibility of in vitro transcription/translation (IVTT) with reagents required to form and stabilize droplets (**Figure 4A**). To identify solvents compatible with IVTT, we combined a plasmid encoding expression of GFP with PURE (Protein expression Using Recombinant Elements) reagents and hydrocarbon solvents at a 1:1 ratio, incubated to allow for protein expression, and then quantified GFP fluorescence (**Figure 4B**). All four 8-carbon solvents supported expression (1-octanol, 2-octanone, hexyl acetate, and octane), with GFP signal negatively correlated with solvent miscibility in water (**Figures 4B-C, Supplementary Table 2**). Next, we investigated which water-soluble surfactants were compatible with IVTT by quantifying fluorescence intensity after expressing GFP in PURE reactions containing 0.625–5% of water-soluble surfactants (**Figure 4D**). While non-ionic surfactants (Tween, Triton) had no impact on GFP expression, cationic and anionic surfactants (benzalkonium chloride, sarkosyl) completely ablated expression; the zwitterionic surfactant CHAPS reduced GFP signal in a concentrationdependent manner (**Figure 4D**). Next, we tested for the ability to successfully produce proteins within triple emulsions for high-throughput screening by: (1) generating triple emulsion droplets with inner cores comprised of octanol/aqueous droplets containing Nile Red dye, Tween 20 and Span 80 surfactants, plasmid DNA encoding GFP expression, and all reagents required for IVTT, (2) incubating to allow IVTT and GFP production, and then (3) imaging to quantify the amount of expressed GFP in each droplet (**Figure 4E, Supporting Information**). To eliminate the possibility of GFP expression prior to droplet formation, inner core reagents were introduced separately via 2 inputs such the plasmid DNA and IVTT reagents did not contact each other until just before droplet formation within the device. Droplets incubated at 37°C for 2 hrs showed strong fluorescence in the green channel while negative control droplets incubated at 4°C (a temperature below that required for IVTT) did not fluoresce (**Figure 4F**). Varying flow rates for the plasmid-octanol solution and the IVTT reagents (and thereby changing the relative volume fractions of each solution within the inner core) led to concomitant changes in Nile Red and GFP intensities, consistent with either IVTT reagents being the limiting factor for expression in this experiment or an inhibitory impact of octanol when present at very high surface areas (**Figure 4G**). For the 100:100 flow ratio condition, the output droplets consisted of 94.0±1.5% (n=1525) triple emulsions with a mean triple emulsion radius of 35.0±4.8 µm (**Figure 4H, Supplementary Figure 25**). 75.3±3.8% of triple emulsions incubated for 2 hrs at 37°C survived to 24 hrs at room temperature and 61.5±6.1% survived to 48 hrs (**Figure 4I**). These results indicate that biocatalysis can be performed in triple emulsion picoreactors, including for challenging reactions that require 10s of hours.

**Figure 4:**
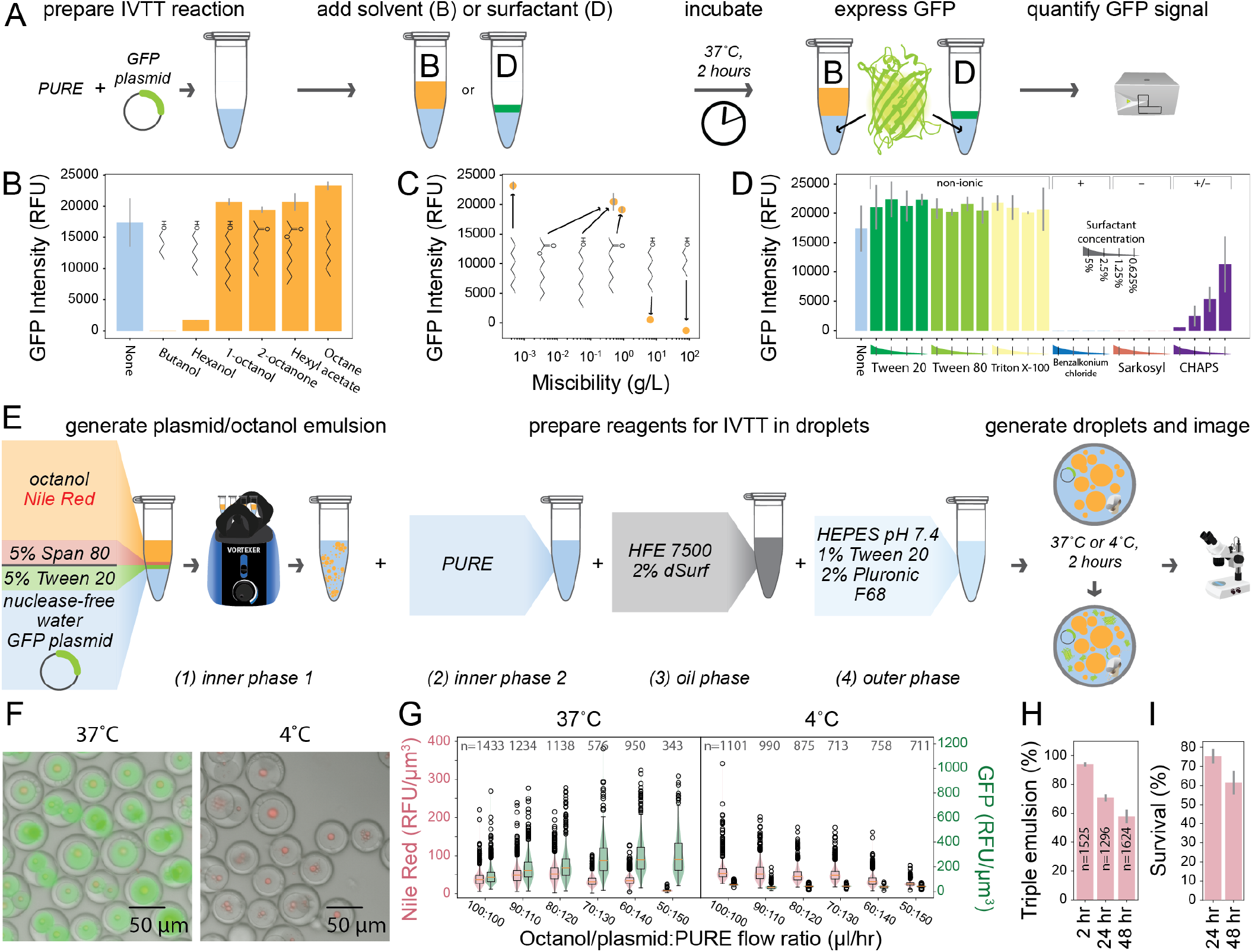
in vitro *transcription translation yields expressed protein within triple emulsion picoreactors*. **A)** Workflow for testing IVTT compatibility with aqueous-hydrocarbon emulsions. GFP expression with PURE reagents in the presence of solvent **(B)** or surfactant (D) quantifies transcription/translation activity in solution. B) GFP signal for IVTT reaction in the presence of equal volumes of hydrocarbon solvent. Error bar represents standard deviation from measurements of 3 reactions. **C)** GFP signal from B plotted against solvent miscibility in water. **D)** GFP signal for IVTT reaction with 0.625–5% water-soluble surfactant. Error bar represents standard deviation from measurements of 3 reactions. Labels indicate surfactant charge. **E)** Workflow for expressing GFP in triple emulsion picoreactors. Protein expression is performed at 37°C (positive control) or 4°C (negative control); octanol is fluorescently labeled with Nile Red. **F)** Merged images of triple emulsion picoreactors with IVTT reagents incubated at 37°C and 4°C showing brightfield, GFP fluorescence (produced protein), and Nile Red fluorescence (octanol droplets). **G)** Nile Red and GFP fluorescence intensities across various flow rate ratios for octanol/plasmid + PURE reaction mixture. **H)** Percent triple emulsions within the output droplet population. Error bars represent the standard deviation across 3 images. I) Survival rates reported as the ratio of triple emulsion percentages at 24 and 48 hrs versus at 2 hrs.

### Desired picoreactor populations can be selected and recovered via FACS sorting

Next, we tested if our triple emulsion picoreactors could be analyzed and sorted via FACS. To increase the recovery of sorted droplets, we reduced the size of the droplets by scaling the dimensions of the microfluidic devices to 2/3. With these smaller droplet generators, we used the optimized surfactants described above to produce triple emulsions containing IVTT reagents and Nile Red-labeled octanol and double emulsions that contained no fluorophore (**Figure 5A-C**). The triple emulsion output contained 90.9±1.8% (n=1154) triple emulsions with a mean triple emulsion radius of 26.4±2.6 µm, and the double emulsion output contained >99.0±0.0% (n=506) double emulsions with a mean double emulsion radius of 29.4±1.9 µm (**Supplementary Figure 26-27**). We incubated the triple emulsions at 4°C (to inhibit GFP expression) or 37°C (to promote GFP expression) for 2 hrs and sorted the separate droplet populations by FACS. Event rates during sorting varied between 100-800 Hz. Double and triple emulsion populations were clearly visible on plots of FSC-A vs. SSC-A (**Figure 5A-C**). As expected, double emulsions showed low signal for Nile Red (indicating an absence of octanol) and both triple emulsion samples showed increased and comparable Nile Red signal, confirming that triple emulsions can be reliably identified by fluorescence (**Figure 5D**). Again as expected, signal intensity was higher for triple emulsions incubated at 37°C (median = 1595.9) than for triple emulsions incubated at 4°C (median = 390.3) and double emulsions (median = 76.2) (**Figure 5E**). After sorting with gates selecting for droplets with the highest GFP signal (**Supplementary Figure 28, Supplementary Table 4**), recovered droplets contained 96.0±3.1% (n=703) triple emulsions and were enriched for high fluorescence droplets (**Figure 5F-G, Supplementary Figure 26**). 75-80% of sorted droplets were recovered (see **Supporting Information**). These results confirm that triple emulsion picoreactors can be sorted by fluorescence and recovered for downstream assays.

**Figure 5:**
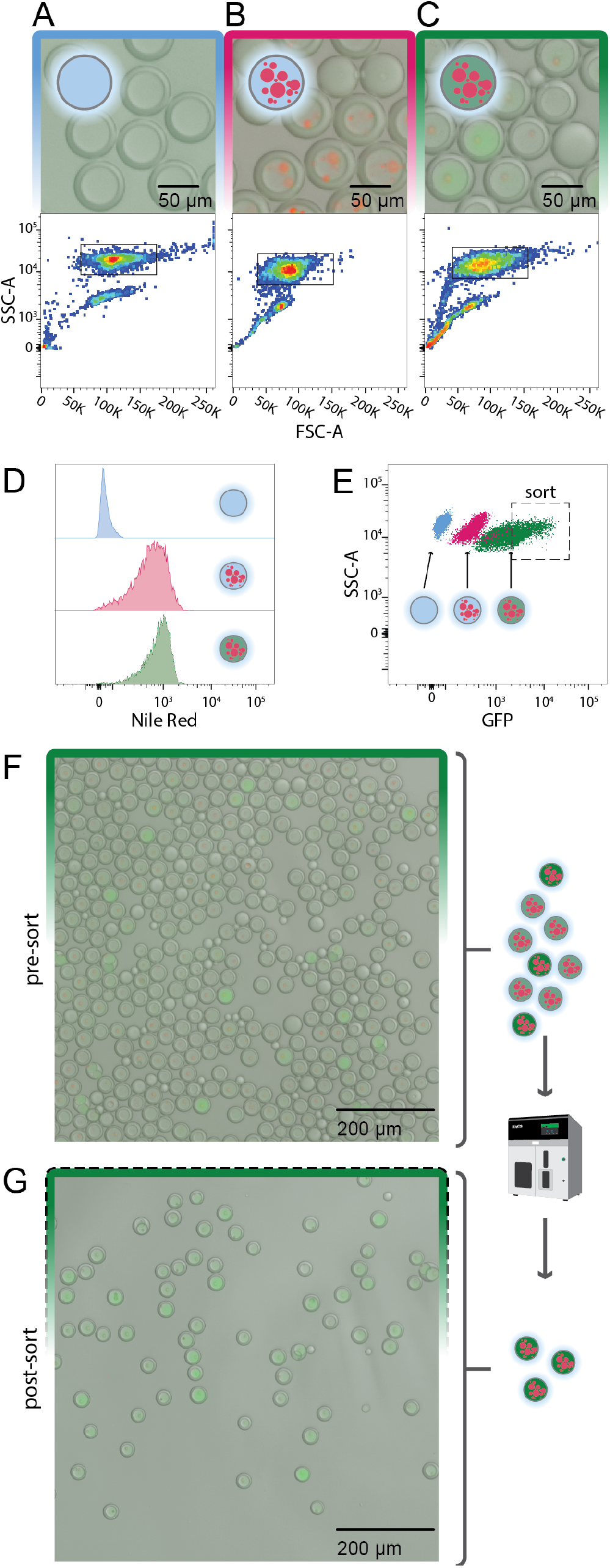
FACS sorting recovers target picoreactor subpopulation. Pre-sort images (top) and side scatter *vs*. forward scatter FACS distributions (bottom) for A) double emulsions, **B)** triple emulsion picorectors incubated at 4°C to prevent protein expression. or **C)** incubated at 37°C to promote protein expression. Black outline indicates FACS droplet quality gates. **D)** Histograms of Nile Red FACS signals for each sample. **E)** GFP FACS signals *vs*. side scatter for each sample. Dashed black outline indicates FACS sorting gates. **F)** Images of triple emulsion picoreactors before and **G)** after sorting.

## DISCUSSION

For new catalysts in general, much attention has been directed to developing large-scale screens for evaluating catalysts for aqueous reactions^28^ and using machine learning algorithms to predict likely candidates from prior data^42,43^. Thus, there is a natural synergy between high-throughput screening methods and state-of-the-art data analysis. Droplet microfluidic technologies have great promise for screening complex reaction conditions where the optimization space is large such as combinatorial condition space of biphasic reactions versus the chemical space of small molecule catalysts or the sequence space of enzymes using bead-display libraries.

Here, we demonstrated the generation of triple emulsion picoreactors compatible with FACS sorting. The selection of optimal surfactants was aided by a rapid plate-based screening strategy that identified surfactants compatible with *in situ* expression of protein candidates for enzyme screening. As one example, these triple emulsions with an octanol-in-water core could be used in future work to engineer lipases that generate biodiesel. This optimization approach is extensible to other organic solvents that are not immediately compatible with microfluidic droplets or are compatible with other droplet-based screening platforms (such as FADS).

Compared to other microfluidic methods, this method uses simple, robust microfluidic devices and commercial sorting machines, enabling screening at very high throughput using minimal amounts of reagents. Even at conservative sorter analysis rates of 100–200 Hz, 1 million droplet reactions can be screened in 2-3 hours, making it possible to screen 10s of millions of reactions per day. Moreover, the small internal volumes demonstrated here (∼20 pL) make it possible to screen 10^6^ reactions using 20 µL of reagents. Assuming reactions of 5 µL for each phase in 1536-well plates, this results in a 500,000-fold reduction in reagents from 10 L of reagents over 10^6^ reactions in >650 plates. Compared to benchtop scale experiments where the less dense hydrocarbon phase has the tendency to separate or cream, encapsulating biphasic reactions within a droplet picoreactor ensures that the hydrocarbon phase is well dispersed within the aqueous phase without the need for mixing.

Further technological developments may improve the robustness and applicability of triple emulsion picoreactors to diverse applications. For example, redesigning the droplet generator to form octanol/aqueous droplets on-chip upstream of encapsulation could eliminate the need for the pre-emulsification step and result in more regular droplet geometries. Similarly, the addition of multiple inlet channels could enable more complex onchip mixing, facilitating condition screening. Polymerizable or mineralizable shells may expand the range of compatible organic solvents beyond those that are insoluble in HFE7500 and other fluorocarbon oils^44,45^. In the future, FACS sortable triple emulsion picoreactors could be used to test assay conditions with industrially relevant non-aqueous phases such as petroleum products^10,11,46^, pyrolysis oil from thermocracked plastics^47–49^, and recycled cooking oils^4,50^. In biphasic reactions with these hydrocarbon phases, catalysts – either small molecule or enzymatic – could be screened from DNA-encoded libraries where one bead displays many copies of a small molecule or gene. With the scale possible from such a platform, the data generated would both complement and enable robust learning algorithms for catalyst design.

## METHODS

Extended methods are available in **Supporting Information**.

### Accessible data and materials

Plasmid constructs are deposited in Addgene under accession number 216849. All python scripts used for data and image analysis are available at https://osf.io/gbq5r/ with all inputs used in analysis.

### Plate reader screening

Solutions of hydrocarbon solvents (Sigma-Aldrich) and PBS pH 7.4 (Gibco) were pre-mixed with surfactants at concentrations of 0.625-5% (w/v). 25 µL of hydrocarbon solution was added to 75 µL of aqueous solution in wells of a 96-well plate (Nunc). Plates were vortexed before reading optical absorbance in a multi-mode plate reader (Tecan) at 2 hrs and 24 hrs. At 24 hrs, all wells were imaged with an optical microscope (AmScope). Turbidity values were calculated from absorbance of 400 nm light using custom python scripts.

### Droplet generation

Droplets were generated of custom fabricated PDMS devices using previously reported designs^34^. Input solutions described in **Figures 3** and **4** were loaded into plastic syringes (BD) and connected to the PDMS device via LDPE medical tubing (Scientific Commodities Inc.). Syringe pumps drove the flow of reagents into the device, with standard flow rates of 100 µL/hr for each inner solution, 400 µL/hr for the fluorocarbon oil, and 4000 µL/hr for the outer aqueous phase. For triple emulsions with PBS as the aqueous phase, inner solutions were generated from hydrocarbon and aqueous solutions used in the plate reader screen. Octanol was labeled with 400 µM Nile Red (Sigma Aldrich). For protein expression in droplets, the two inner solutions were comprised of 1) PURExpress reagents (NEB) or 2) emulsions of octanol + 5% (w/v) Span 80 + 40 µM Nile Red in an aqueous phase of nuclease free water (Promega) + 200 ng/µL miniprepped plasmid + 5% (w/v) Tween-20.

### Microscopy

Droplet emulsions were images with bright field and fluorescence microscopy (Nikon) with green and red fluorescence filter cubes. Flatfield correction, particle detection, and fluorescence quantification were performed using custom python scripts and the cv2/OpenCV library.

### FACS

Droplets were sorted on a FACSAria II cell sorter (BD) with a 130 µm nozzle. Laser delays were manually calibrated with 32 µm AccuCount Ultra Rainbow beads (Spherotech). Forward and side scattering voltage were manually optimized samples of double emulsions. Droplet delays were manually calibrated by sorting 50 double emulsions onto a glass slide at each setting and manually counting the recovered droplets by optical microscopy. Detector voltages were optimized using triple emulsion samples with expressed GFP and Nile Red labeled octanol.

## ASSOCIATED CONTENT

### Supporting Information

A Supporting Information document (.pdf) including supplementary figures, tables, and methods is available free of charge on the ACS Publications website. Plate reader turbidity measurements (.csv) are included as Supplementary Data 1-8. Image (.tif) and .csv files for octanol/water emulsions are included as Supplementary Data 9-10. Images (.tif), .csv files, and .fcs for the generation of triple emulsions and their analysis by microscopy and FACS are included as Supplementary Data 11-17.

## AUTHOR INFORMATION

## Author Contributions

SMT and PMF conceived the project, provided funding, and wrote the manuscript. SMT produced materials, emulsions, and droplets; collected plate reader and image data; and analyzed all data. SMT, PMF, and YZ prepared figures. SMT, YZ, ZY, and LAN performed FACS sorting. All authors have given approval to the final version of the manuscript. ‡These authors contributed equally.

### NOTES

The authors declare no following competing financial interests.

## ABBREVIATIONS

FACS: fluorescence activated cell sorting
FADS: fluorescence activated droplet sorting
FSC-A: forward scatter area
GFP: green fluorescent protein
HLB: hydrophilic lipophilic balance
IVTT: *in vitro* transcription/translation
MADS: mass activated droplet sorting
PBS: phosphate buffered saline
PDMS: polydimethylsiloxane
PURE: protein synthesis using recombinant elements
SSC-A: side scatter area

## ACKNOWLEDGMENT

The authors would like to thank the members of the Fordyce lab for helpful discussion and insightful feedback. The SMT000 plasmid was a gift from Akshay Maheshwari and the Endy lab. This work was supported by a Garden Grant from the Homeworld Collective. Data was collected on an instrument in the Shared FACS Facility obtained using NIH S10 Shared Instrument Grants S10RR027431-01 and S10RR025518-01. SMT is supported by the Arnold and Mabel Beckman Foundation through the Arnold O. Beckman Postdoctoral Fellowship in Chemical Sciences. ZY was supported by the Sanjiv Sam Gambhir - Philips Fellowship in Precision Health Program in the Department of Radiology of Stanford University. PMF is a Chan Zuckerberg Biohub Investigator.

## Table of Contents

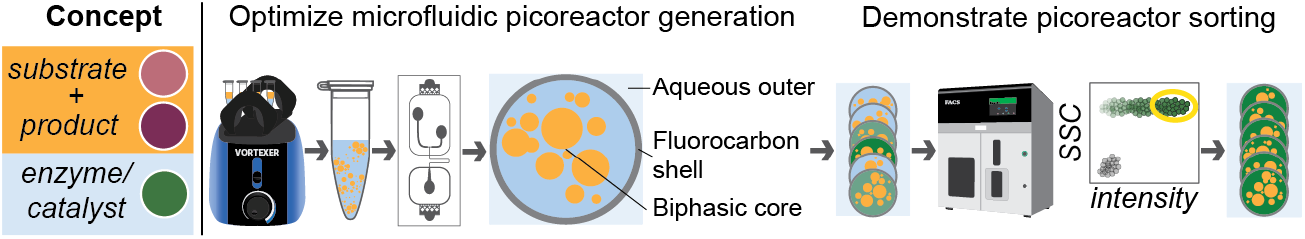

Biphasic reactions can enhance reactions where catalyst and substrate have different solvent preferences. To develop a high-throughput platform for screening biphasic reaction conditions, a method was optimized for producing triple emulsion picoreactors that encapsulate a biphasic solvent environment. These triple emulsion picoreactors can be sorted in commercially available cell sorters for future highthroughput screens.

## REFERENCES

(1) Nasibullin, I.; Yoshioka, H.; Mukaimine, A.; Nakamura, A.; Kusakari, Y.; Chang, T.-C.; Tanaka, K. Catalytic Olefin Metathesis in Blood. Chem. Sci. 2023, 14 (40), 11033–11039. 10.1039/D3SC03785A.

(2) Mitchell, S.; Qin, R.; Zheng, N.; Pérez-Ramírez, J. Nanoscale Engineering of Catalytic Materials for Sustainable Technologies. Nat. Nanotechnol. 2021, 16 (2), 129–139. 10.1038/s41565-020-00799-8.

(3) Hai, X.; Zheng, Y.; Yu, Q.; Guo, N.; Xi, S.; Zhao, X.; Mitchell, S.; Luo, X.; Tulus, V.; Wang, M.; Sheng, X.; Ren, L.; Long, X.; Li, J.; He, P.; Lin, H.; Cui, Y.; Peng, X.; Shi, J.; Wu, J.; Zhang, C.; Zou, R.; Guillén-Gosálbez, G.; Pérez-Ramírez, J.; Koh, M. J.; Zhu, Y.; Li, J.; Lu, J. Geminal-Atom Catalysis for Cross-Coupling. Nature 2023, 622 (7984), 754–760. 10.1038/s41586-023-06529-z.

(4) Korman, T. P.; Sahachartsiri, B.; Charbonneau, D. M.; Huang, G. L.; Beauregard, M.; Bowie, J. U. Dieselzymes: Development of a Stable and Methanol Tolerant Lipase for Biodiesel Production by Directed Evolution. Biotechnol. Biofuels 2013, 6 (1), 70. 10.1186/1754-6834-6-70.

(5) Tournier, V.; Topham, C. M.; Gilles, A.; David, B.; Folgoas, C.; Moya-Leclair, E.; Kamionka, E.; Desrousseaux, M.-L.; Texier, H.; Gavalda, S.; Cot, M.; Guémard, E.; Dalibey, M.; Nomme, J.; Cioci, G.; Barbe, S.; Chateau, M.; André, I.; Duquesne, S.; Marty, A. An Engineered PET Depolymerase to Break down and Recycle Plastic Bottles. Nature 2020, 580 (7802), 216–219. 10.1038/s41586-020-2149-4.

(6) Zhong, W.; Mao, L.; Yi, W.; Zou, G.; Li, Y.; Yin, D. Highly Efficient Light-Driven HNO3 Nitration–Oxidation of Cyclohexane to Co-Product Nitrocyclohexane and Adipic Acid in a Biphasic System. Res. Chem. Intermed. 2016, 42 (2), 461–470. 10.1007/s11164-015-2030-5.

(7) Schrimpf, M.; Graefe, P. A.; Holl, A.; Vorholt, A. J.; Leitner, W. Effect of Liquid–Liquid Interfacial Area on Biphasic Catalysis Exemplified by Hydroformylation. ACS Catal. 2022, 12 (13), 7850–7861. 10.1021/acscatal.2c01972.

(8) Schowner, R.; Elser, I.; Toth, F.; Robe, E.; Frey, W.; Buchmeiser, M. R. Mono- and Bisionic Mo- and W-Based Schrock Catalysts for Biphasic Olefin Metathesis Reactions in Ionic Liquids. Chem. – Eur. J. 2018, 24 (50), 13336–13347. 10.1002/chem.201802472.

(9) Sato, Y.; Liu, J.; Ndukwe, I. E.; Elipe, M. V. S.; Griffin, D. J.; Murray, J. I.; Hein, J. E. Liquid/Liquid Heterogeneous Reaction Monitoring: Insights into Biphasic Suzuki-Miyaura Cross-Coupling. Chem Catal. 2023, 3 (7). 10.1016/j.checat.2023.100687.

(10) Stevenson, J.-A.; Westlake, A. C. G.; Whittock, C.; Wong, L.-L. The Catalytic Oxidation of Linear and Branched Alkanes by Cytochrome P450cam. J. Am. Chem. Soc. 1996, 118 (50), 12846–12847. 10.1021/ja963087q.

(11) Kellner, D. G.; Maves, S. A.; Sligar, S. G. Engineering Cytochrome P450s for Bioremediation. Curr. Opin. Biotechnol. 1997, 8 (3), 274–278. 10.1016/S0958-1669(97)80003-1.

(12) Nabgan, W.; Nabgan, B.; Tuan Abdullah, T. A.; Ikram, M.; Jadhav, A. H.; Jalil, A. A.; Ali, M. W. Highly Active Biphasic Anatase-Rutile Ni-Pd/TNPs Nanocatalyst for the Reforming and Cracking Reactions of Microplastic Waste Dissolved in Phenol. ACS Omega 2022, 7 (4), 3324–3340. 10.1021/acsomega.1c05488.

(13) Murali, V.; Kim, J. R.; Park, Y.-K.; Ha, J.-M.; Jae, J. Water-Assisted Single-Step Catalytic Hydrodeoxygenation of Polyethylene Terephthalate into Gasoline- and Jet Fuel-Range Cycloalkanes over Supported Ru Catalysts in a Biphasic System. Green Chem. 2023, 25 (21), 8570–8583. 10.1039/D3GC02145A.

(14) Wen, F.; Nair, N. U.; Zhao, H. Protein Engineering in Designing Tailored Enzymes and Microorganisms for Biofuels Production. Curr. Opin. Biotechnol. 2009, 20 (4), 412–419. 10.1016/j.cop-bio.2009.07.001.

(15) Muranaka, Y.; Matsubara, K.; Maki, T.; Asano, S.; Nakagawa, H.; Mae, K. 5-Hydroxymethylfurfural Synthesis from Monosaccharides by a Biphasic Reaction–Extraction System Using a Microreactor and Extractor. ACS Omega 2020, 5 (16), 9384–9390. 10.1021/acsomega.0c00399.

(16) Zhang, C.; Xu, Q.; Hou, H.; Wu, J.; Zheng, Z.; Ouyang, J. Efficient Biosynthesis of Cinnamyl Alcohol by Engineered Escherichia Coli Overexpressing Carboxylic Acid Reductase in a Biphasic System. Microb. Cell Factories 2020, 19 (1), 163. 10.1186/s12934-020-01419-9.

(17) Ferreira, R. da G.; Azzoni, A. R.; Freitas, S. Techno-Economic Analysis of the Industrial Production of a Low-Cost Enzyme Using E. Coli: The Case of Recombinant β-Glucosidase. Biotechnol. Biofuels 2018, 11 (1), 81. 10.1186/s13068-018-1077-0.

(18) de Castro, A. M.; Carvalho, D. F.; Freire, D. M. G.; Castilho, L. dos R. Economic Analysis of the Production of Amylases and Other Hydrolases by Aspergillus Awamori in Solid-State Fermentation of Babassu Cake. Enzyme Res. 2010, 2010, 576872. 10.4061/2010/576872.

(19) Klein-Marcuschamer, D.; Oleskowicz-Popiel, P.; Simmons, B. A.; Blanch, H. W. The Challenge of Enzyme Cost in the Production of Lignocellulosic Biofuels. Biotechnol. Bioeng. 2012, 109 (4), 1083–1087. 10.1002/bit.24370.

(20) Piradashvili, K.; Alexandrino, E. M.; Wurm, F. R.; Landfester, K. Reactions and Polymerizations at the Liquid–Liquid Interface. Chem. Rev. 2016, 116 (4), 2141–2169. 10.1021/acs.chemrev.5b00567.

(21) Starks, C. M. Interfacial Area Generation in Two-Phase Systems and Its Effect on Kinetics of Phase Transfer Catalyzed Reactions. Tetrahedron 1999, 55 (20), 6261–6274. 10.1016/S0040-4020(99)00203-3.

(22) Jovanović, J.; Rebrov, E. V.; Nijhuis, T. A. (Xander); Hessel, V.; Schouten, J. C. Phase-Transfer Catalysis in Segmented Flow in a Microchannel: Fluidic Control of Selectivity and Productivity. Ind. Eng. Chem. Res. 2010, 49 (6), 2681–2687. 10.1021/ie9017918.

(23) Nieves-Remacha, M. J.; Kulkarni, A. A.; Jensen, K. F. Hydrodynamics of Liquid–Liquid Dispersion in an Advanced-Flow Reactor. Ind. Eng. Chem. Res. 2012, 51 (50), 16251–16262. 10.1021/ie301821k.

(24) von Keutz, T.; Cantillo, D.; Kappe, C. O. Enhanced Mixing of Biphasic Liquid-Liquid Systems for the Synthesis of Gem-Dihalocyclopropanes Using Packed Bed Reactors. J. Flow Chem. 2019, 9 (1), 27–34. 10.1007/s41981-018-0026-1.

(25) Volk, A. A.; Epps, R. W.; Yonemoto, D.; Castellano, F. N.; Abolhasani, M. Continuous Biphasic Chemical Processes in a Four-Phase Segmented Flow Reactor. React. Chem. Eng. 2021, 6 (8), 1367–1375. 10.1039/D1RE00247C.

(26) Kreutz, J. E.; Shukhaev, A.; Du, W.; Druskin, S.; Daugulis, O.; Ismagilov, R. F. Evolution of Catalysts Directed by Genetic Algorithms in a Plug-Based Microfluidic Device Tested with Oxidation of Methane by Oxygen. J. Am. Chem. Soc. 2010, 132 (9), 3128–3132. 10.1021/ja909853x.

(27) Beulig, R. J.; Warias, R.; Heiland, J. J.; Ohla, S.; Zeitler, K.; Belder, D. A Droplet-Chip/Mass Spectrometry Approach to Study Organic Synthesis at Nanoliter Scale. Lab. Chip 2017, 17 (11), 1996–2002. 10.1039/C7LC00313G.

(28) Nieuwelink, A.-E.; Vollenbroek, J. C.; Tiggelaar, R. M.; Bomer, J. G.; van den Berg, A.; Odijk, M.; Weckhuysen, B. M. High-Throughput Activity Screening and Sorting of Single Catalyst Particles with a Droplet Microreactor Using Dielectrophoresis. Nat. Catal. 2021, 4 (12), 1070–1079. 10.1038/s41929-021-00718-7.

(29) Sun, A. C.; Steyer, D. J.; Allen, A. R.; Payne, E. M.; Kennedy, R. T.; Stephenson, C. R. J. A Droplet Microfluidic Platform for High-Throughput Photochemical Reaction Discovery. Nat. Commun. 2020, 11 (1), 6202. 10.1038/s41467-020-19926-z.

(30) Blomberg, R.; Kries, H.; Pinkas, D. M.; Mittl, P. R. E.; Grütter, M. G.; Privett, H. K.; Mayo, S. L.; Hilvert, D. Precision Is Essential for Efficient Catalysis in an Evolved Kemp Eliminase. Nature 2013, 503 (7476), 418–421. 10.1038/nature12623.

(31) Ma, F.; Chung, M. T.; Yao, Y.; Nidetz, R.; Lee, L. M.; Liu, A. P.; Feng, Y.; Kurabayashi, K.; Yang, G.-Y. Efficient Molecular Evolution to Generate Enantioselective Enzymes Using a Dual-Channel Microfluidic Droplet Screening Platform. Nat. Commun. 2018, 9 (1), 1030. 10.1038/s41467-018-03492-6.

(32) Mei, F.; Lin, H.; Hu, L.; Dou, W.-T.; Yang, H.-B.; Xu, L. Homogeneous, Heterogeneous, and Enzyme Catalysis in Microfluidics Droplets. Smart Mol. 2023, 1 (1), e20220001. 10.1002/smo.20220001.

(33) Holland-Moritz, D. A.; Wismer, M. K.; Mann, B. F.; Farasat, I.; Devine, P.; Guetschow, E. D.; Mangion, I.; Welch, C. J.; Moore, J. C.; Sun, S.; Kennedy, R. T. Mass Activated Droplet Sorting (MADS) Enables High-Throughput Screening of Enzymatic Reactions at Nanoliter Scale. Angew. Chem. Int. Ed. 2020, 59 (11), 4470–4477. 10.1002/anie.201913203.

(34) Brower, K. K.; Khariton, M.; Suzuki, P. H.; Still, C.; Kim, G.; Calhoun, S. G. K.; Qi, L. S.; Wang, B.; Fordyce, P. M. Double Emulsion Picoreactors for High-Throughput Single-Cell Encapsulation and Phenotyping via FACS. Anal. Chem. 2020, 92 (19), 13262–13270. 10.1021/acs.anal-chem.0c02499.

(35) Brower, K. K.; Carswell-Crumpton, C.; Klemm, S.; Cruz, B.; Kim, G.; Calhoun, S. G. K.; Nichols, L.; Fordyce, P. M. Double Emulsion Flow Cytometry with High-Throughput Single Droplet Isolation and Nucleic Acid Recovery. Lab. Chip 2020, 20 (12), 2062–2074. 10.1039/D0LC00261E.

(36) Nuti, N.; Rottmann, P.; Stucki, A.; Koch, P.; Panke, S.; Dittrich, P. S. A Multiplexed Cell-Free Assay to Screen for Antimicrobial Peptides in Double Emulsion Droplets. Angew. Chem. Int. Ed. 2022, 61 (13), e202114632. 10.1002/anie.202114632.

(37) Stucki, A.; Vallapurackal, J.; Ward, T. R.; Dittrich, P. S. Droplet Microfluidics and Directed Evolution of Enzymes: An Intertwined Journey. Angew. Chem. Int. Ed Engl. 2021, 60 (46), 24368–24387. 10.1002/anie.202016154.

(38) Reddy, S. R.; Fogler, H. S. Emulsion Stability: Determination from Turbidity. J. Colloid Interface Sci. 1981, 79 (1), 101–104. 10.1016/0021-9797(81)90052-7.

(39) Woodard, L. F.; Jasman, R. L. Stable Oil-in-Water Emulsions: Preparation and Use as Vaccine Vehicles for Lipophilic Adjuvants. Vaccine 1985, 3 (2), 137–144. 10.1016/0264-410X(85)90063-5.

(40) Holstein, J. M.; Gylstorff, C.; Hollfelder, F. Cell-Free Directed Evolution of a Protease in Microdroplets at Ultrahigh Throughput. ACS Synth. Biol. 2021, 10 (2), 252–257. 10.1021/acssynbio.0c00538.

(41) Manteca, A.; Gadea, A.; Van Assche, D.; Cossard, P.; Gillard-Bocquet, M.; Beneyton, T.; Innis, C. A.; Baret, J.-C. Directed Evolution in Drops: Molecular Aspects and Applications. ACS Synth. Biol. 2021, 10 (11), 2772–2783. 10.1021/acssyn-bio.1c00313.

(42) Taniike, T.; Takahashi, K. The Value of Negative Results in Data-Driven Catalysis Research. Nat. Catal. 2023, 6 (2), 108–111. 10.1038/s41929-023-00920-9.

(43) Zhai, S.; Xie, H.; Cui, P.; Guan, D.; Wang, J.; Zhao, S.; Chen, B.; Song, Y.; Shao, Z.; Ni, M. A Combined Ionic Lewis Acid Descriptor and Machine-Learning Approach to Prediction of Efficient Oxygen Reduction Electrodes for Ceramic Fuel Cells. Nat. Energy 2022, 7 (9), 866–875. 10.1038/s41560-022-01098-3.

(44) Kim, S.-H.; Weitz, D. A. One-Step Emulsification of Multiple Concentric Shells with Capillary Microfluidic Devices. Angew. Chem. Int. Ed. 2011, 50 (37), 8731–8734. 10.1002/anie.201102946.

(45) Bawazer, L. A.; McNally, C. S.; Empson, C. J.; Marchant, W. J.; Comyn, T. P.; Niu, X.; Cho, S.; McPherson, M. J.; Binks, B. P.; deMello, A.; Meldrum, F. C. Combinatorial Microfluidic Droplet Engineering for Biomimetic Material Synthesis. Sci. Adv. 2016, 2 (10), e1600567. 10.1126/sci-adv.1600567.

(46) Peixoto, R. S.; Vermelho, A. B.; Rosado, A. S. Petroleum-Degrading Enzymes: Bioremediation and New Prospects. Enzyme Res. 2011, 2011, 475193. 10.4061/2011/475193.

(47) Stelmachowski, M. Feedstock Recycling of Waste Polymers by Thermal Cracking in Molten Metal: Thermodynamic Analysis. J. Mater. Cycles Waste Manag. 2014, 16 (2), 211–218. 10.1007/s10163-013-0178-x.

(48) Yan, G.; Jing, X.; Wen, H.; Xiang, S. Thermal Cracking of Virgin and Waste Plastics of PP and LDPE in a Semibatch Reactor under Atmospheric Pressure. Energy Fuels 2015, 29 (4), 2289–2298. 10.1021/ef502919f.

(49) Li, H.; Aguirre-Villegas, H. A.; Allen, R. D.; Bai, X.; Benson, C. H.; Beckham, G. T.; Bradshaw, S. L.; Brown, J. L.; Brown, R. C.; Cecon, V. S.; Curley, J. B.; Curtzwiler, G. W.; Dong, S.; Gaddameedi, S.; García, J. E.; Hermans, I.; Kim, M. S.; Ma, J.; Mark, L. O.; Mavrikakis, M.; Olafasakin, O. O.; Osswald, T. A.; Papanikolaou, K. G.; Radhakrishnan, H.; Sanchez Castillo, M. A.; Sánchez-Rivera, K. L.; Tumu, K. N.; Van Lehn, R. C.; Vorst, K. L.; Wright, M. M.; Wu, J.; Zavala, V. M.; Zhou, P.; Huber, G. W. Expanding Plastics Recycling Technologies: Chemical Aspects, Technology Status and Challenges. Green Chem. 2022, 24 (23), 8899–9002. 10.1039/D2GC02588D.

(50) Tan, Y. H.; Abdullah, M. O.; Nolasco-Hipolito, C. The Potential of Waste Cooking Oil-Based Biodiesel Using Heterogeneous Catalyst Derived from Various Calcined Eggshells Coupled with an Emulsification Technique: A Review on the Emission Reduction and Engine Performance. Renew. Sustain. Energy Rev. 2015, 47, 589–603. 10.1016/j.rser.2015.03.048.

