## Supporting Information for "FACS-Sortable Triple Emulsion Picoreactors for Screening Reactions in Biphasic Environments"

**Supplementary Files**

Supplementary files are separate files available at https://osf.io/gbq5r. All data are paired with any custom python scripts used for analysis. Data Supplement 16 includes workspace files for the FlowJo commercial software suite.

**Data Supplements 1-8** are .csv files that contain plate reader absorbance and turbidity measurements.

**Data Supplement 1:** ***Surfactant combination screen for octane.***

**Data Supplement 2:** ***Surfactant combination screen for hexyl acetate.***

**Data Supplement 3:** ***Surfactant combination screen for 1-octanol.***

**Data Supplement 4:** ***Surfactant combination screen for 2-octanone.***

**Data Supplement 5:** ***Surfactant concentration screen for octane.***

**Data Supplement 6:** ***Surfactant concentration screen for hexyl acetate.***

**Data Supplement 7:** ***Surfactant concentration screen for 1-octanol.***

**Data Supplement 8:** ***Surfactant concentration screen for 2-octanone.***

**Data Supplements 9-10** are zipped directories and files of images and .csv files for the analysis of octanol/aqueous emulsions.

**Data Supplement 9: *Microscopy images of octanol/aqueous emulsions***

**Data Supplement 10: 1-*octanol droplet diameters***

**Data Supplements 11-16** are zipped directories and files of images, .csv, and .fsc files for the analysis of triple emulsion picoreactors.

**Data supplement 11: *Platereader data for Nile Red extractions***

**Data supplement 12: *Microscopy images of triple emulsions***

**Data supplement 13: *Fluorescence quantification for IVTT in the presence of hydrocarbon solvents***

**Data supplement 14: *Fluorescence quantification for IVTT in the presence of surfactants***

**Data supplement 15: *Microscopy images of triple emulsions with IVTT reagents***

**Data supplement 16: *Microscopy images of triple emulsions for FACS analysis and sorting***

**Data supplement 17: *FACS events and gates and FlowJo workspace files***

**Supplementary Tables**

Supplementary Table 2 is a separate .csv file available at https://osf.io/gbq5r. Supplementary Tables 1, 3, and 4 are included below

**Supplementary Table 1** compiles literature data on solvent miscibility.

**Supplementary Table 2** contains a summary of related studies.

**Supplementary Table 3** compiles literature data on surfactant physiochemical properties.

**Supplementary Table 4** contains statistics for FACS sorting.

**Supplementary Figures**

**Supplementary Figure 1: *Diagram of double and triple emulsion picoreactors.***

**Supplementary Figure 2: *Platform for picoreactor droplet generation.***

**Supplementary Figure 3: *Encapsulation of 1-octanol/aqueous biphasic solutions inside a fluorocarbon shell requires surfactant optimization.***

**Supplementary Figure 4: *Relationship between hydrocarbon solvent miscibility in water and solvent/surfactant hydrophobic-lipophobic balance.***

**Supplementary Figure 5: *Plate reader turbidity measurements for hydrocarbon solvent/surfactant combinations at 2 hours.***

**Supplementary Figure 6: *Changes in plate reader turbidity measurements for hydrocarbon solvent/surfactant combinations over time.***

**Supplementary Figure 7: *Reproducibility of plate reader turbidity measurements across wells for initial hydrocarbon solvent/surfactant combinations at 24 hours.***

**Supplementary Figure 8: *Well images for octane/surfactant combinations at 24 hours.***

**Supplementary Figure 9: *Well images for hexyl acetate/surfactant combinations at 24 hours.***

**Supplementary Figure 10: *Well images for 1-octanol/surfactant combinations at 24 hours.***

**Supplementary Figure 11: *Well images for 2-octanone/surfactant combinations at 24 hours.***

**Supplementary Figure 12: *Correspondence between microscopy and plate-based turbidity measurements used for emulsion characterization.***

**Supplementary Figure 13: *Fluorescence microscopy images of selected emulsions.***

**Supplementary Figure 14: *Quantifying droplet size distribution in an octanol /aqueous emulsion.***

**Supplementary Figure 15: *Plate reader turbidity measurements for surfactant combinations as a function of concentration at 2 hours.***

**Supplementary Figure 16: *Changes in plate reader turbidity measurements for surfactant combinations as a function of concentration between 24 hours vs. 2 hours.***

**Supplementary Figure 17: *Reproducibility of plate reader turbidity measurements across wells for optimal hydrocarbon solvent/surfactant combinations at 24 hours.***

**Supplementary Figure 18: *Well images for optimized octane/surfactant combinations at multiple concentrations at 24 hours.***

**Supplementary Figure 19: *Well images for optimized hexyl acetate/surfactant combinations at multiple concentrations after 24 hours.***

**Supplementary Figure 20: *Well images for optimized 1-octanol/surfactant combinations at multiple concentrations after 24 hours.***

**Supplementary Figure 21: *Well images for optimized 2-octanone/surfactant combinations at multiple concentrations at 24 hours.***

**Supplementary Figure 22: *Nile Red selectively partitions into 1-octanol.***

**Supplementary Figure 23: *Automated detection of triple emulsions.***

**Supplementary Figure 24: *Distribution of droplet sizes.***

**Supplementary Figure 25: *Distribution of droplet sizes with IVTT reagents.***

**Supplementary Figure 26: *Double and Triple Emulsion populations in samples for FACS analysis and sorting.***

**Supplementary Figure 27: *Distribution of droplet sizes for FACS analysis and sorting.***

**Extended Methods**

**Screening surfactants**

***Calculating HLB***

***Plate reader turbidity measurements***

***Microscopy reader turbidity measurements***

***Testing Nile Red partitioning***

***Generating microfluidic devices***

**Operating microfluidic devices**

***Generating double emulsions***

***Generating triple emulsions with pre-emulsified octanol/aqueous inner solutions***

***Plasmid construction and miniprep***

***Expressing protein in the presence of hydrocarbon solvents and surfactants***

***Generating triple emulsions for expressing protein in biphasic droplet picoreactors with octanol/aqueous emulsion cores***

***Imaging droplets and emulsions***

**FACS sorting microfluidics droplets**

**Supplementary Tables**

**Supplementary Table 1. *Solvent miscibility in water***

| **Solvent** | **Miscibility in water (g/L)** | **Temperature** | **References** |
| --- | --- | --- | --- |
| Water | 1000 (density of water) | 25˚C |  |
| Butanol | 63.2 | 25˚C | NIH PubChem^46^ |
| Hexanol | 5.9 | 25˚C | NIH PubChem^47^ |
| 1-octanol | 0.540 | 25˚C | NIH PubChem^48^ |
| 1-octanol | 0.30 | 20˚C | NIH PubChem^48^ |
| 2-octanone | 0.899 | 20˚C | NIH PubChem^49^ |
| Hexyl acetate | 0.511 | 25˚C | NIH PubChem^50^ |
| Octane | 0.00066 | 25˚C | NIH PubChem^51^ |

**Supplementary Table 3. *Surfactant physiochemical properties***

| **Surfactant** | **Soluble to 5% (w/v)** | **Class** | **HLB** | **Reference** |
| --- | --- | --- | --- | --- |
| Span 85 | 1-octanol  2-octanone  Hexyl acetate  Octane | Non-ionic | 1.8 | Gorman and Hall, 1963^3^ |
| Span 80 | 1-octanol  2-octanone  Hexyl acetate  Octane | Non-ionic | 4.3 | Gorman and Hall, 1963^3^ |
| EM-90 | 1-octanol  2-octanone  Hexyl acetate  Octane | Non-ionic | 5 | ABIL Technical data sheet |
| Span 20 | 1-octanol  2-octanone  Octane | Non-ionic | 8.6 | Gorman and Hall, 1963^3^ |
| Triton CG-110 | Water | Non-ionic | 13 | Gala Marti et al., 2021^4^ |
| Triton X-100 | 1-octanol  2-octanone  Hexyl acetate  Water | Non-ionic | 13.4 | Gala Marti et al., 2021^4^ |
| Tween 80 | 1-octanol  2-octanone  Hexyl acetate  Water | Non-ionic | 15.0 | Gorman and Hall, 1963^3^ |
| Tween 20 | 1-octanol  2-octanone  Hexyl acetate  Water | Non-ionic | 16.7 | Gorman and Hall, 1963^3^ |
| NP-40 | Water* | Non-ionic | 17.8 | Dow Technical data sheet,  Sigma detergent sheet |
| Benzalkonium Chloride | 1-octanol  2-octanone  Water | Cationic | ~18.5,  16.175-20.925 | Calculated in this work. |
| CTAB | Water | Cationic | 21.4, 10 | Miraglia et al.,2011^6^, Sigma detergent sheet |
| Sarkosyl | Water | Anionic | 29.8 | Stepan Technical data sheet, Sigma detergent sheet |
| SDS | Water | Anionic | 40 | Schramm et al., 2003^5^ |
| SB3-10  (3-(Decyldimethylammonio) propane sulfonate) | Water | Zwitterionic | 41 | Sigma detergent sheet |
| CHAPS | Water | Zwitterionic | 60.2, 10 | Calculated in this work, Sigma detergent sheet |

*Sold/distributed as an aqueous solution.

**Supplementary Table 4. *FACS Analysis Statistics***

| **Population** | **Double Emulsion** | **% of Parent** | **Triple**  **Emulsion - 4˚C** | **% of Parent** | **Triple**  **Emulsion - 37˚C** | **% of Parent** |
| --- | --- | --- | --- | --- | --- | --- |
| **Total events** | 10000 |  | 10000 |  | 10000 |  |
| **Scatter Gate**  **(SSC-A *vs.* FSC-A)** | 8526 | 85.3% | 6488 | 64.9% | 8611 | 86.1% |
| **Singlet Gate 1**  **(FSC-A *vs.* FSC-H)** | 8327 | 97.7% | 6113 | 94.2% | 8591 | 99.8% |
| **Single Gate 2**  **(SSC-A *vs.* SSC-H)** | 8303 | 99.7% | 5623 | 92.0% | 8264 | 96.2% |
| **Median GFP**  **(B525)** | 76.2 |  | 390.3 |  | 1595.9 |  |
| **Median Nile Red (G660)** | 44.6 |  |  |  | 2546.4 |  |

**Supplementary Figures and Legends**

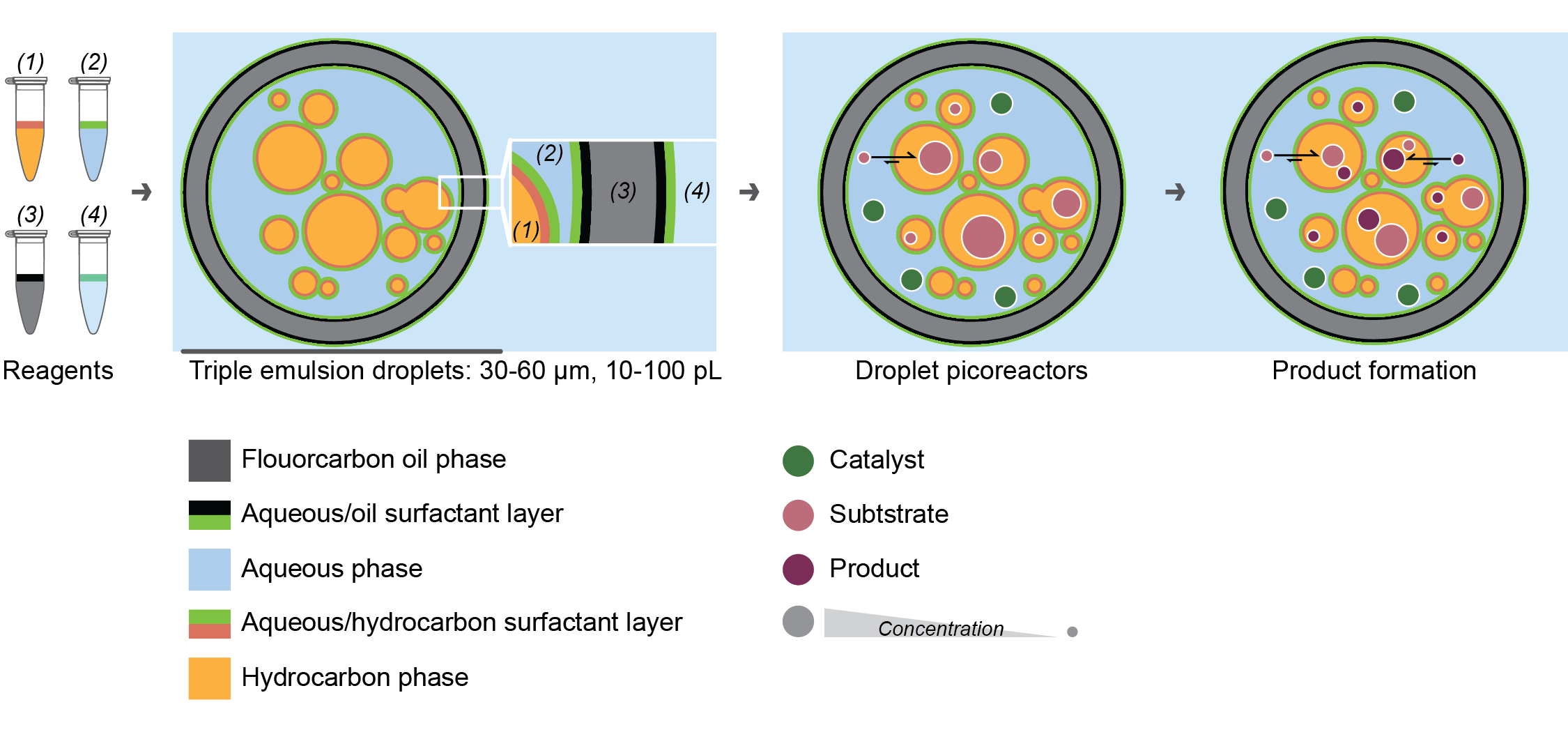

**Supplementary Figure 1: *Diagram of double and triple emulsion picoreactors.***

Triple emulsions are formed from four solutions of a liquid + surfactant: aqueous (inner), hydrocarbon (inner), fluorocarbon (shell), and aqueous (outer). Triple emulsions contain a core of hydrocarbon oil and aqueous solution wrapped in a fluorocarbon shell. The droplet is suspended in an outer aqueous phase for FACS compatibility. The hydrocarbon/aqueous micro-emulsion encapsulated in the core of the droplet can facilitate a biphasic reaction by isolating specific reaction conditions, physically linking the catalyst with the product, and optimizing interfacial surface area to accelerate substrate/product partitioning.

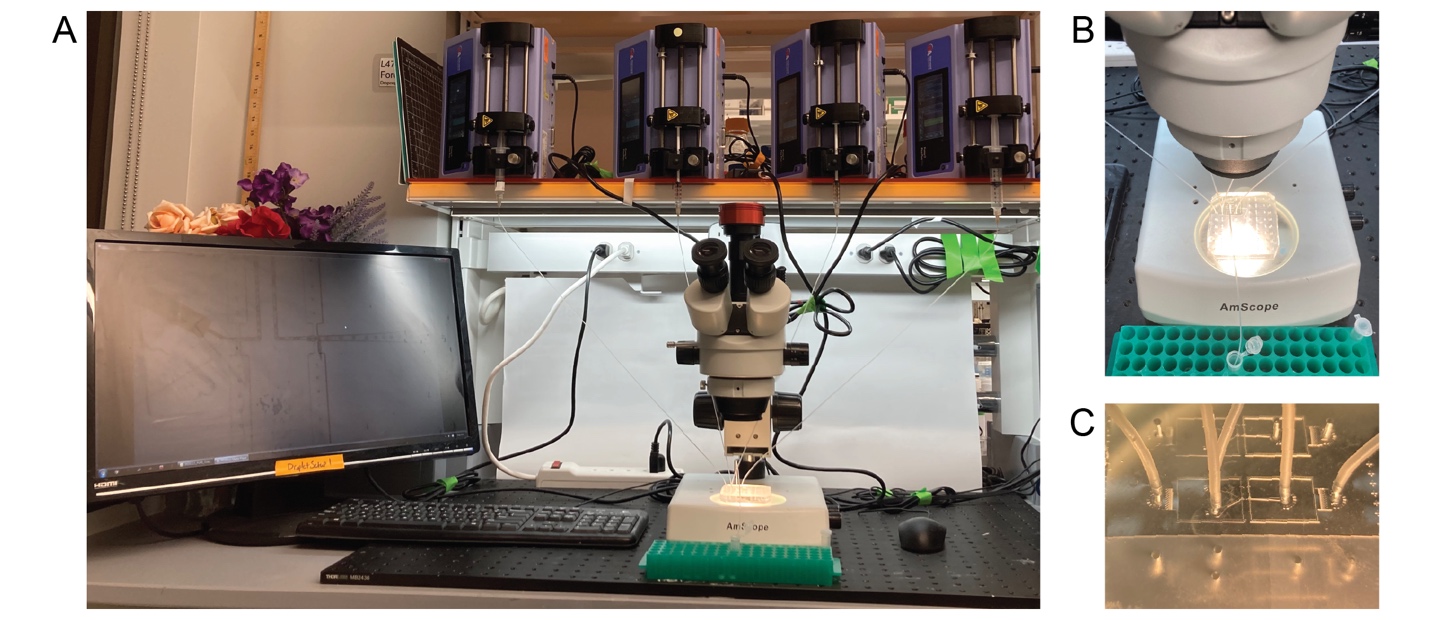

**Supplementary Figure 2: *Platform for picoreactor droplet generation.***

**A)** Droplet generation set-up with 4 syringe pumps, syringes and tubing, PDMS device, optical microscope with high-speed camera, and computer. **B)** Array of devices in PDMS block. **C)** Close-up of tubing connected to one PDMS device.

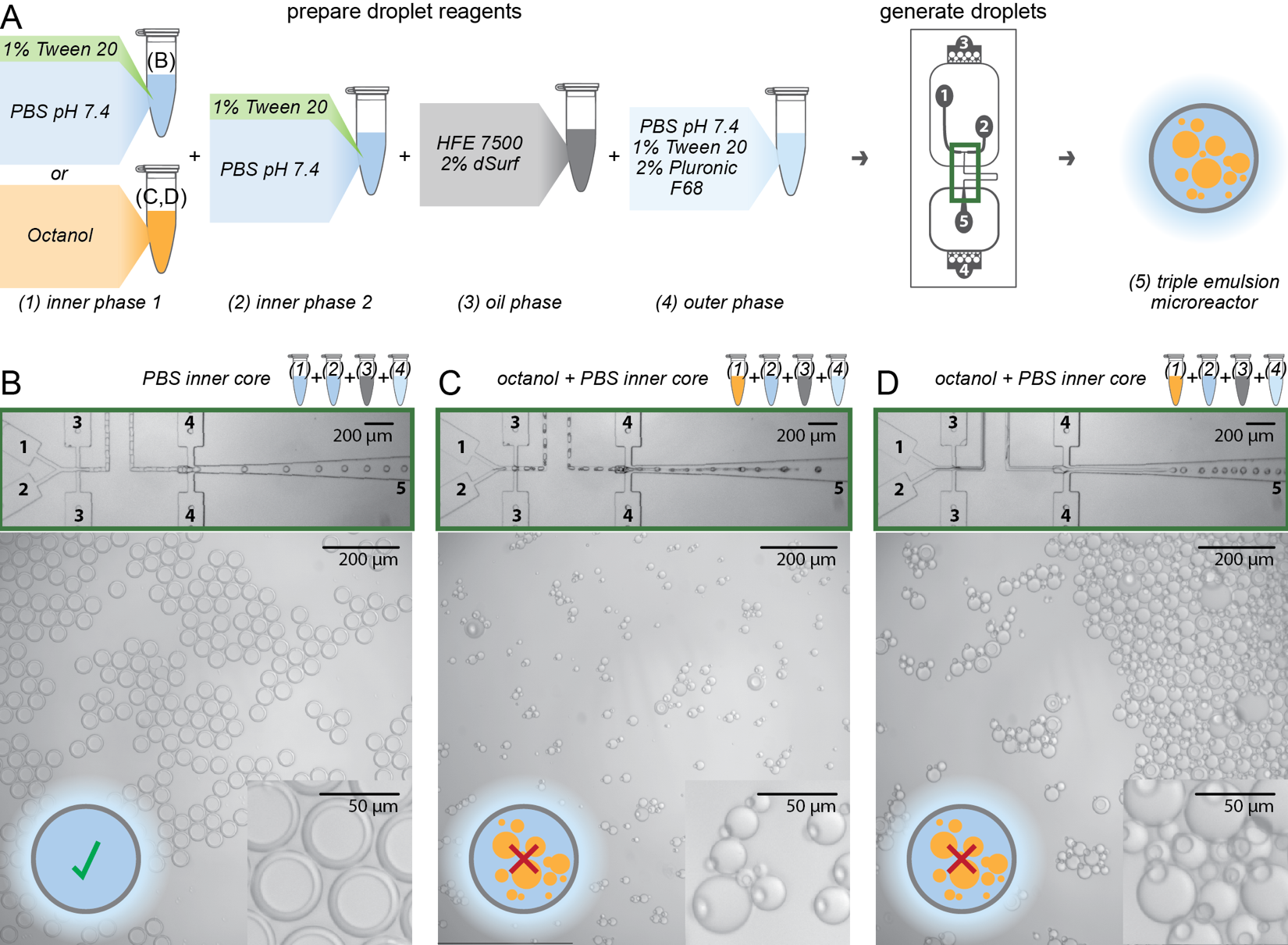

**Supplementary Figure 3: *Encapsulation of octanol/aqueous biphasic solutions inside a fluorocarbon shell requires surfactant optimization.***

**A)** Workflow for generating double (B) and triple emulsions (C,D). Surfactants are optimized only for aqueous and fluorocarbon phases. **B)** Successful generation of double emulsion droplets with view of on-chip generation, output droplets, and a zoom view of output droplets. Scale bars for each image are 200 µm, 200 µm, and 50 µm, respectively. Numbered channels match the numbers for reagents and channels from A. On-chip generation view corresponds to the device with a green outline in A. **C,D)** Without surfactants for 1-octanol, droplet formation is disrupted. Channel labels and scale bars as in B.

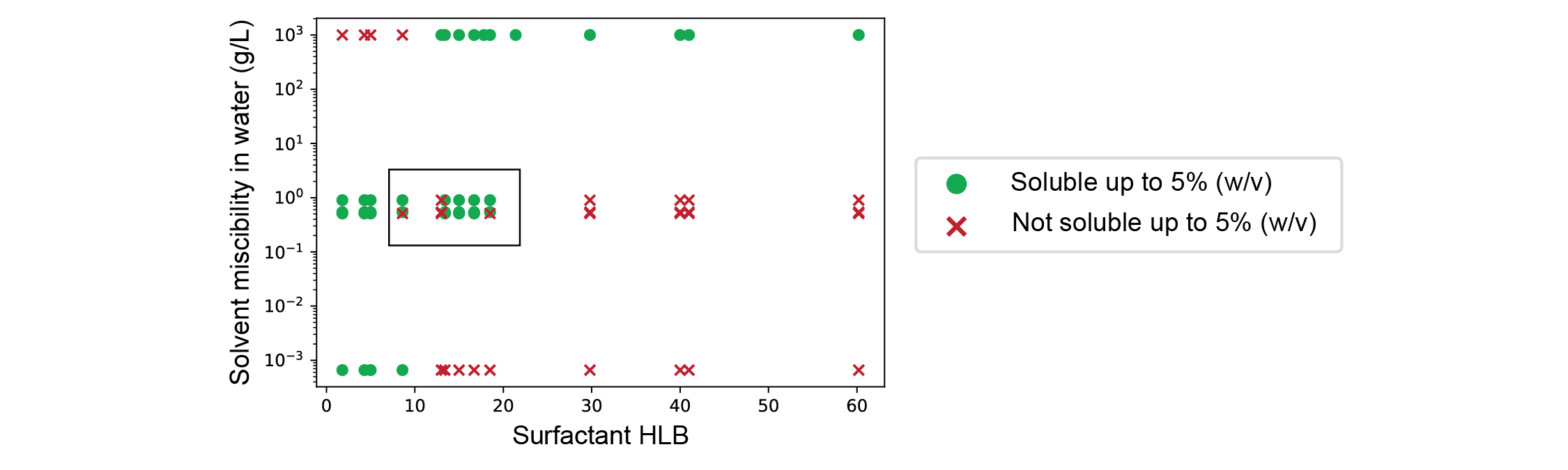

**Supplementary Figure 4: *Relationship between hydrocarbon solvent miscibility in water and hydrocarbon solvent/surfactant hydrophobic-lipophobic balance.***

Scatter plot showing hydrocarbon solvent miscibility in water for octane, hexyl acetate, 1-octanol, 2-octanone, and PBS pH 7.4 against hydrophobic-lipophobic balance (HLB) for particular solvent/surfactant combinations. Markers indicate surfactant solubility as evaluated against a threshold of 5% (w/v) with either a green circle (soluble) or red ‘X’ (not soluble); all values are given in **Supplementary Tables 1** and **2**.

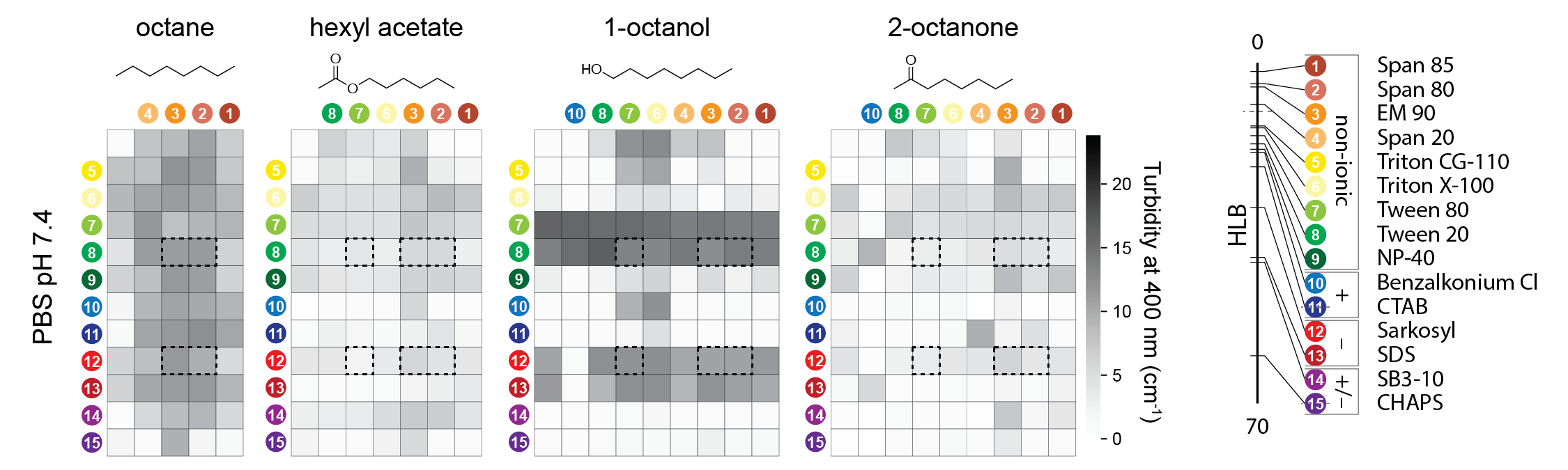

**Supplementary Figure 5: *Plate reader turbidity measurements for hydrocarbon solvent/surfactant combinations at 2 hours.***

Plate reader turbidity measurements after 2 hours for one aqueous buffer (PBS pH 7.4) and 4 hydrocarbon solvents. Surfactant labels (right) reproduced from **Figure 2B**. Data after 24 hours shown in Figure 2C; dashed black lines indicate selected surfactant combinations shown in **Figures 2D**,**E**.

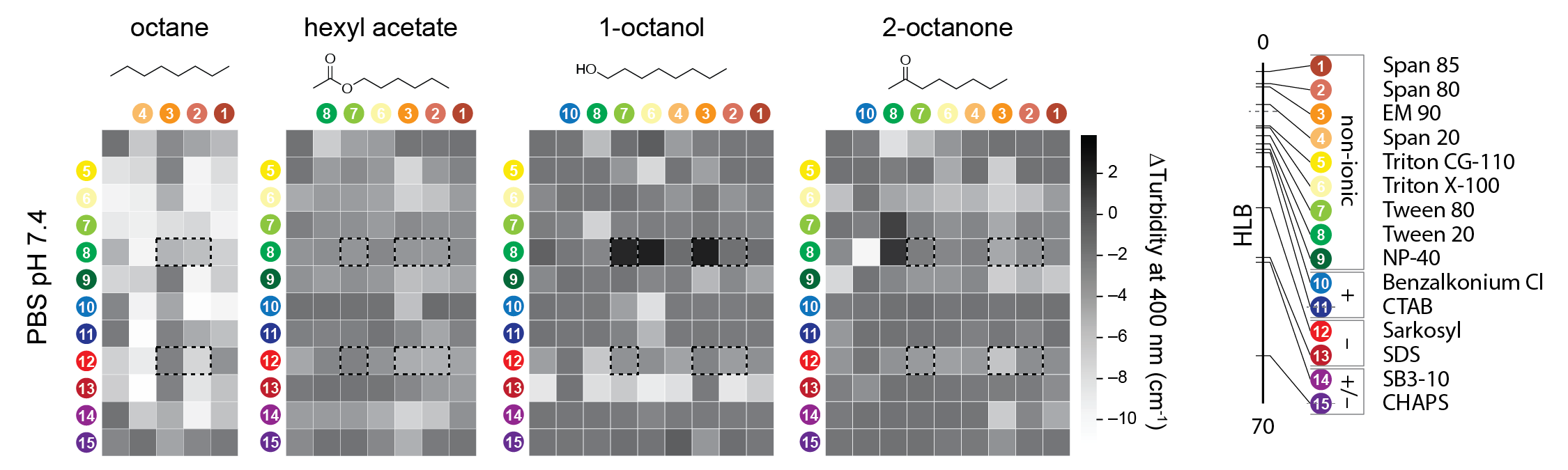

**Supplementary Figure 6: *Changes in plate reader turbidity measurements for hydrocarbon solvent/surfactant combinations over time.***

Change in plate reader turbidity measurements between 2 hours and 24 hours for one aqueous buffer (PBS pH 7.4) and 4 hydrocarbon solvents. Surfactant labels (right) reproduced from **Figure 2B**; dashed black lines indicate selected surfactant combinations shown in **Figures 2D**,**E**.

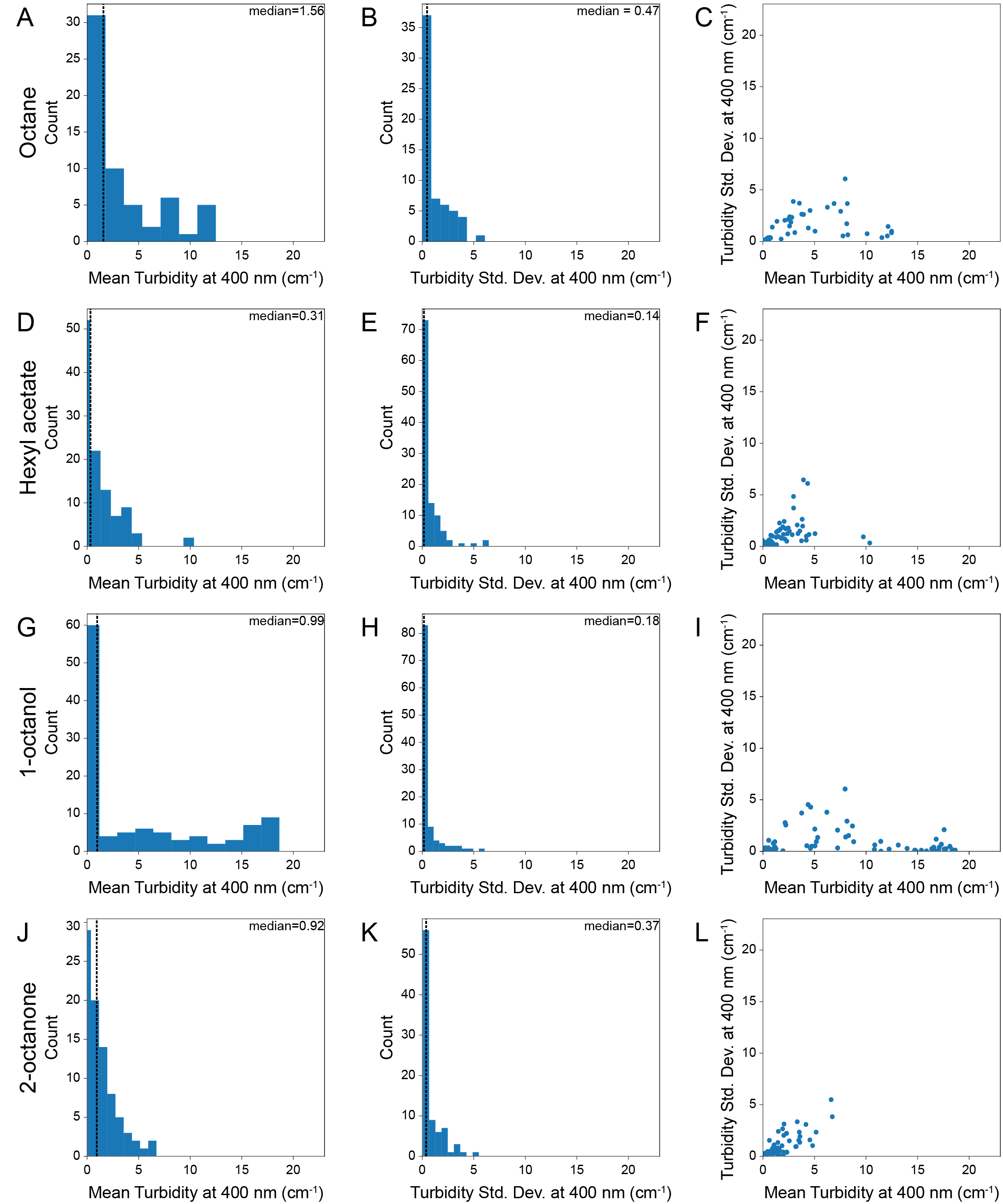

**Supplementary Figure 7: *Reproducibility of plate reader turbidity measurements across wells for initial hydrocarbon solvent/surfactant combinations at 24 hours.***

Plate reader turbidity measurements after 24 hours for one aqueous buffer (PBS pH 7.4) and 4 hydrocarbon solvents corresponding to data in in **Figure 2C**. Histograms of mean turbidity (n=3), histograms of turbidity standard deviation (n=3), and scatter plots of mean *vs.* standard deviation for combinations of surfactants at 5% (w/v) in octane (**A**-**C**), hexyl acetate (**D**-**F**), 1-octanol (**G**-**I**), and 2-octanone (**J**-**L**).

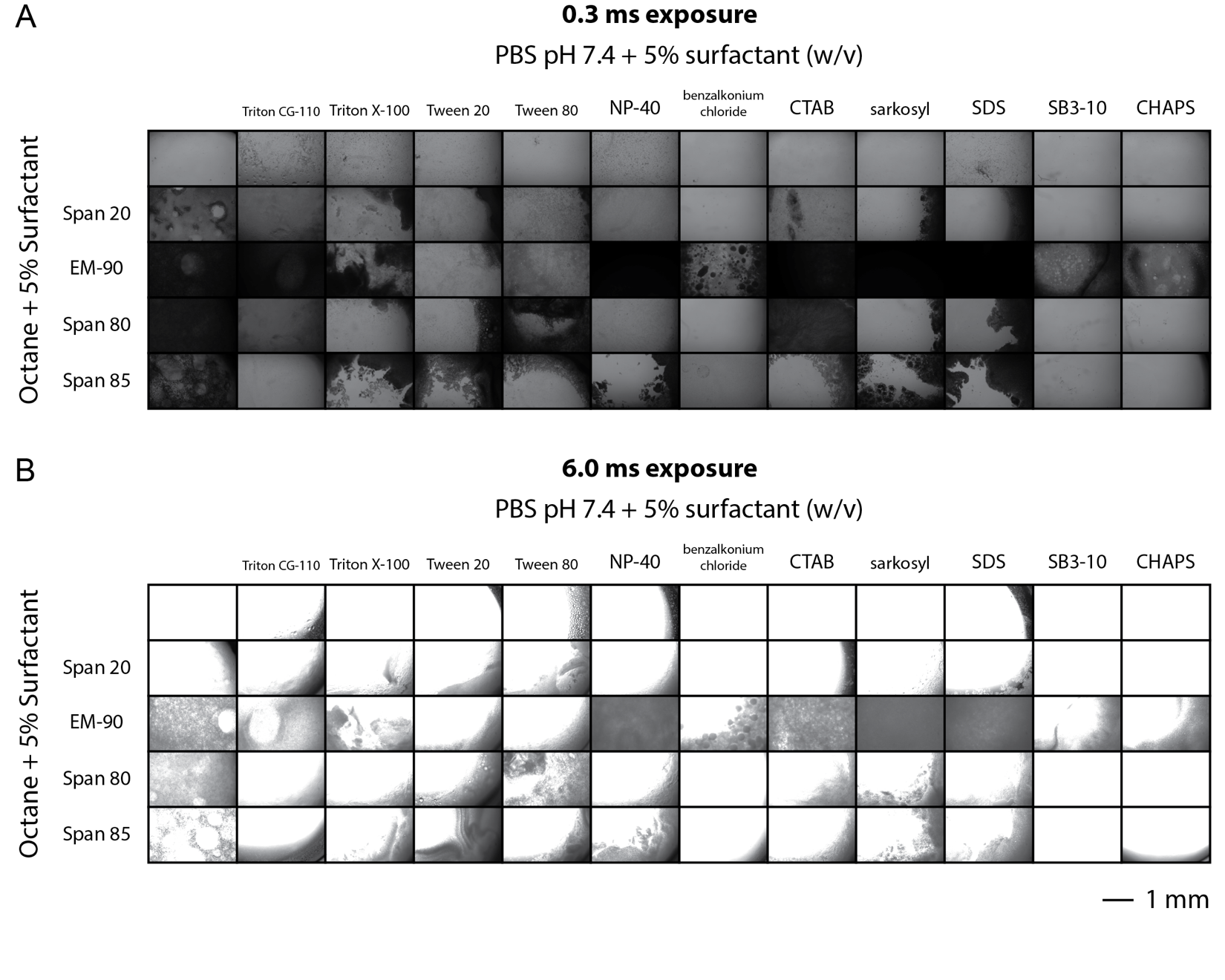

Legend on following page.

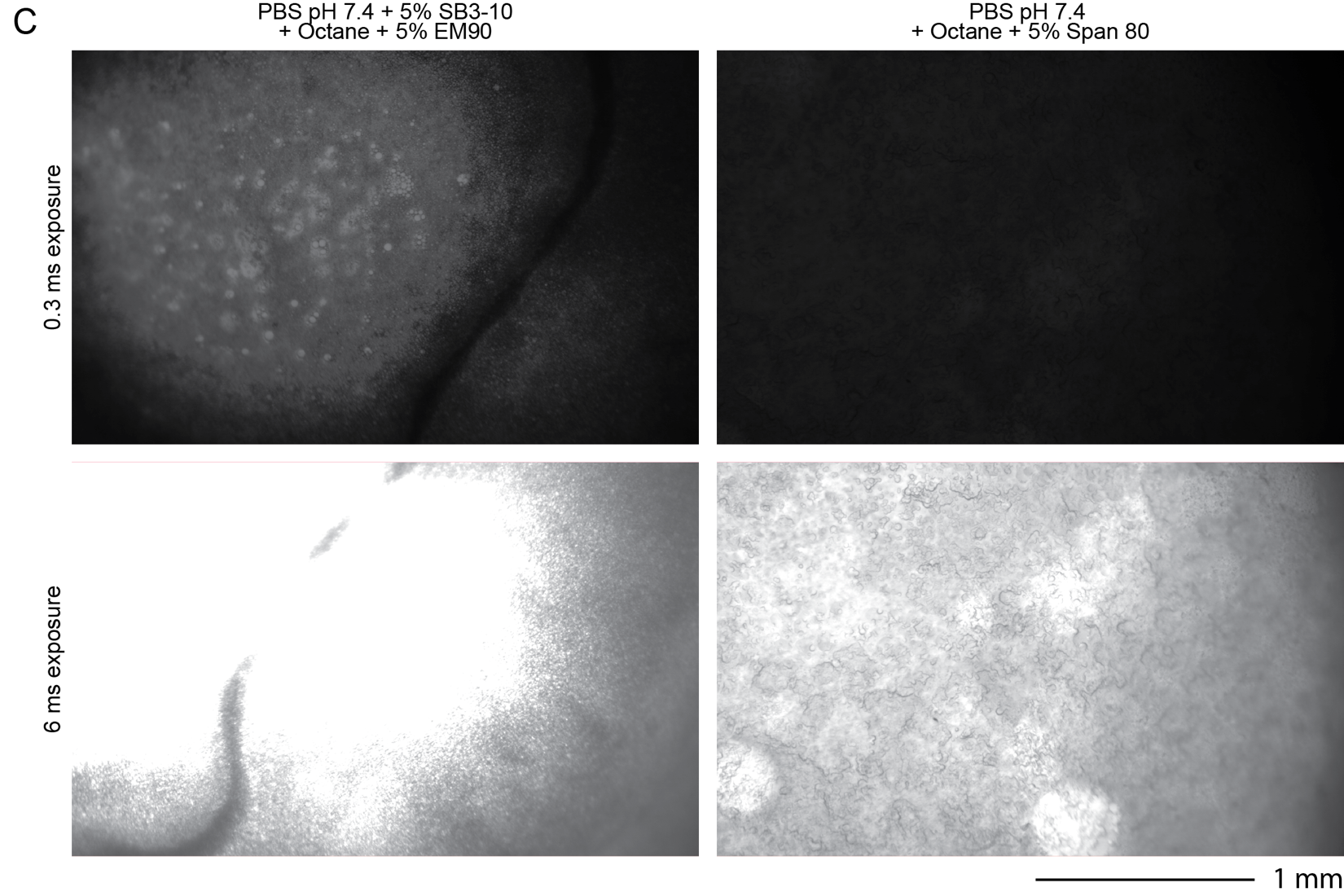

**Supplementary Figure 8: *Well images for octane/surfactant combinations at 24 hours.***

**A)** 0.3 ms exposure and **B)** 6.0 ms exposure images of wells from plate reader turbidity assay to screen surfactants for aqueous-octane emulsions. Scale bar: 1 mm. Images correspond to data in **Figure 2C**. **C)** Images of selected conditions. Scale bar: 1 mm.

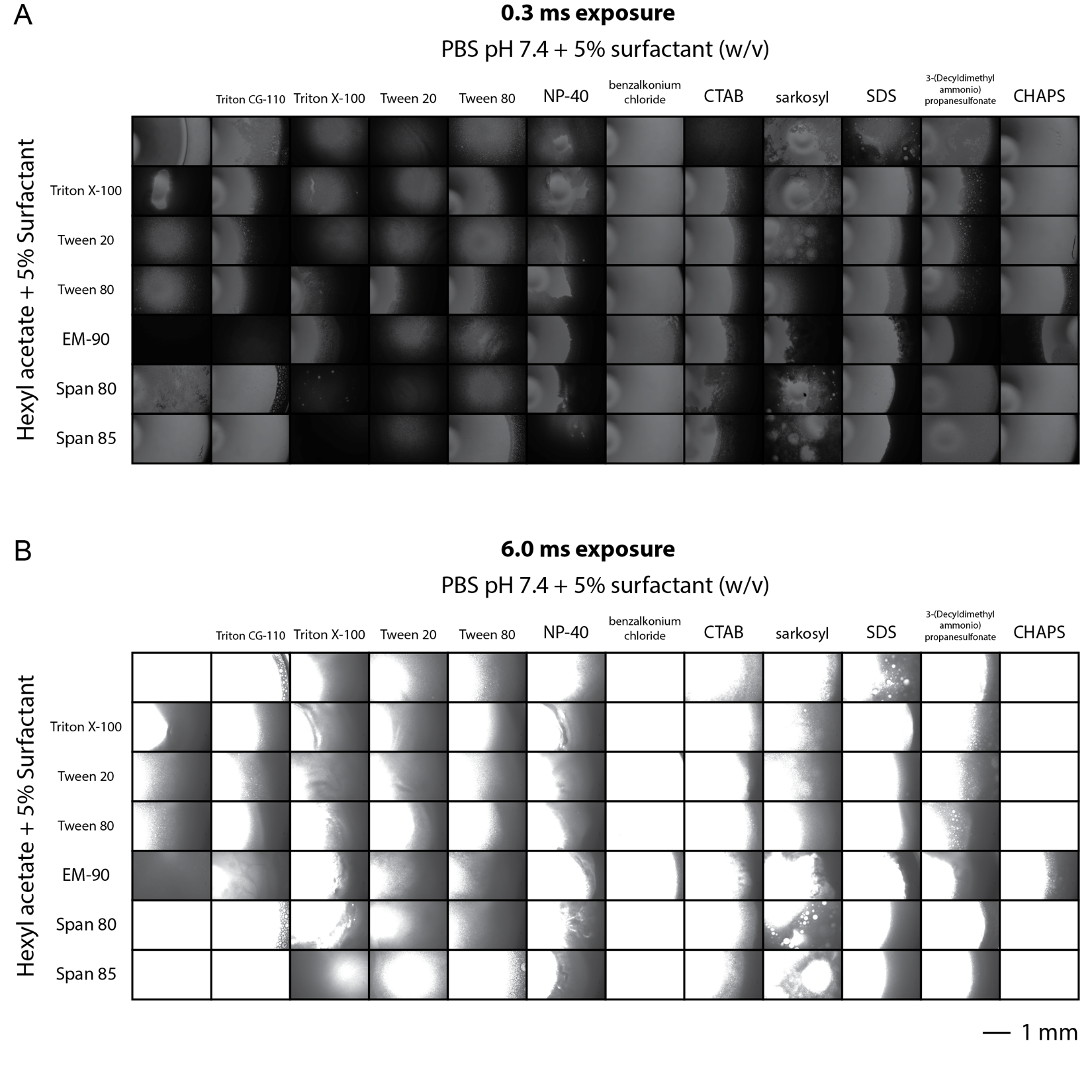

Legend on following page.

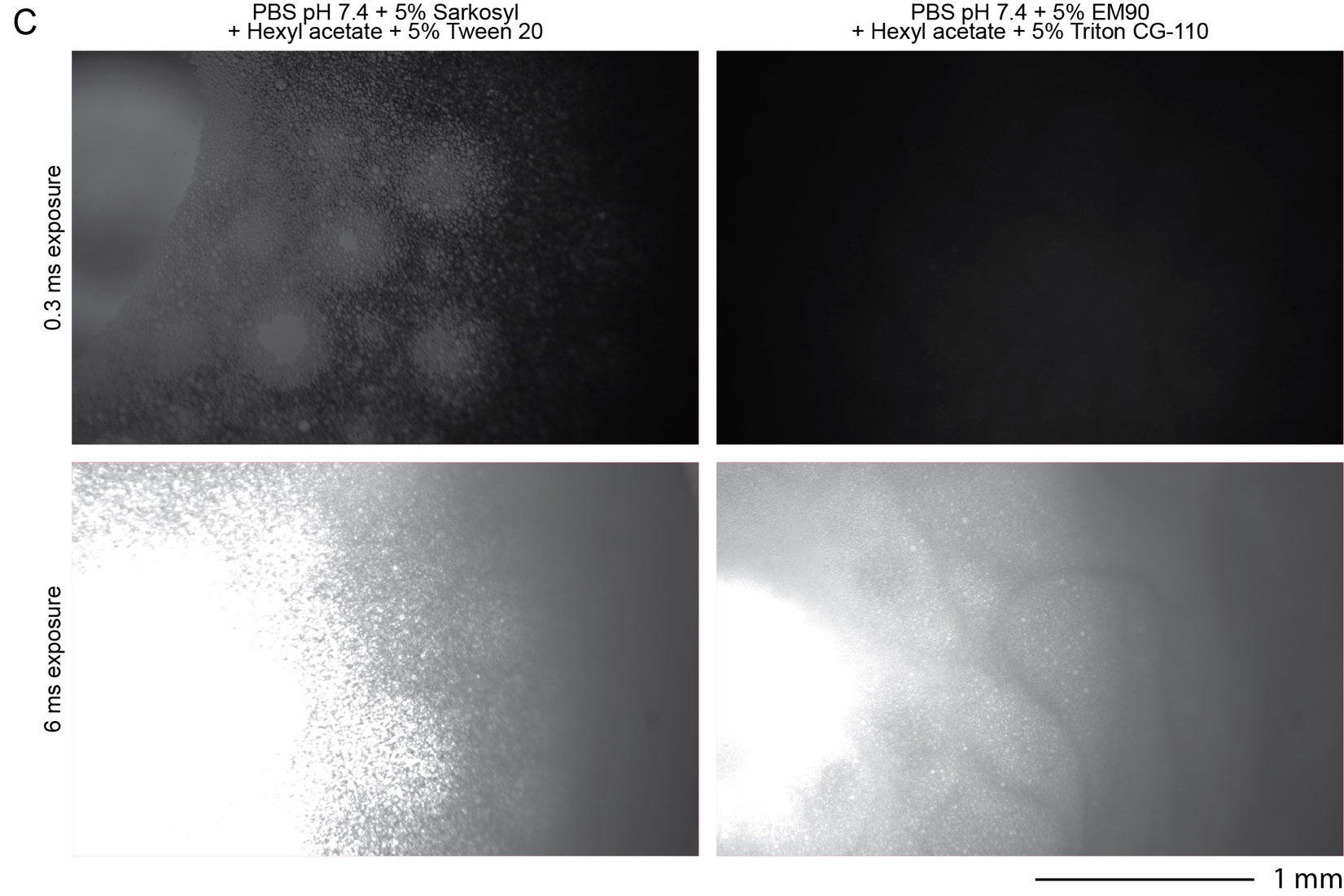

**Supplementary Figure 9: *Well images for hexyl acetate/surfactant combinations at 24 hours.***

**A)** 0.3 ms exposure and **B)** 6.0 ms exposure images of wells from plate reader turbidity assay to screen surfactants for aqueous-hexyl acetate emulsions. Scale bar: 1 mm. Images correspond to data in **Figure 2C**. **C)** Images of selected conditions. Scale bar: 1 mm.

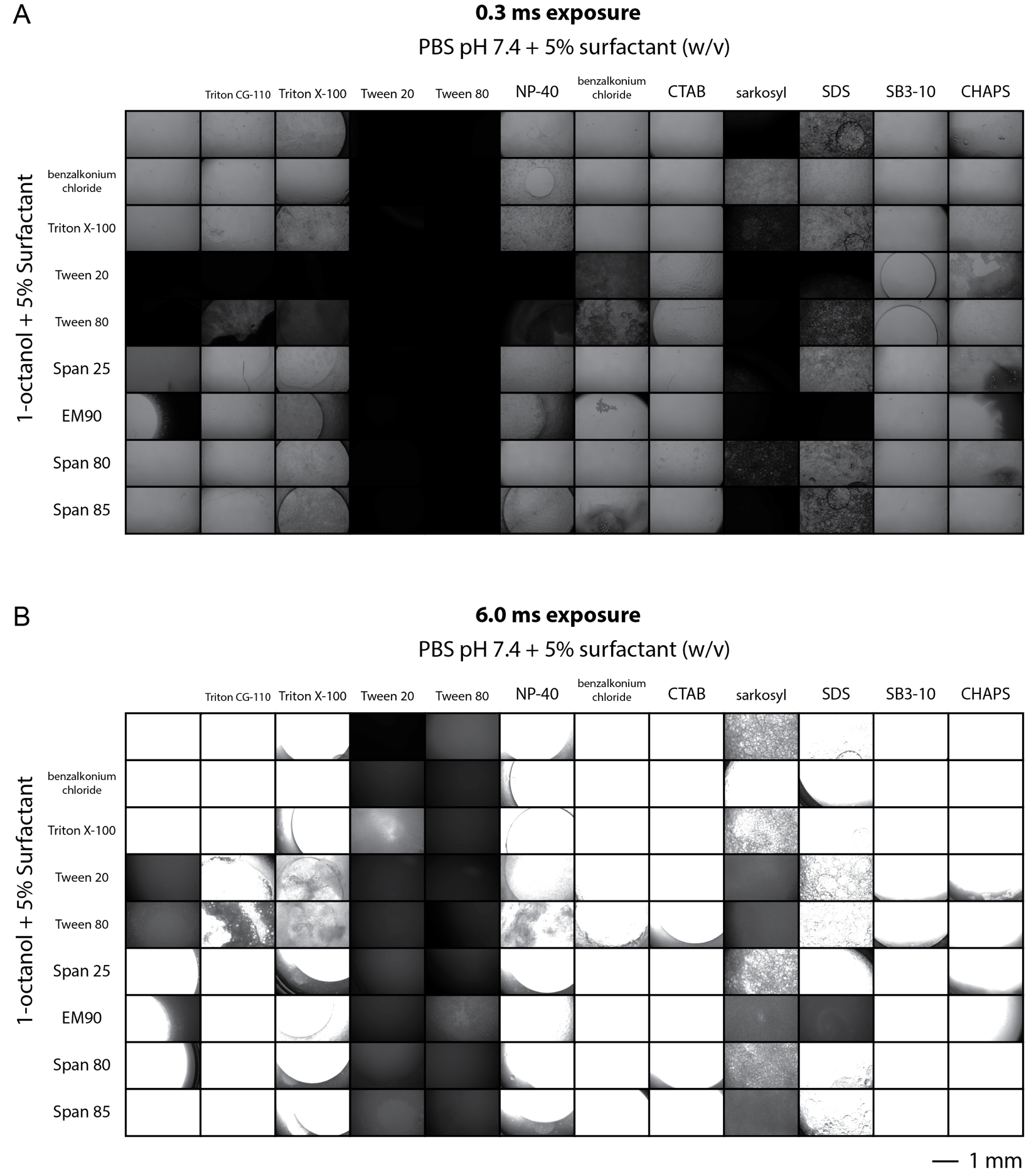

Legend on following page.

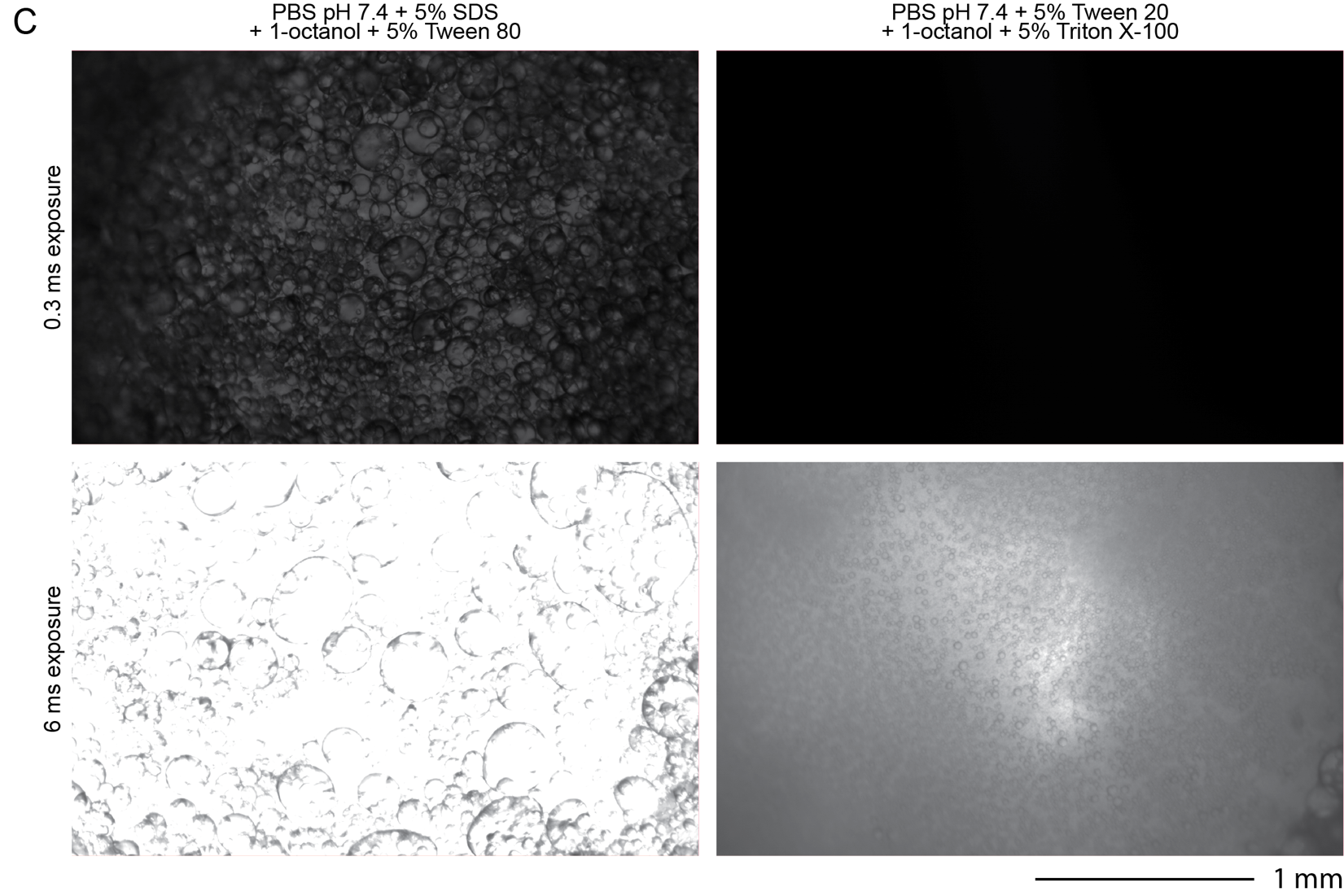

**Supplementary Figure 10: *Well images for 1-octanol/surfactant combinations at 24 hours.***

**A)** 0.3 ms exposure and **B)** 6.0 ms exposure images of wells from plate reader turbidity assay to screen surfactants for octanol/aqueous emulsions. Scale bar: 1 mm. Images correspond to data in **Figure 2C**. **C)** Images of selected conditions. Scale bar: 1 mm.

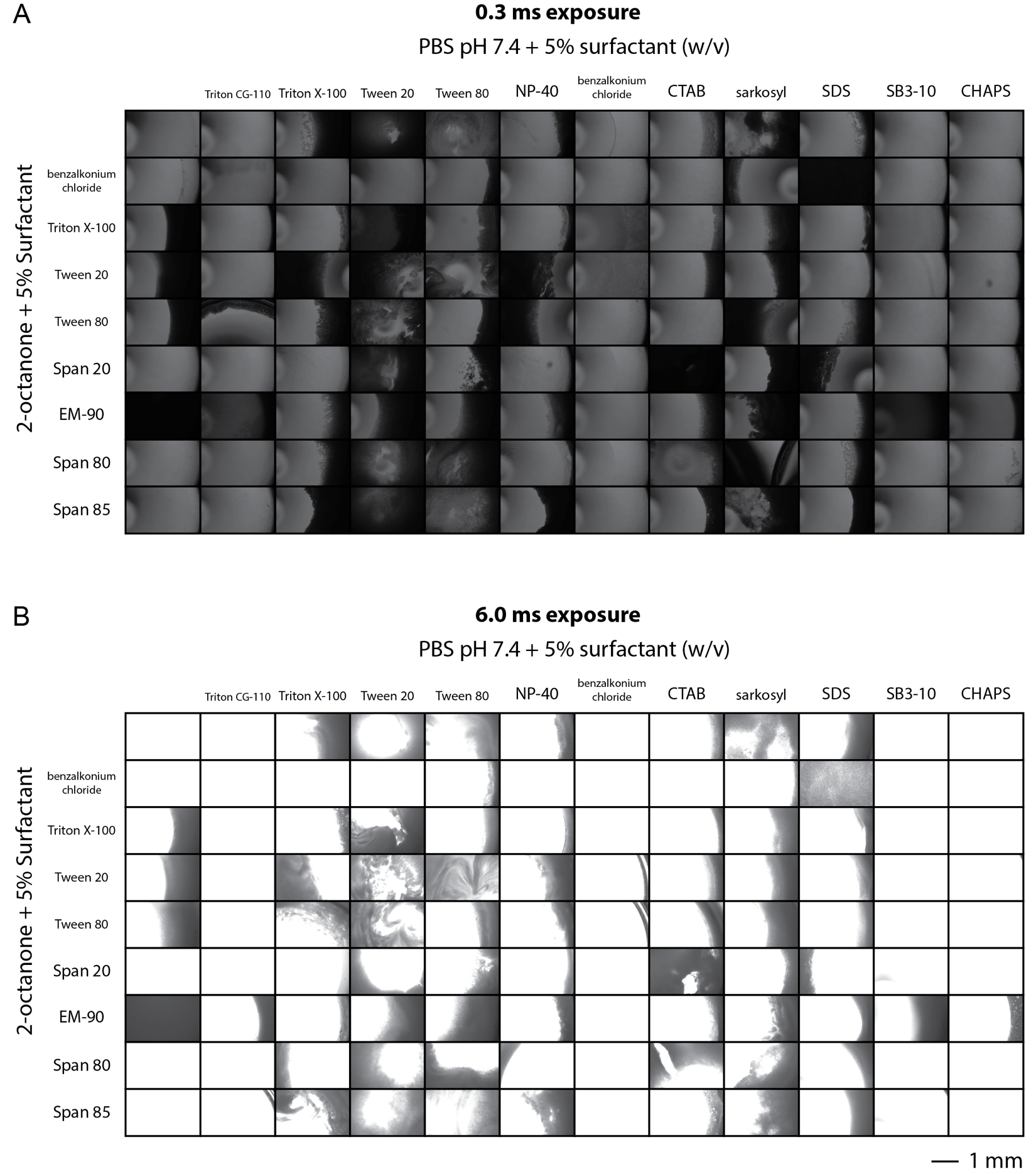

Legend on following page.

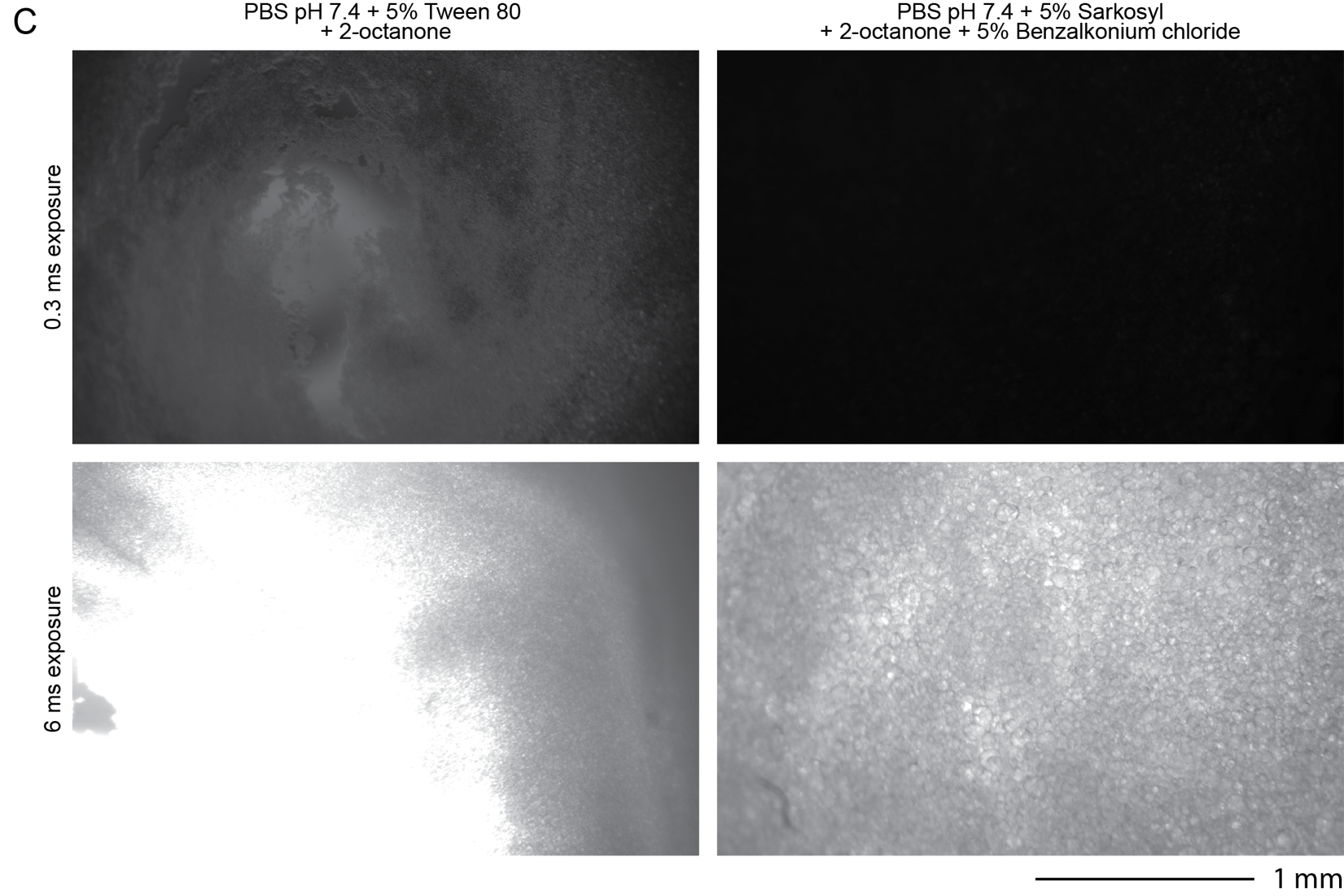

**Supplementary Figure 11: *Well images for 2-octanone/surfactant combinations at 24 hours.***

**A)** 0.3 ms exposure and **B)** 6.0 ms exposure images of wells from plate reader turbidity assay to screen surfactants for aqueous-octanone emulsions. Scale bar: 1 mm. Images correspond to data in **Figure 2C**. **C)** Images of selected conditions. Scale bar: 1 mm.

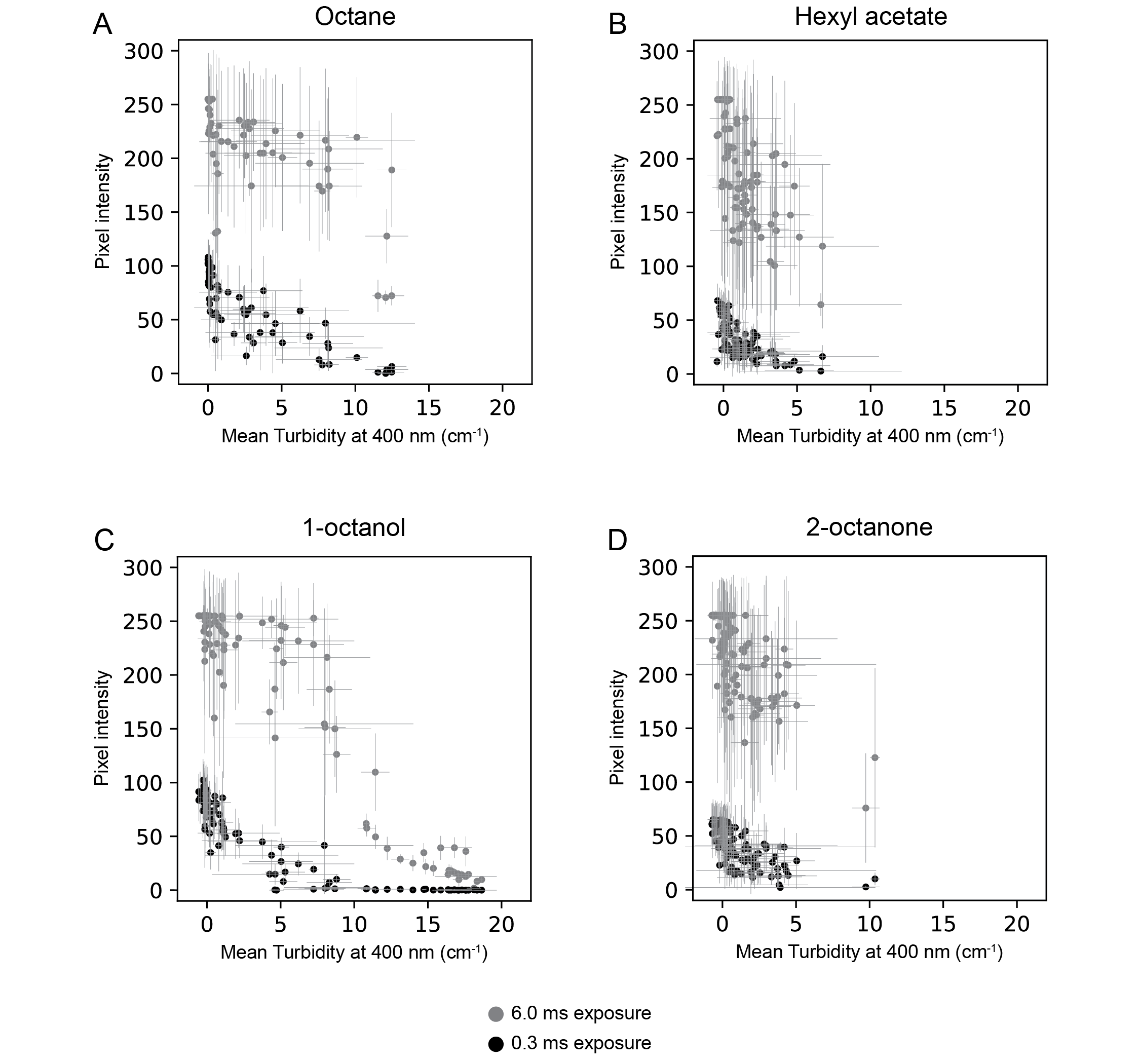

**Supplementary Figure 12: *Correspondence between microscopy and plate-based turbidity measurements used for emulsion characterization.***

Mean pixel intensities *vs.* mean turbidity from plate-based measurements for well images of aqueous emulsions with **A)** octane, **B)** hexyl acetate, **C)** 1-octanol, and **D)** 2-octanone. Well images are shown in **Supplementary figures 8-11**. Points for 6.0 ms exposure images are colored grey, and points for 0.3 ms exposure images are colored black. X-errors represent the standard deviation of turbidity measurements (n=3) from **Supplementary figure 7**. Y-errors represent the standard deviation of pixel values in each well image.

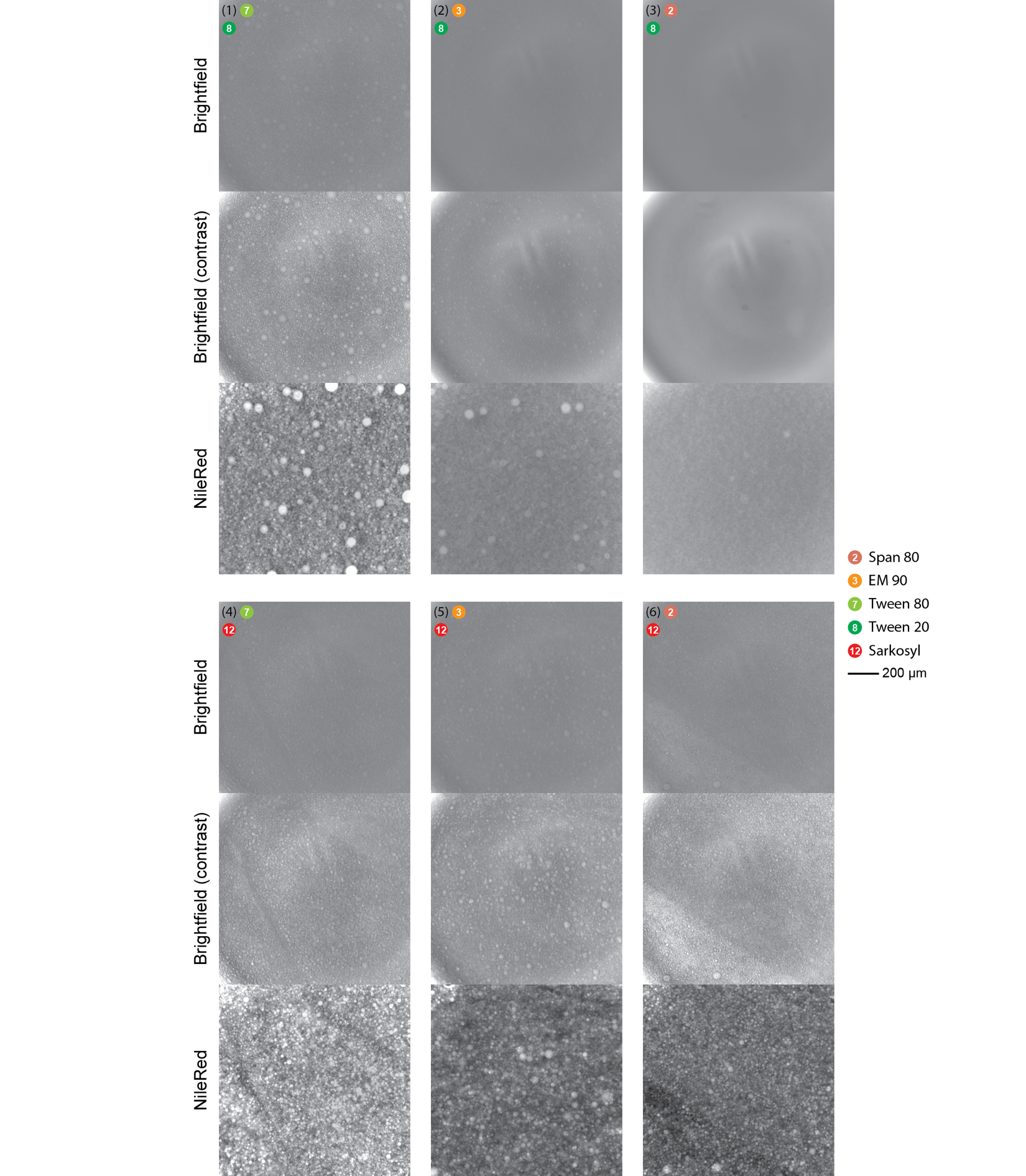

**Supplementary Figure 13: *Fluorescence microscopy images of selected emulsions.***

Brightfield, brightfield with increased contrast, and red fluorescence images of octanol/aqueous emulsions stabilized by surfactant combinations. Images and numbers correspond to the images in **Figure 2D**. Surfactants for 1-octanol are indicated by the number and circle to the right of the image number. All surfactants are 5% (w/v). 1-octanol is labeled with Nile Red.

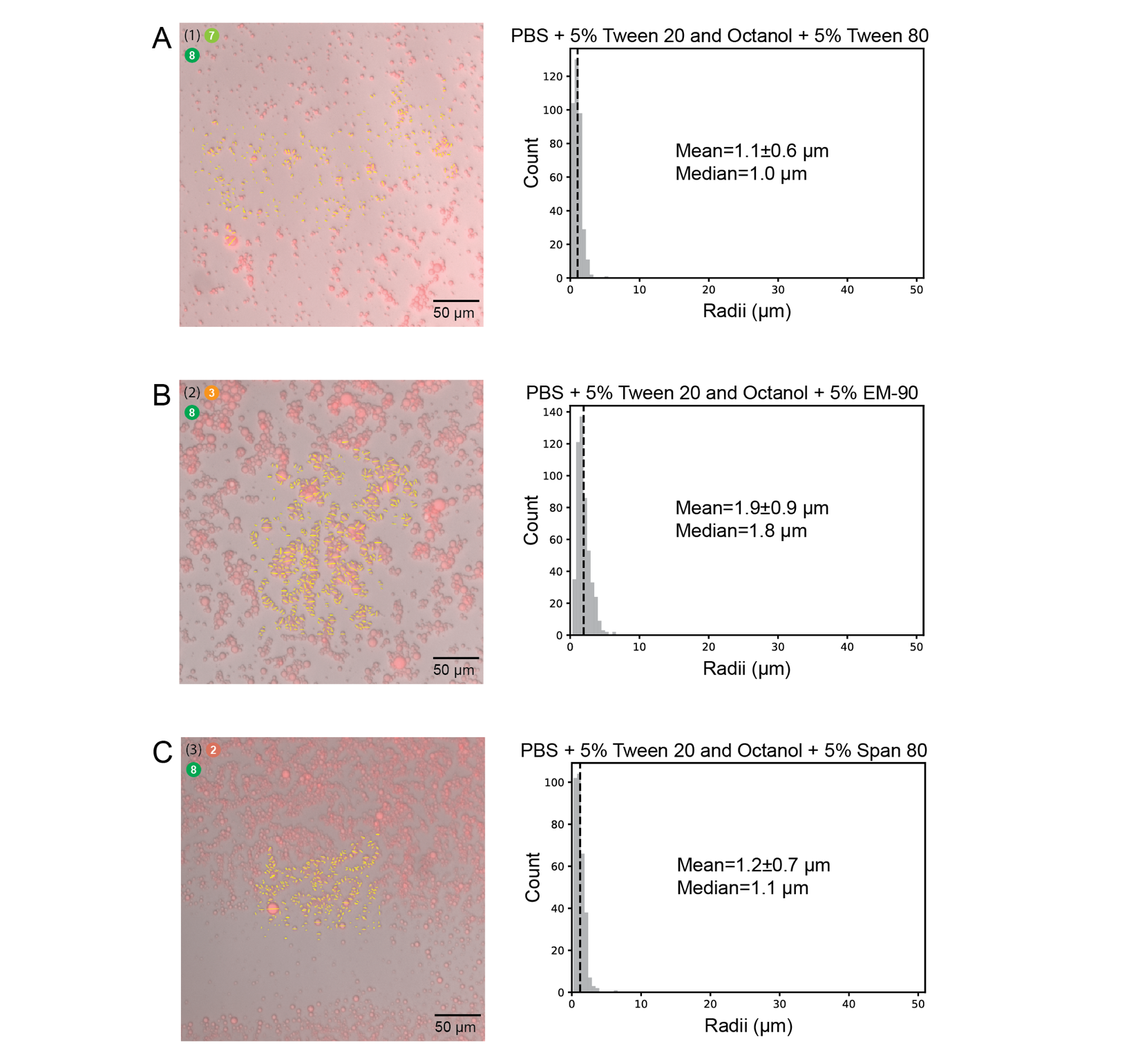

Legend on following page.

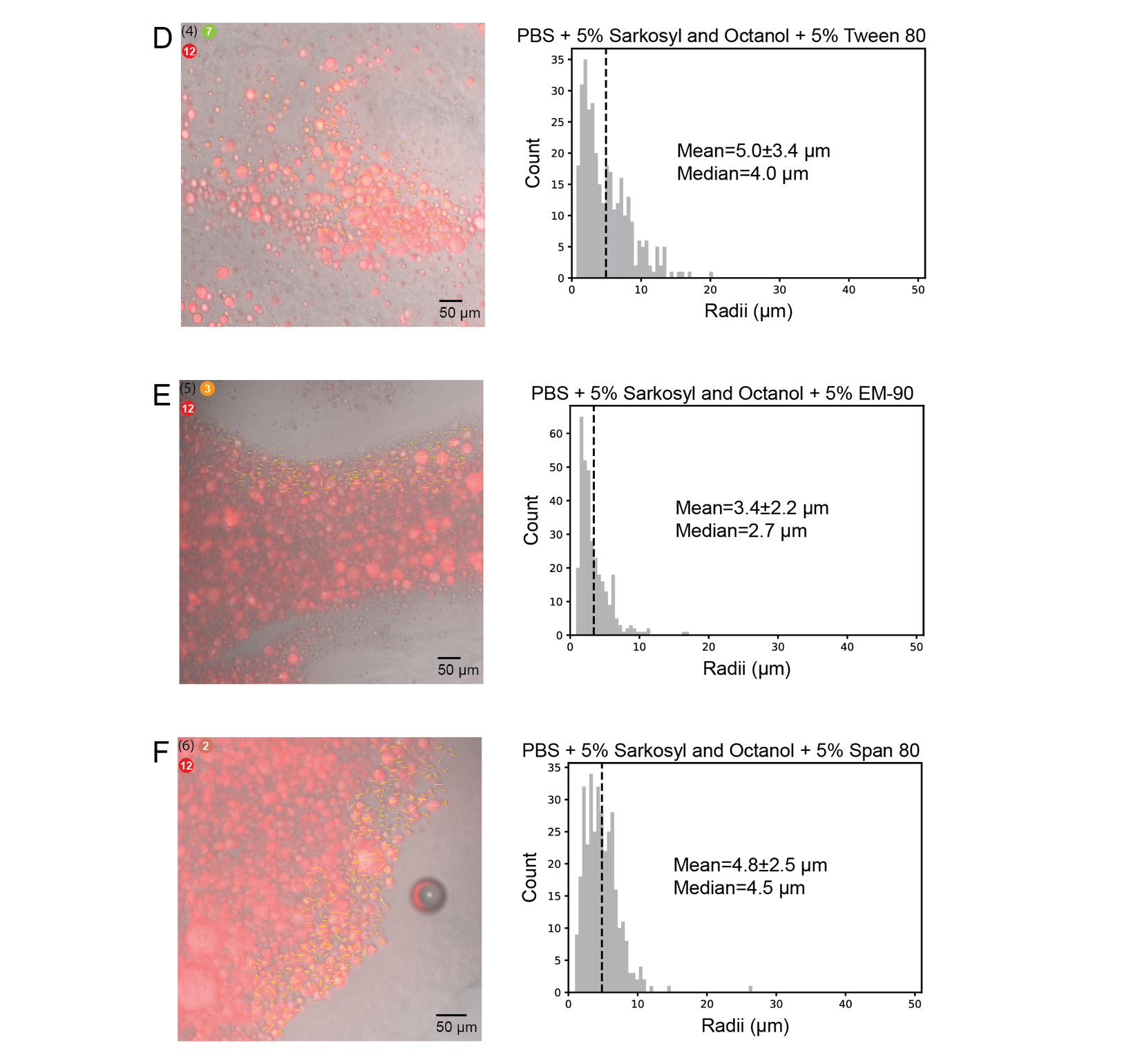

**Supplementary Figure 14: *Quantifying droplet size distribution in an octanol/aqueous emulsion.***

Merge fluorescence and brightfield images of diluted octanol/aqueous emulsions used for droplet sizing and histograms of droplet sizes for the six surfactant combinations in from **Figure 2D** (**A-F**). Surfactant keys (image, top left) are the same as in figure 2D and match the histogram titles. Merge images from manual droplet diameter measurement in ImageJ are shown with all line measurements overlaid (yellow). Radii calculated from measurements are plotted as histograms for each condition. Median radii are indicated with a dashed black line.

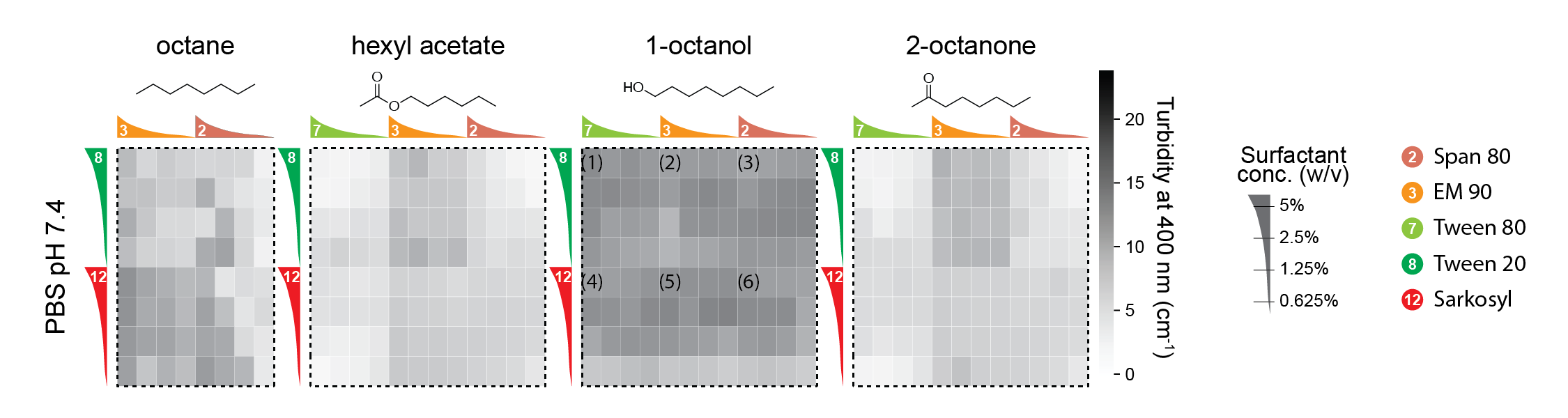

**Supplementary Figure 15: *Plate reader turbidity measurements for surfactant combinations as a function of concentration at 2 hours.***

Plate reader turbidity measurements after 2 hours for one aqueous buffer (PBS pH 7.4) and 4 hydrocarbon solvents in the presence of 2 surfactants for the aqueous phase and 2-3 surfactants for the hydrocarbon phase added at concentration of 0.625-5% (w/v). Surfactant labels (right) reproduced from **Figure 2B**. Data after 24 hours shown in **Figure 2E**. Numbers correspond to selected conditions and images in **Figure 2D**.

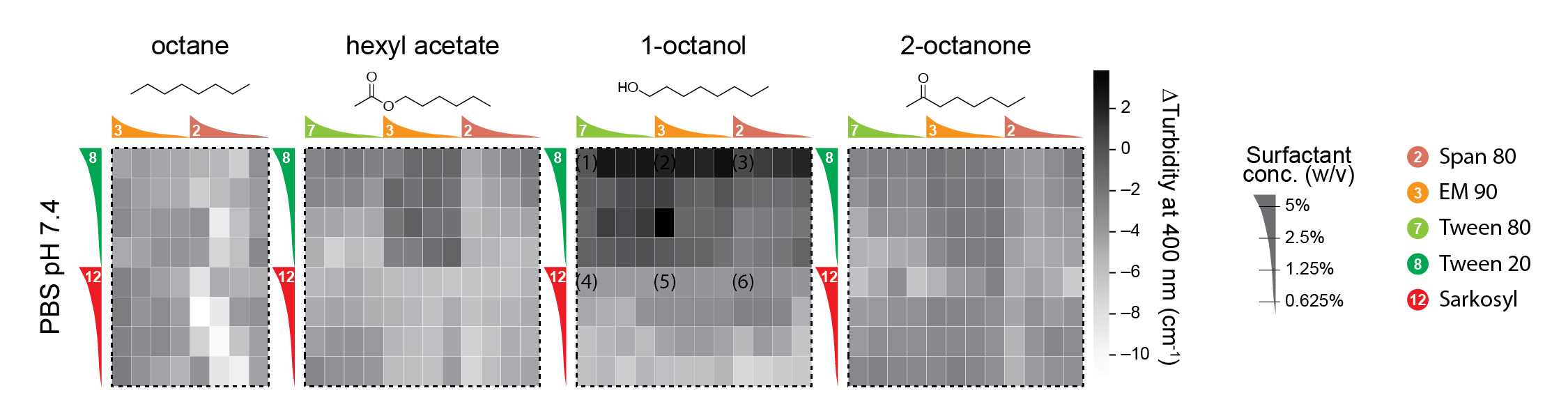

**Supplementary Figure 16: *Changes in plate reader turbidity measurements for surfactant combinations as a function of concentration between 24 hours vs. 2 hours.***

Change in plate reader turbidity measurements between 2 hours and 24 hours for one aqueous buffer (PBS pH 7.4) and 4 hydrocarbon solvents in the presence of 2 surfactants for the aqueous phase and 2-3 surfactants for the hydrocarbon phase added at concentration of 0.625-5% (w/v). Surfactant labels (right) reproduced from **Figure 2B**. Numbers correspond to selected conditions and images in Figure 2D.

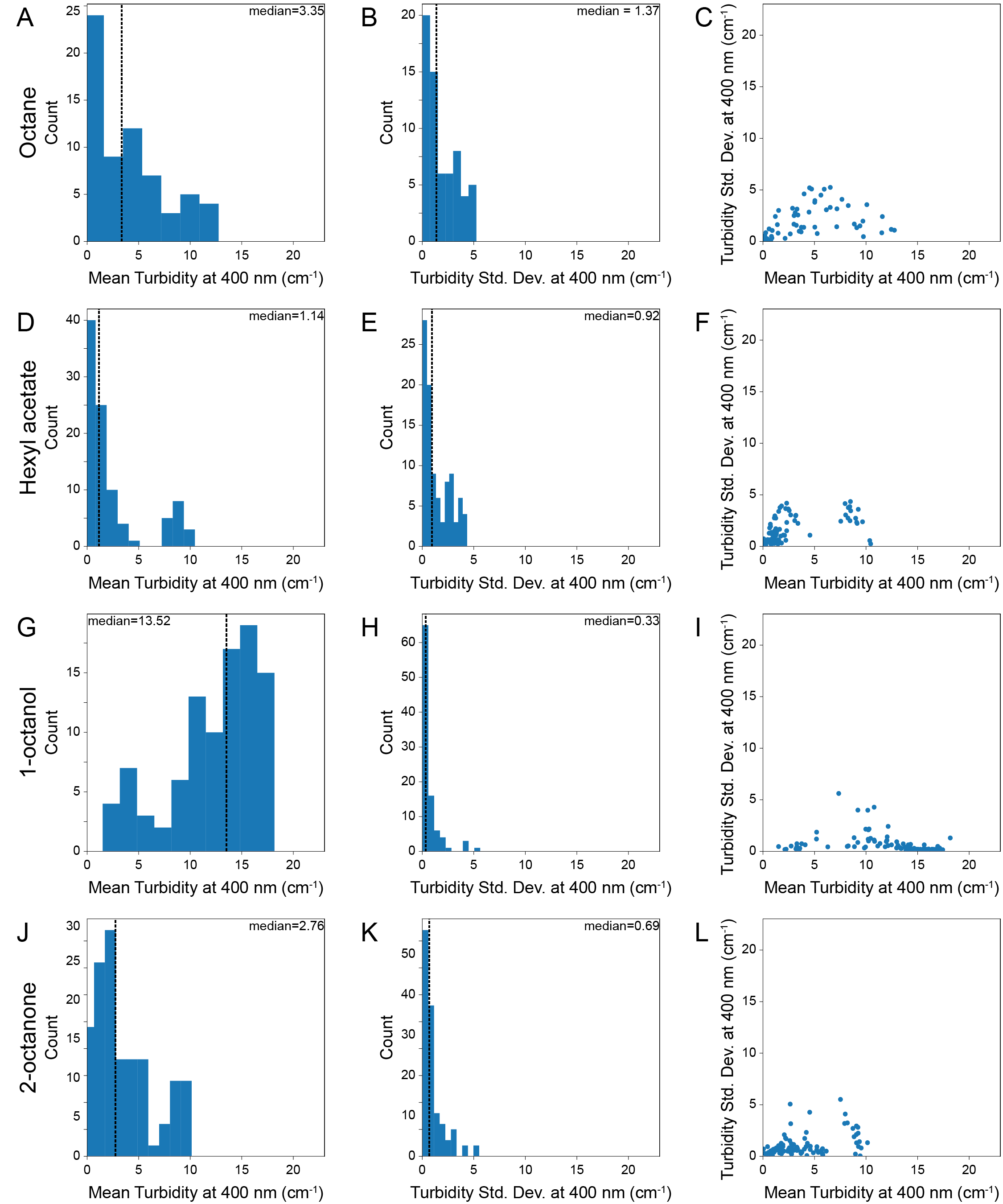

**Supplementary Figure 17: *Reproducibility of plate reader turbidity measurements across wells for optimal hydrocarbon solvent/surfactant combinations at 24 hours.*** Plate reader turbidity measurements after 24 hours for one aqueous buffer (PBS pH 7.4) and 4 hydrocarbon solvents corresponding to data in in **Figure 2E**. Histogram of mean turbidity (n=3), histogram of turbidity standard deviation (n=3), and scatter plot of mean *vs.* standard deviation for combinations of surfactants at 5% (w/v) in octane (**A**-**C**), hexyl acetate (**D**-**F**), 1-octanol (**G**-**I**), and 2-octanone (**J**-**L**).

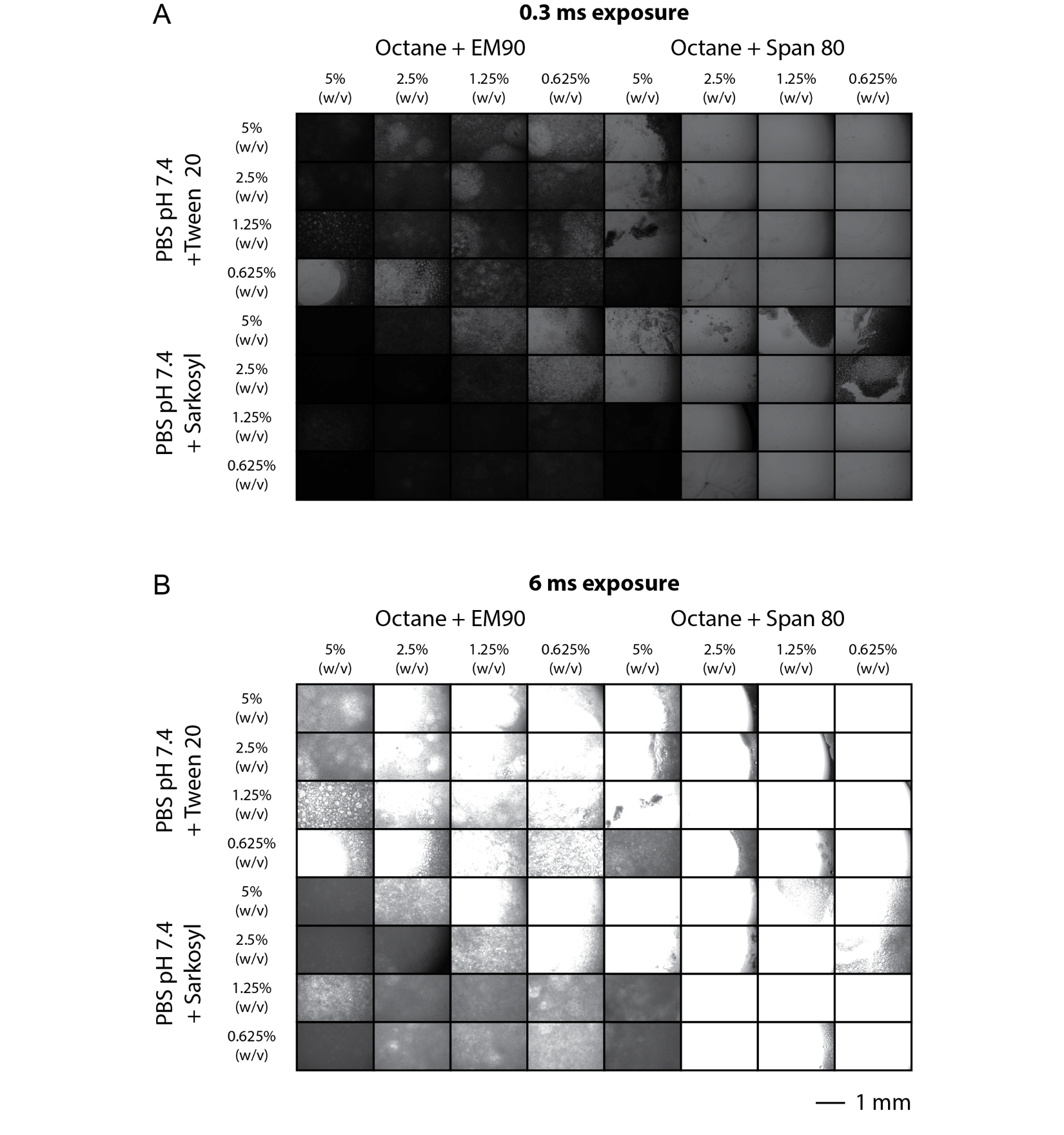

Legend on following page.

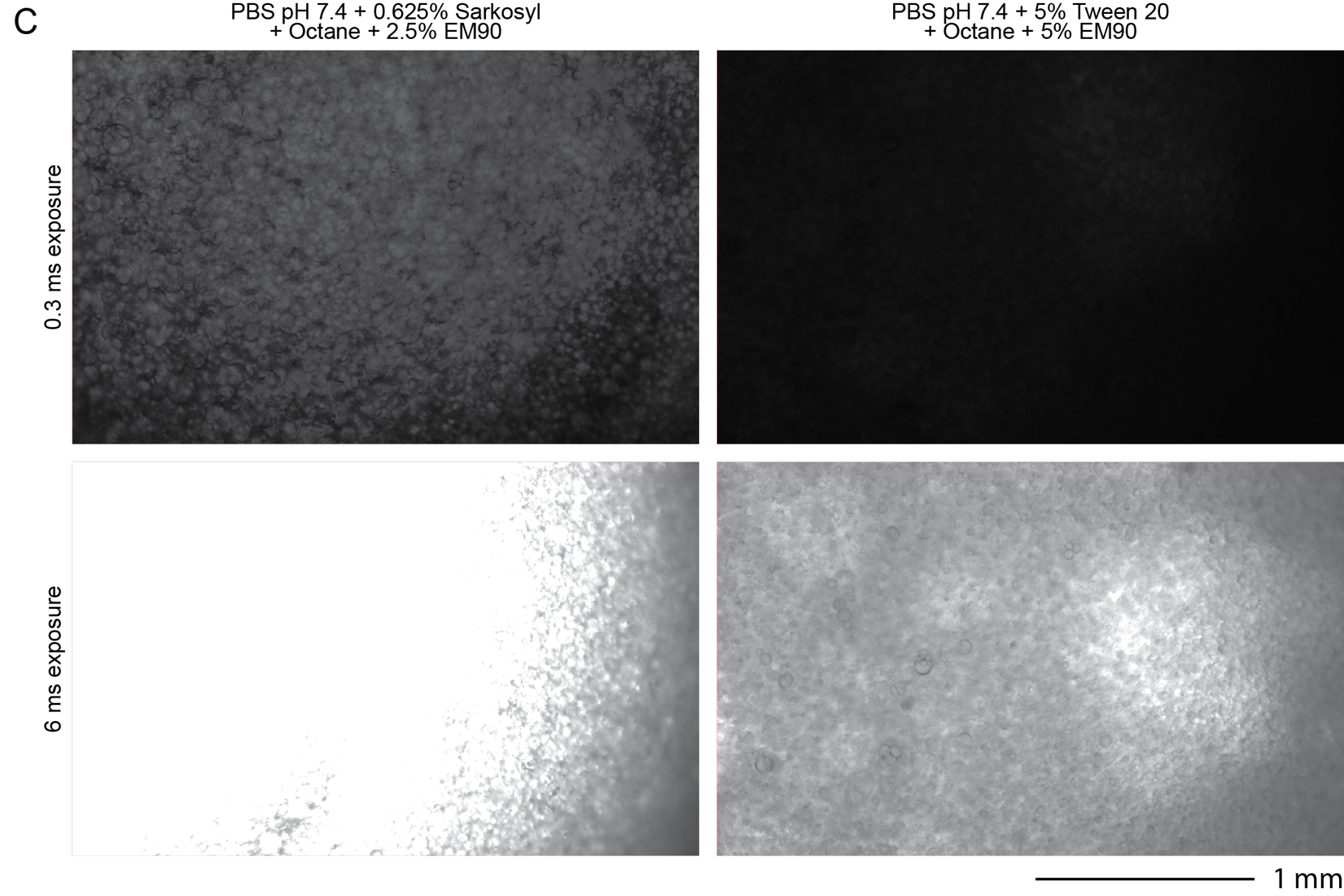

**Supplementary Figure 18: *Well images for optimized octane/surfactant combinations at multiple concentrations at 24 hours.***

**A)** 0.3 ms exposure and **B)** 6.0 ms exposure images of wells from plate reader turbidity assay to screen surfactant at varying concentration for aqueous-octane emulsions. Scale bar is 1 mm. Images correspond to data in **Figure 2E**. **C)** Images of selected conditions. Scale bar: 1 mm.

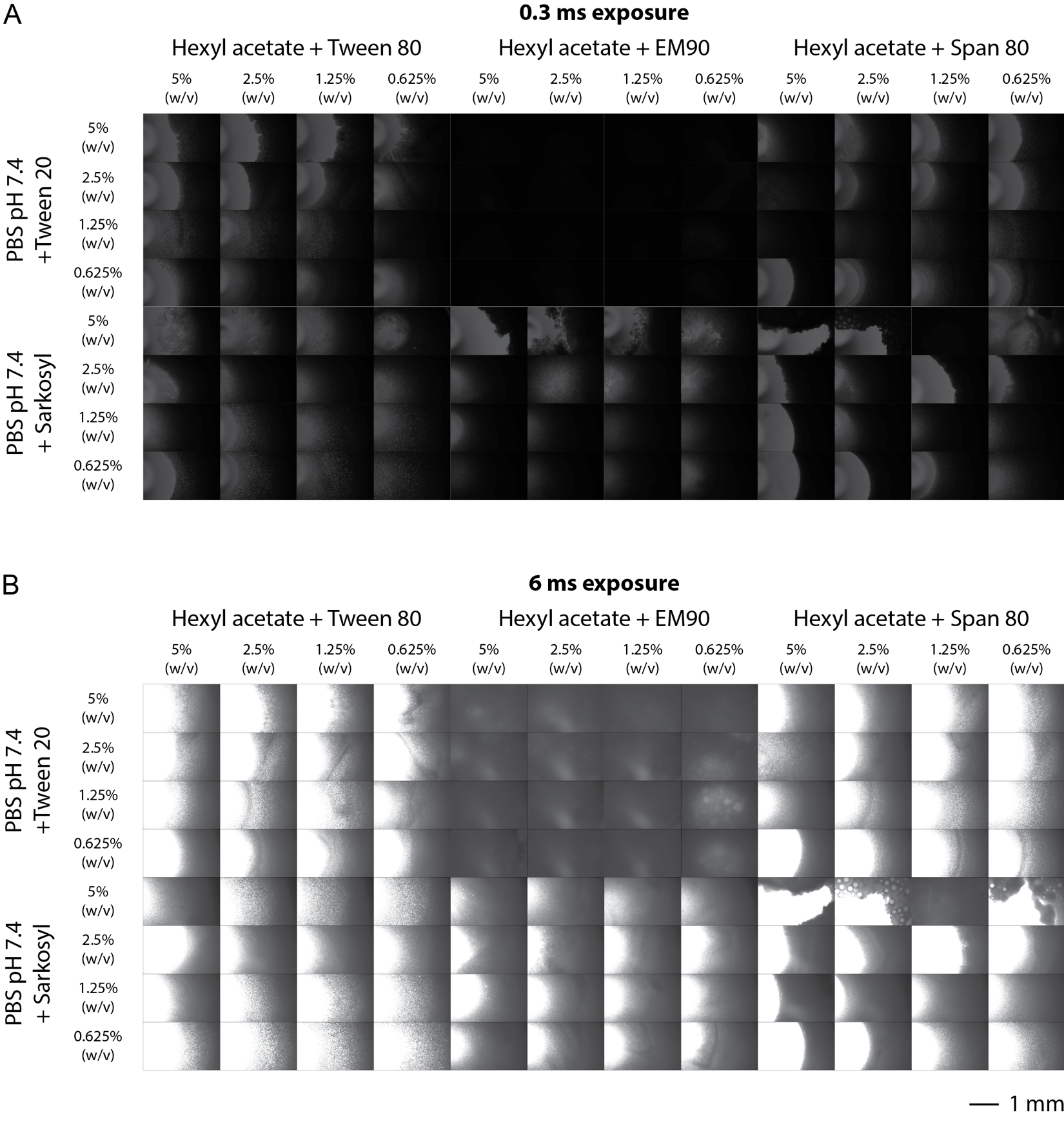

Legend on following page.

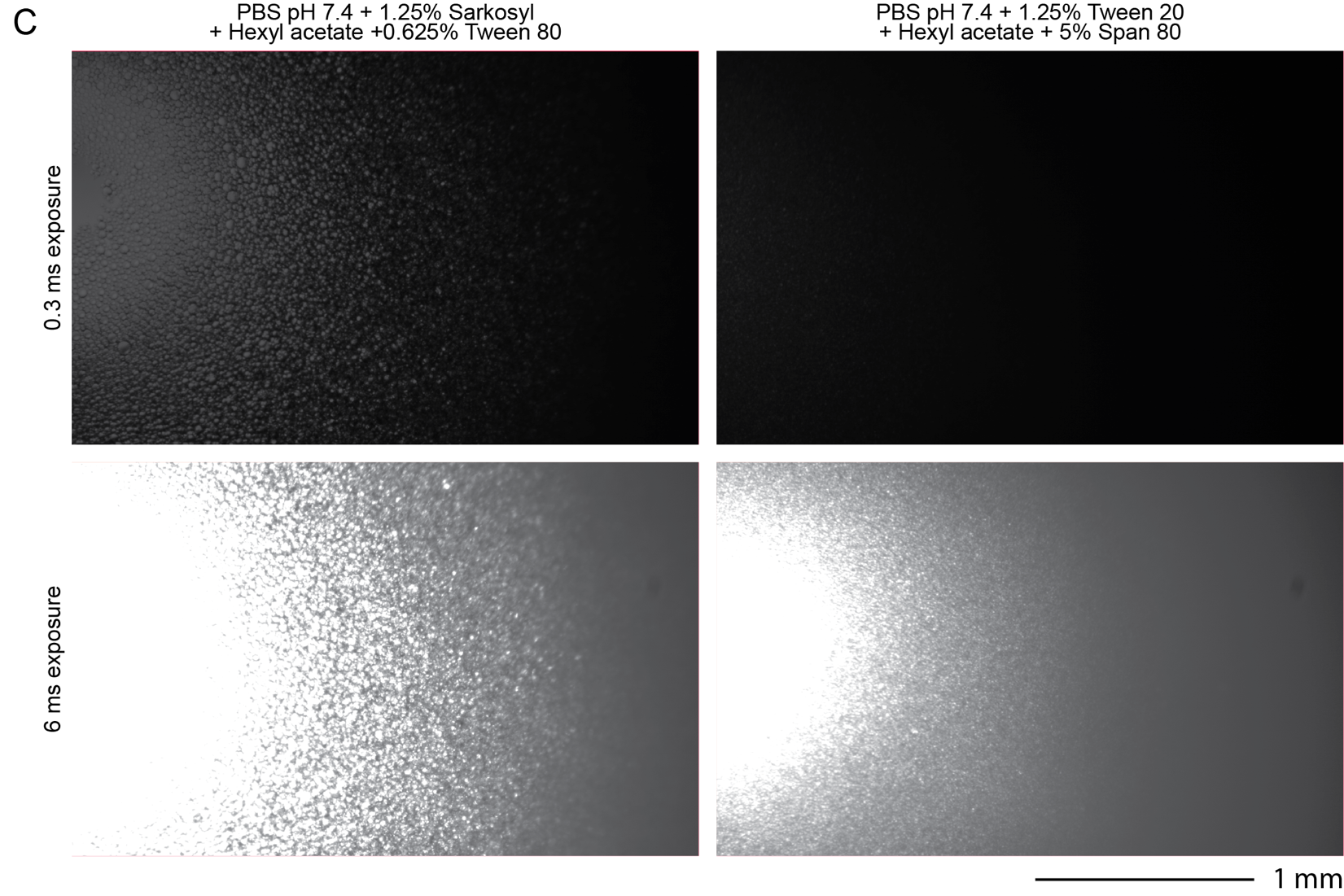

**Supplementary Figure 19: *Well images for optimized hexyl acetate/surfactant combinations at multiple concentrations after 24 hours.***

**A)** 0.3 ms exposure and **B)** 6.0 ms exposure images of wells from plate reader turbidity assay to screen surfactant at varying concentration for aqueous-hexyl acetate emulsions. Scale bar is 1 mm. Images correspond to data in **Figure 2E**. **C)** Images of selected conditions. Scale bar: 1 mm.

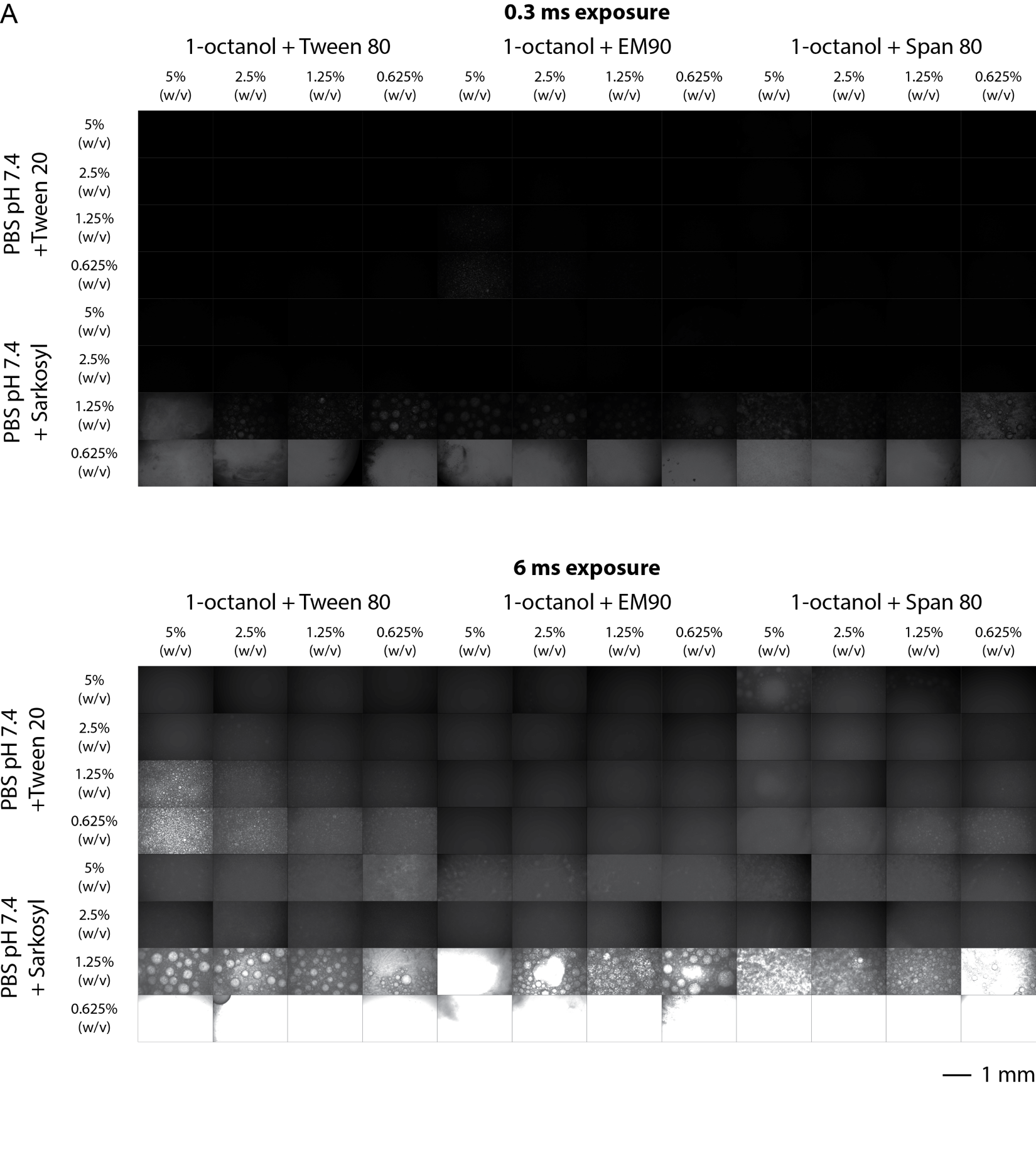

Legend on following page.

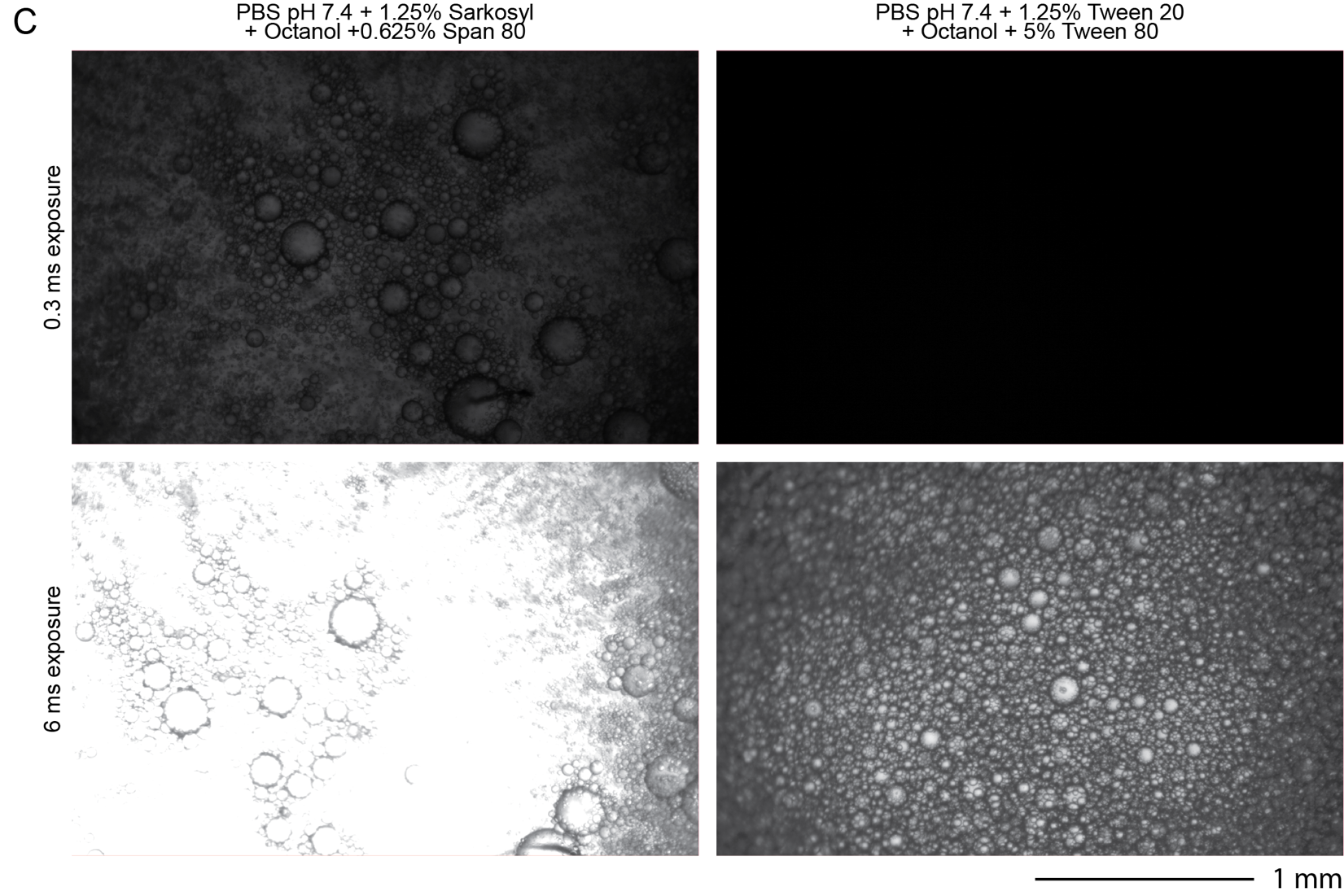

**Supplementary Figure 20: *Well images for optimized 1-octanol/surfactant combinations at multiple concentrations after 24 hours.***

**A)** 0.3 ms exposure and **B)** 6.0 ms exposure images of wells from plate reader turbidity assay to screen surfactant at varying concentration for octanol/aqueous emulsions. Scale bar is 1 mm. Images correspond to data in **Figure 2E**. **C)** Images of selected conditions. Scale bar: 1 mm.

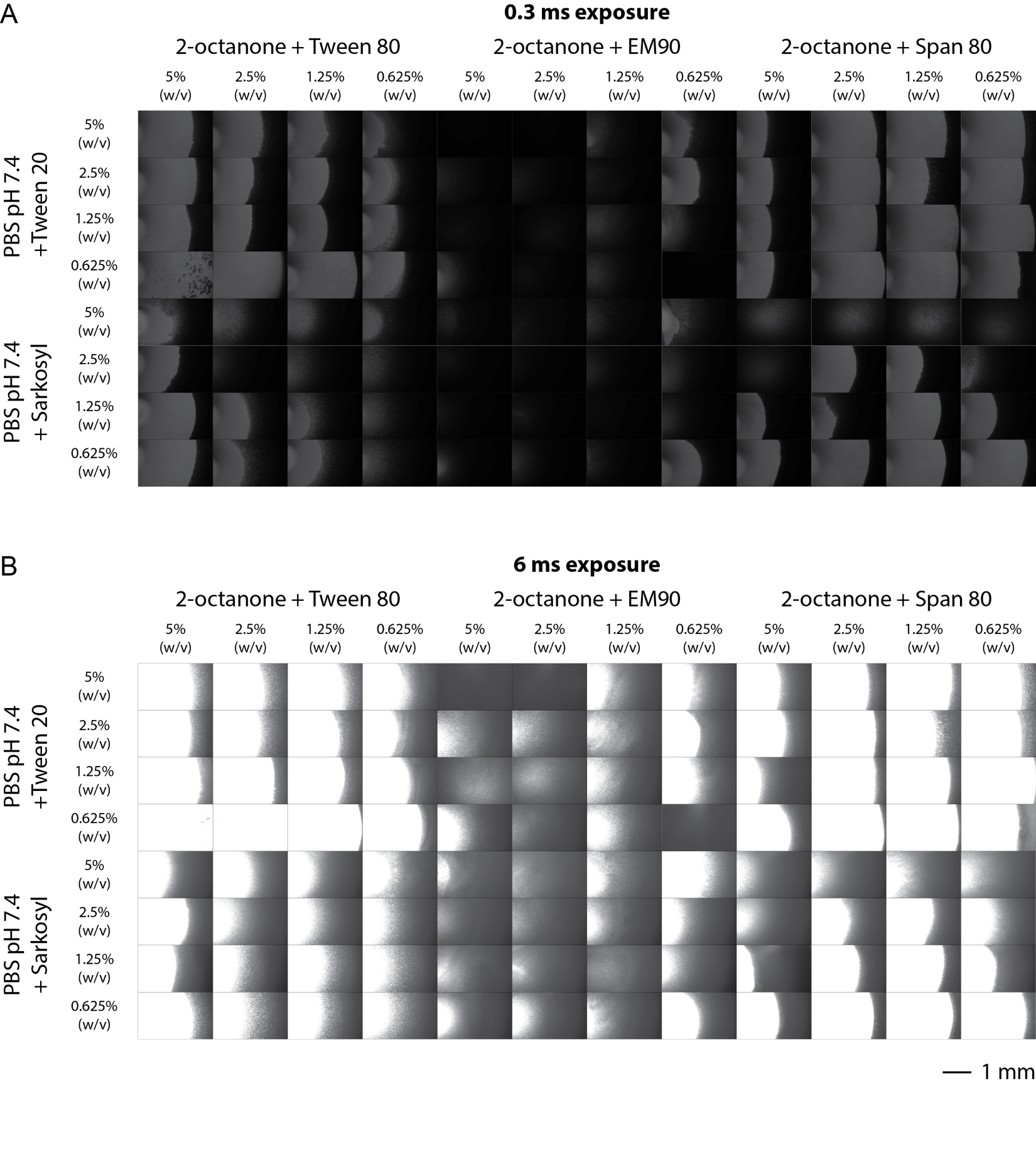

Legend on following page.

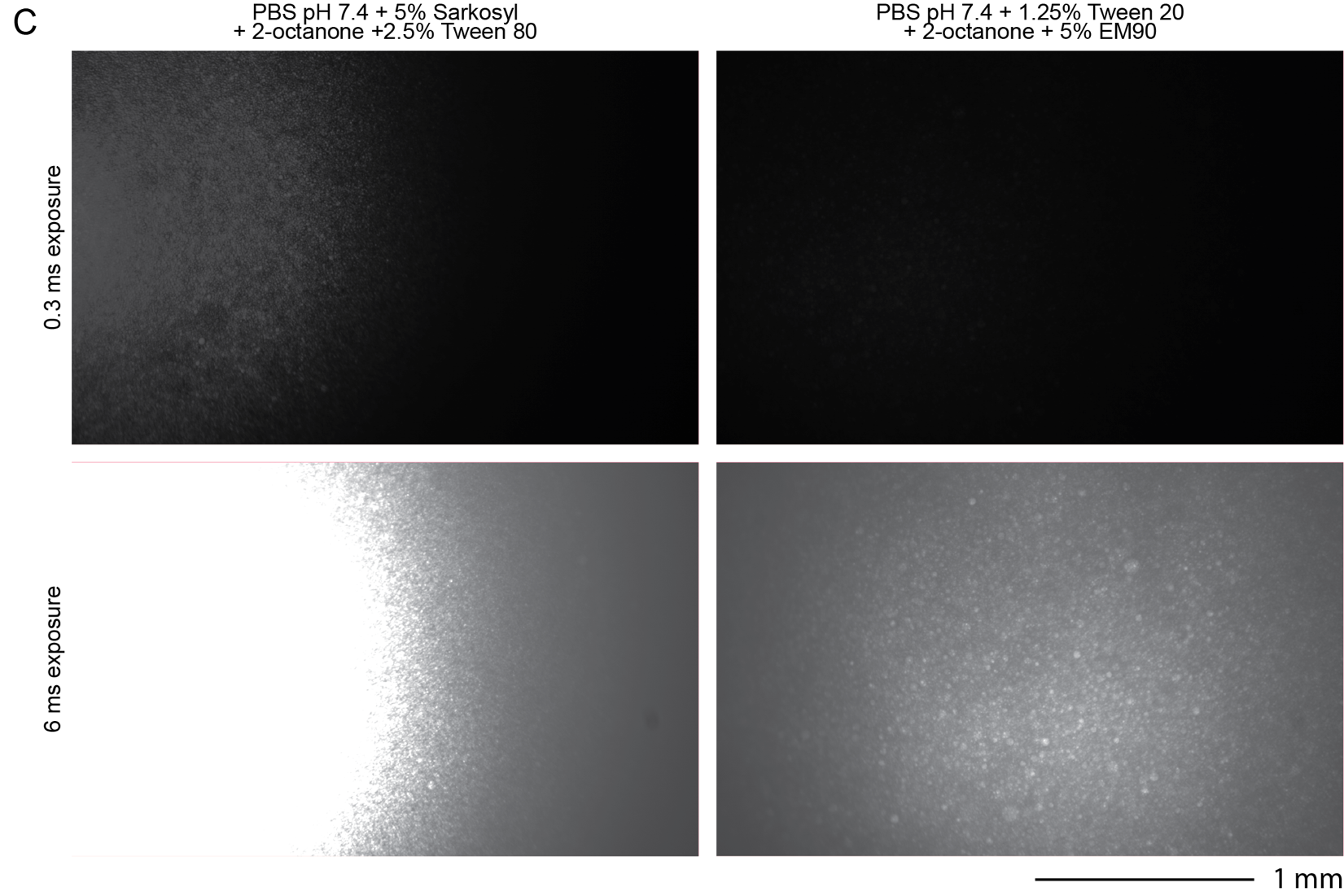

**Supplementary Figure 21: *Well images for optimized 2-octanone/surfactant combinations at multiple concentrations at 24 hours.***

**A)** 0.3 ms exposure and **B)** 6.0 ms exposure images of wells from plate reader turbidity assay to screen surfactant at varying concentration for aqueous-octanone emulsions. Scale bar is 1 mm. Images correspond to data in **Figure 2E**. **C)** Images of selected conditions. Scale bar: 1 mm.

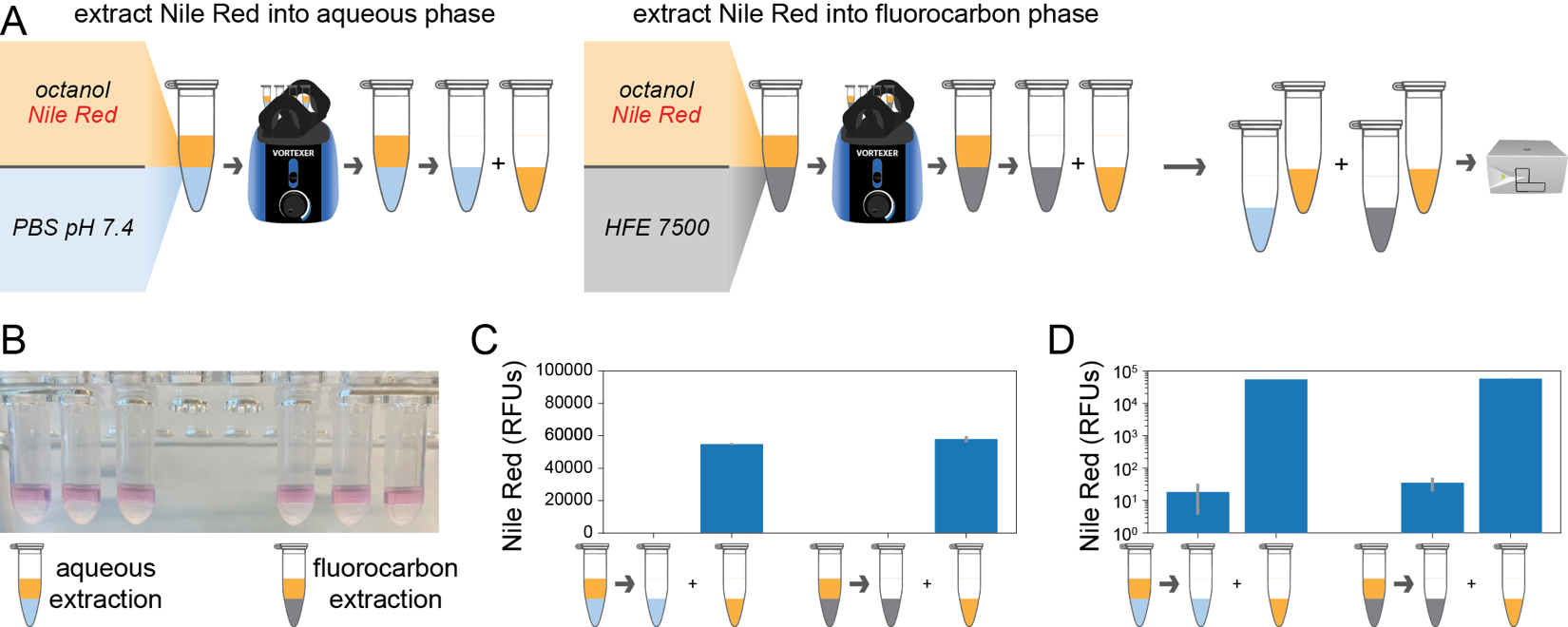

**Supplementary Figure 22: *Nile Red selectively partitions into 1-octanol.***

**A)** Workflow for testing selective partitioning of Nile Red by extraction and fluorescence measurement in a plate reader. **B)** Images of extraction samples after vortexing. Octanol phase is less dense that both aqueous and fluorocarbon phases. **C)** Mean fluorescence measurements for extraction for three experimental replicates. Error bars represent the standard deviation (n=3). **D)** Mean fluorescence from C plotted on Log10 scale.

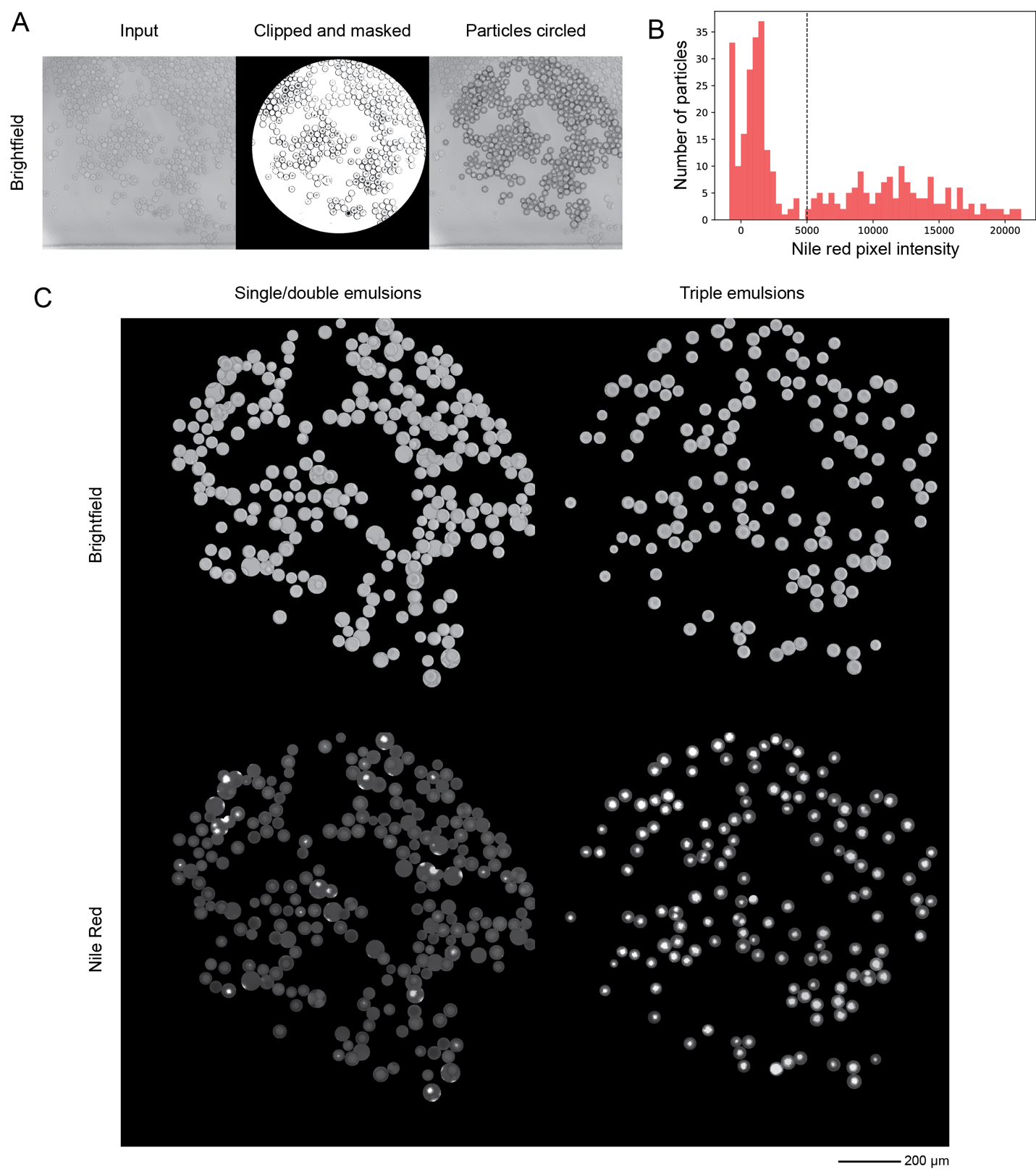

**Supplementary Figure 23: *Automated detection of triple emulsions.***

**A)** Droplets are detected in brightfield and fluorescence microscopy images with a custom python script using the PIL and cv2/OpenCV libraries. Brightfield images are clipped and circles corresponding to the droplets are detected for masking. **B)** Histogram of Nile Red fluorescence intensities (from octanol phase) for individual droplets. After masking individual droplets, the pixel intensities for Nile Red fluorescence are integrated for smaller circle (radius = radius - edge_band) and a user-defined threshold (dashed line) is applied. Droplets with integrated fluorescence intensities above the threshold are classified as triple emulsions. **C)** Example image after classification. All detected and masked droplets are shown in images that represent their classification bin: triple emulsion or not triple emulsion. Brightfield and Nile Red fluorescence images are shown. Scale bar: 200 µm.

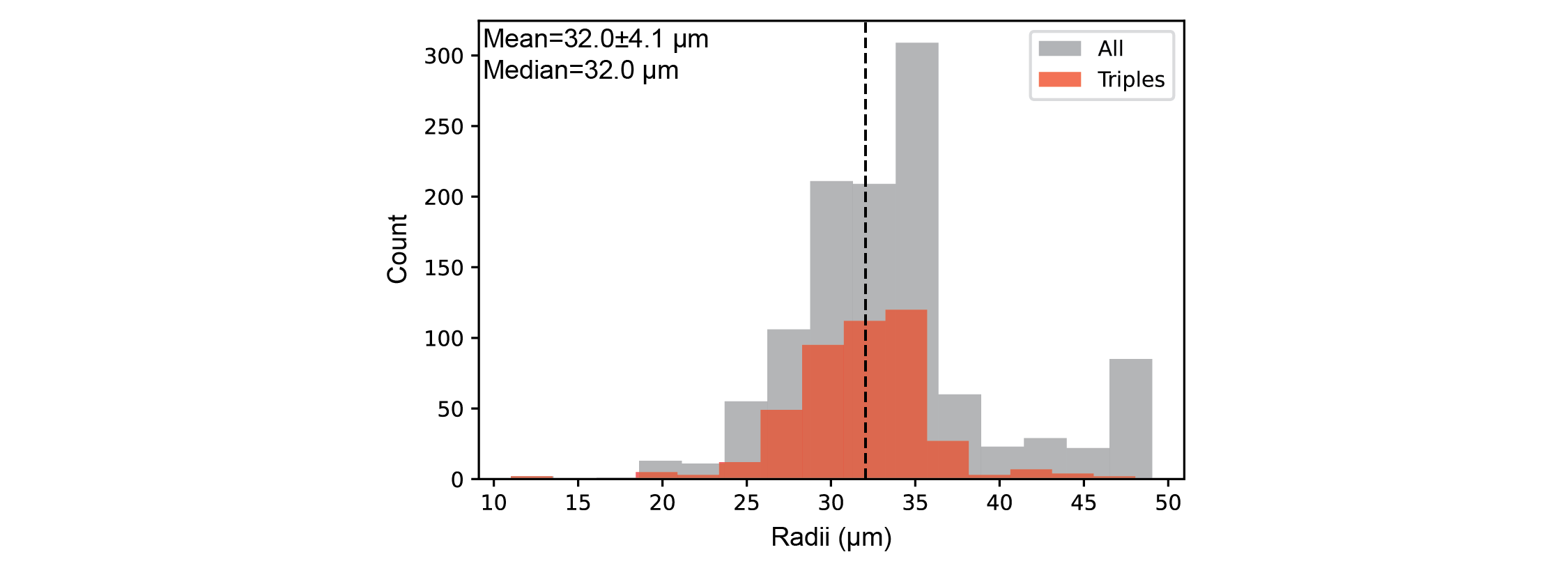

**Supplementary Figure 24: *Distribution of droplet sizes.***

Distribution of droplet radii from automated detection for all droplets (gray) and triple emulsions (red). Median droplet size is indicated with a dashed line. Droplets correspond to data for **Figure 3**. Image data from 2 hours after generation was used for analysis.

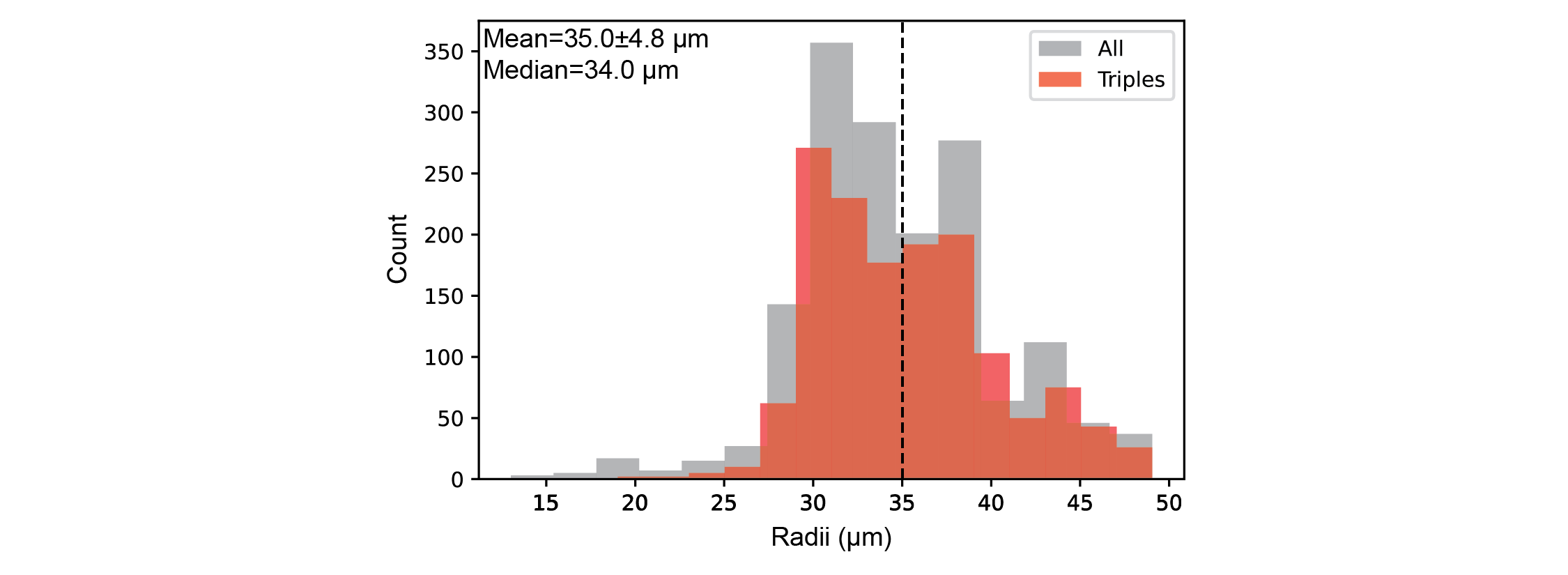

**Supplementary Figure 25: *Distribution of droplet sizes with IVTT reagents.***

Distribution of droplet radii from automated detection for all droplets (gray) and triple emulsions (red). Median droplet size is indicated with a dashed line. Droplets correspond to data for **Figure 4G**, 100:100 (µl/hr) flow ratio, and 2 hour incubation at 37˚C. Image data from 2 hours after generation was used for analysis.

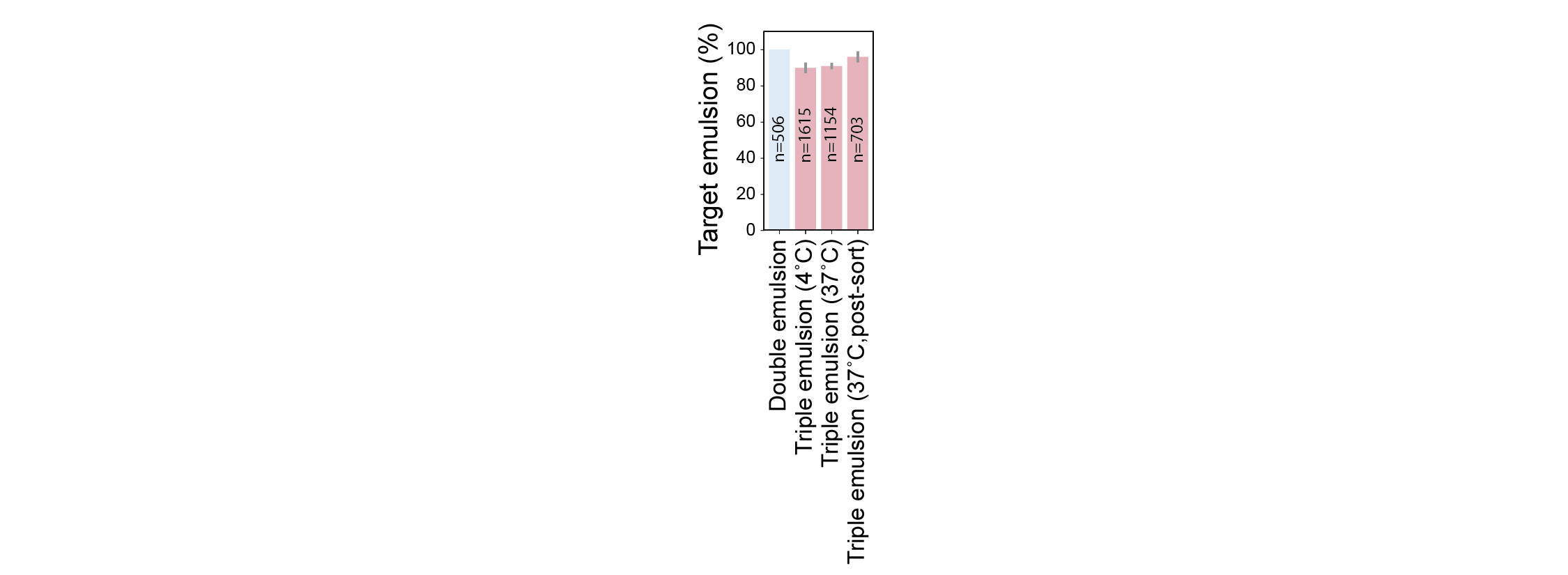

**Supplementary Figure 26: *Double and Triple Emulsion populations in samples for FACS analysis and sorting.***

Percentage of target emulsions within droplet samples produced for FACS sorting and analysis. Bar color indicates target droplet architecture, double (blue) or triple (red). Error bars represent standard deviation in percentages across at least 3 images. No single or triple emulsions were observed in the double emulsion images.

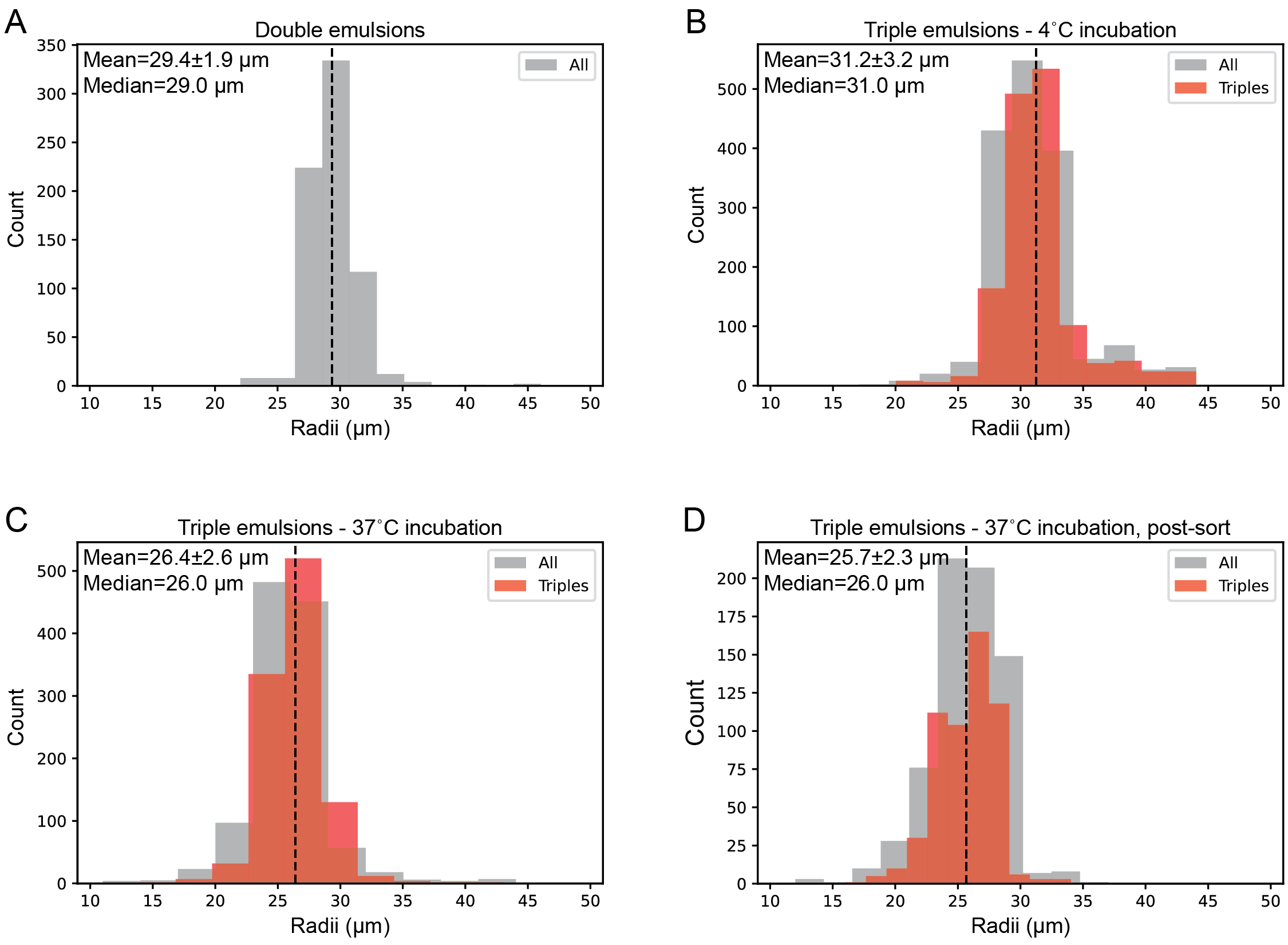

**Supplementary Figure 27: *Distribution of droplet sizes for FACS analysis and sorting.***

Distribution of droplet radii from automated detection for all droplets (gray) and triple emulsions (red) for double emulsions (**A**), triple emulsions incubated at 4˚C (**B**) and 37˚C (**C**), and post-sorting (**D**) droplet populations. Median droplet size is indicated with a dashed line. Droplets correspond to data for all droplets in Figure 5. For unsorted droplets, image data was collected after the incubation step. Sorted droplets were imaged 1 hour after sorting.

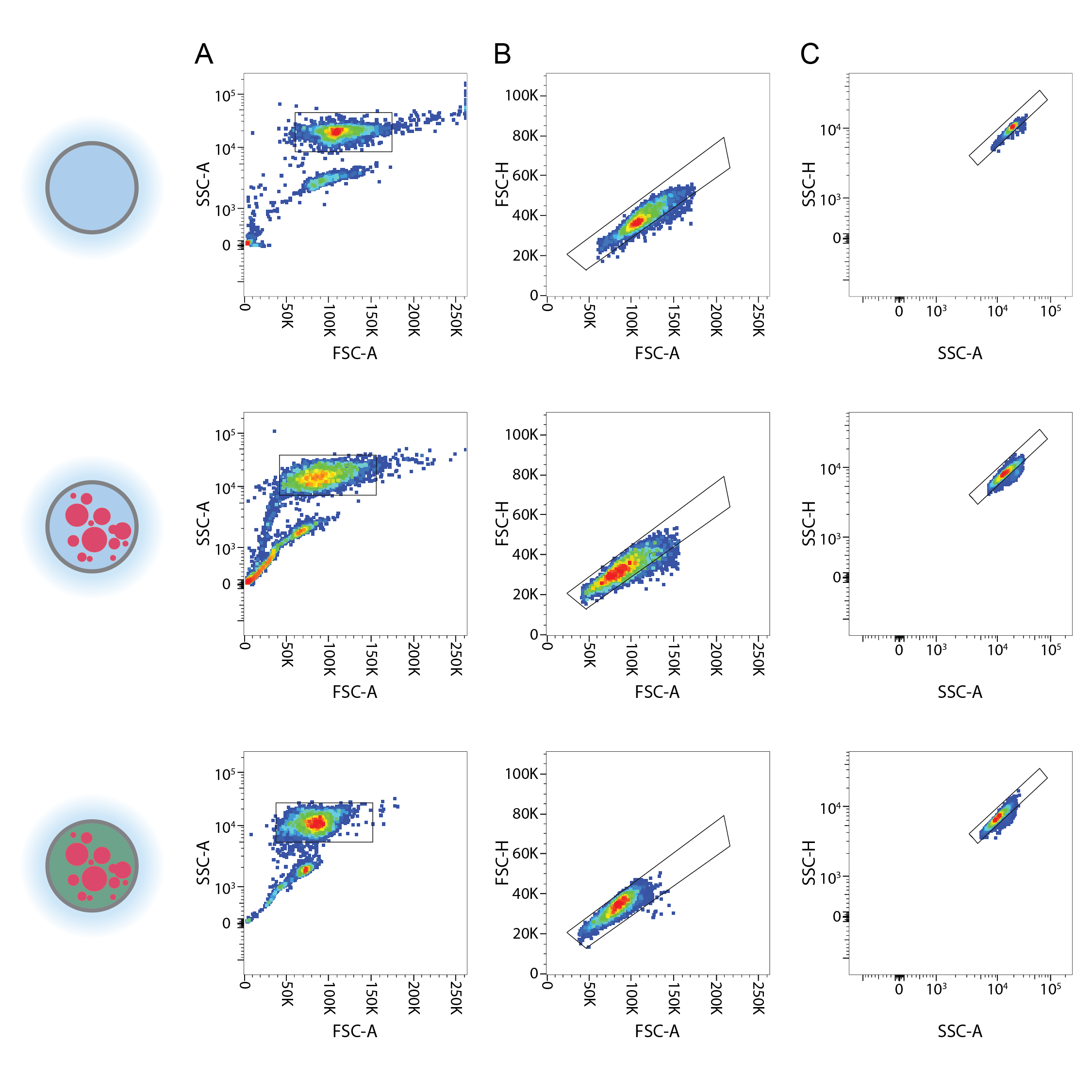

Legend on following page.

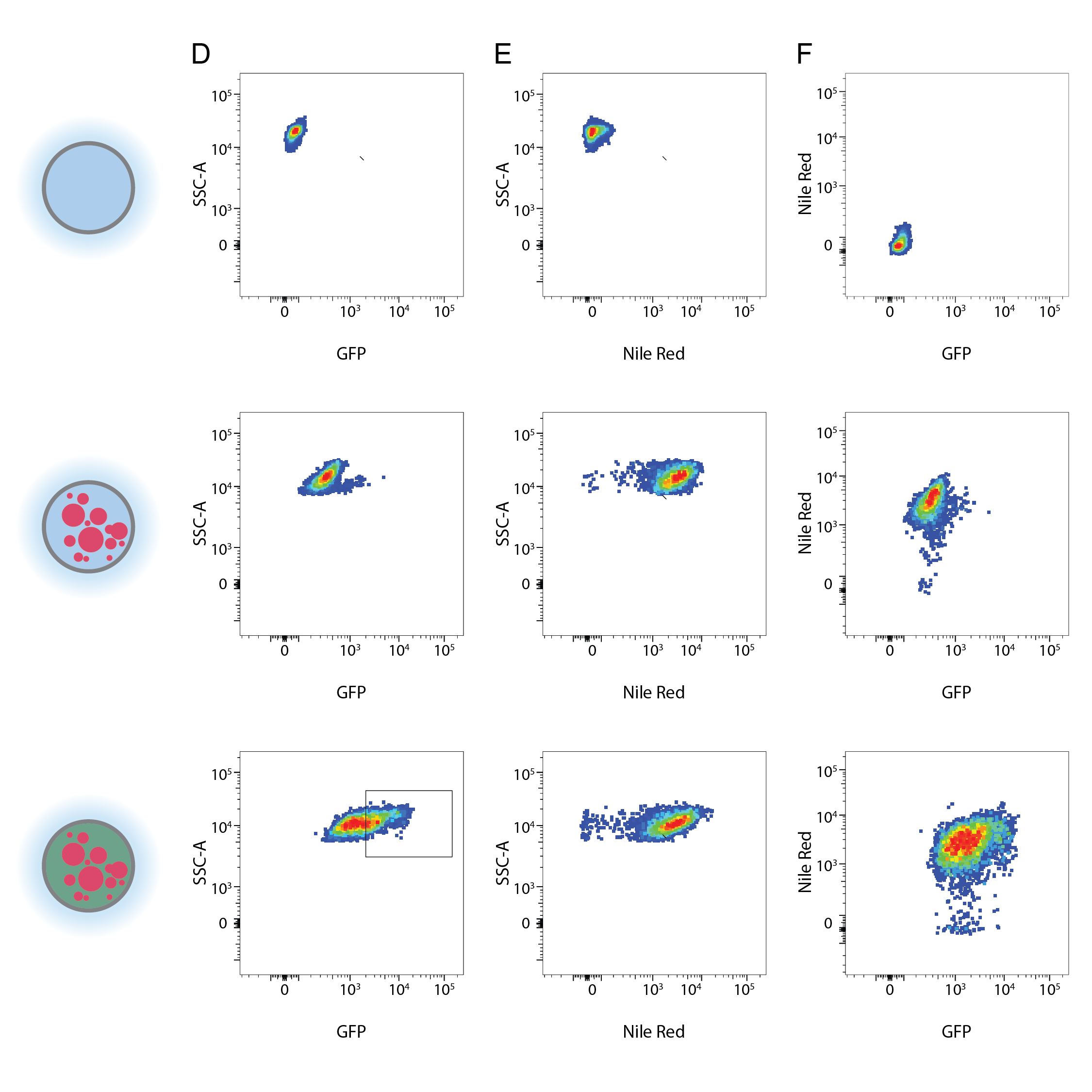

**Supplementary Figure 27: *FACS Analysis of droplet uniformity and fluorescence***

FACS analysis profiles and gates are shown for 10,000 events from samples of double emulsions (top row), triple emulsions with IVTT reagents incubated at 4˚C (middle row), and triple emulsions with IVTT reagents incubated at 37˚C (bottom row). Gates (black outline) were applied sequentially (e.g. all events in B passed the gate in A). See Supplementary Table 4 for event counts. **A)** Using scattering data, droplet picoreactors are selected from satellite oil droplets from a gate on FSC-A *vs.* SSC-A. Singlets are selected from a gate on FSC-A *vs.* FSC-H (**B**) and SSC-A v.s SSC-H (**C**). Droplet fluorescence was analyzed for singlet populations as GFP *vs.* SSC-A (**D**), Nile Red vs. SSC-A (**E**), and GFP vs. Nile Red. A sorting gate was applied to GFP *vs.* SSC-A for triple triple emulsions with IVTT reagents incubated at 37˚C (D, bottom left plot).

**Extended Methods**

***Screening surfactants***

Solvents were purchased and used as provided: octane (Sigma-Aldrich, cat. # 412236), hexyl acetate (Sigma-Aldrich, cat. # 108154), 1-octanol (Sigma-Aldrich cat. # 112615), 2-octanone (Sigma-Aldrich, cat. # O4709), PBS pH 7.4 (Gibco, cat. # 10010031). Surfactants were purchased as powders or pure liquids except for CTAB 25% aqueous solution (Sigma-Aldrich,cat. # 292737) and NP-40 10% aqueous solution (Thermo Scientific, cat. # 85124): Benzalkonium chloride (Sigma-Aldrich, cat. # 12060), CHAPS hydrate (Sigma-Aldrich, cat. # C3023), EM 90 (Evonik), Sarkosyl (IBI Scientific, cat. # IB07080), SB3-10 (Sigma-Aldrich, cat. # D4266), SDS (Sigma-Aldrich, cat. # L3771), Span 20 (Sigma-Aldrich, cat. # S6635), Span 80 (Sigma-Aldrich, cat. # S6760), Span 85 (Sigma-Aldrich, cat. # S7135), Triton CG-110 (Sigma-Aldrich, cat. # STS0005), Triton X-100 (Sigma-Aldrich, cat. # T8787), Tween 20 (Fisher BioReagents, cat. # BP337), Tween 80 (Sigma-Aldrich, cat. # P1754). For each solvent-surfactant combination, 0.5 g of surfactant was measured in a 15 mL polypropylene conical tube (Falcon, cat. # 352196). The volume was brought up to 10 mL with solvent. The tubes were then vortexed for up to 15 minutes. Solutions were determined to be soluble if no solid precipitate or liquid phase separation was observable by eye. Serial 2-fold dilutions were prepared by mixing 5 mL of solvent with surfactant with 5 mL of pure solvent.

Calculating HLB

Where possible, hydrophilic-lipophilic balance (HLB) values for surfactants were taken from literature values of measured and calculated values (see **Table 2**). Otherwise, HLB was calculated according to the method established by Davies^7^.

For benzalkonium chloride, the calculation is as follows:

Because benzalkonium chloride has a mixture of R-groups with different chain lengths, upper and lower bounds were set for a total carbon count of n= 9+18 and 9+8, respectively.

Eqn: base + 4˚ amine – n * hydrocarbon

= 7 + 22 – 27*0.475 = 16.175

or

= 7 + 22 – 17*0.475 = 20.925

For CHAPS, the calculation is as follows:

Eqn: base + 4˚ amine + sulfonate + amide + 3*hydroxyl – 32*hydrocarbon

= 7 + 22 + 37.8 + 2.9 + 3*1.9 – 32 * 0.475 = 60.2

Plate reader turbidity measurements

Emulsions for octane and 1-octanol were prepared in transparent-bottomed polystyrene 96-well plates (Nunc, cat. # 265301). Emulsions for hexyl acetate and 2-octanone were prepared in semi-transparent polypropylene 96-well plates (Genier Bio-One, cat. # 651201). 25 µL of hydrocarbon solvent and 75 µL of PBS solution were added to each well. Surfactant and concentrations were used as indicated in **Figure 2** and **Data Supplement 1-8**. The plates were sealed with silicone mats (Corning, cat # AXYAM2MLRD) and vortexed for 2 minutes at 3000 rpm on a vortexer (Southern Labware, cat. # 120209). The plates were spun down for 30 seconds at 2500 rpm in a microplate centrifuge (Fischer Scientific, cat. # 14-955-300) The silicone mat was removed and replace with an adhesive plate seal. The plate was incubated at room temperature. An empty plate was stacked on top of the sealed plate, which effectively suppressed condensation on the seal. Optical absorbance was measured in an Infinite 200Pro multimode plate reader (Tecan) either continuously (during troubleshooting) or for two timepoints at 2 hrs and 24 hrs (during screening) at wavelengths from 400-800 nm (50 nm intervals), with a bandwidth of 9 nm, and integration over 25 flashes. For continuous reading, the adhesive seal was used for the duration of the experiment. For screen timepoints, the plate seal was removed before each read because hexyl acetate and 2-octanone cause warping and clouding of the seal.

Plate reader data were analyzed and plotted with a custom Python script. Scripts and data are available in https://osf.io/gbq5r/. Briefly, turbidity was calculated from optical absorbance = Log10 (I_0_/I) as described by Reddy et al.^9^, where turbidity = ln(I_0_/I)/path_length = Ln(10) * absorbance / path_length, I_0_ is the intensity of unobstructed light, and I is the intensity of light after passing through the sample. A path length of 0.3 cm was used for all calculations. Values in **Data Supplements 1-8** are presented as these calculated values. Data in figures and supplementary figures are normalized by background subtraction using the condition without surfactants as the background. This background varied between samples primarily due to the type of plate used.

Microscopy reader turbidity measurements

After the 24 hr plate reader measurement, the plate seals were removed and emulsions in each well were imaged with a zoom power stereo microscope (Amscope SM-2T) equipped with a high-speed CMOS camera (ZWO, ASI174MM). For control experiments, images were exposed for 1.8 ms. For screening experiments a 0.3 ms exposure image and a 6.0 ms exposure image were collected. For all experiments, the zoom objective lenses were set to 3.5x magnification.

*Testing Nile Red partitioning*

A stock solution of 200 mM Nile Red (Sigma Aldrich, cat. # 72485) was generated by resuspending 8.4 mg of solid Nile Red in 131 µL of DMSO. 1 µL of 200 mM Nile Red in DMSO was mixed with 1 mL of 1-octanol for a 200 µM Nile Red in 1-octanol solution. 50 µL of this 200 µM solution mixed with 1950 µL of 1-octanol to produce a 5 µM Nile Red in 1-octanol solution. Over three experimental replicates, 200 µL of 1-octanol with 5 µM Nile Red was mixed with 200 µL of PBS pH 7.4 or HFE 7500 (3M, cat. # 7100025016) in a 2 mL tube (USA Scientific, cat. # 1620-2700) and vortexed for 5 minutes at 3000 rpm. The phases were allowed to separate, and 100 µL of each phase was pipetted into a transparent-bottomed polystyrene 96-well plate. Fluorescence was measured on a plate reader with excitation at 560 nm (bandwidth: 20 nm) and emission at 635 nm (bandwidth: 10 nm). Measurements were integrated over 25 flashes with a gain of 80.

*Generating microfluidic devices*

Droplet generators were used as designed in Brower et al., 2020^9^. Briefly, molds for device features were generated on silicon wafers using photolithography. PDMS (MG Chemicals, cat. # RTV615) was cast onto a feature mold and a blank silicon water; after a soft bake (80˚C, 12-15 min), the PDMS was released from the mold. Holes were punched for inlet and outlet ports with a 1 mm biopsy punch (Robbins Instruments, cat. #RBP-10), the PDMS pieces were cleaned with Scotch tape and the featured and blank PDMS slabs were joined and baked at 80˚C for 48-56 hrs. CAD files for photolithography exposure masks are available from the Open Science Framework repository for that reference: <https://osf.io/h4sr9/>.

***Operating microfluidic devices***

*Generating double emulsions*

FACS sortable double emulsions were generated as previously described using a set-up that included the PDMS devices, 4 syringe pumps (Harvard Apparatus, cat. # 70-4511), and a zoom power optical microscope. Aqueous solutions were prepared and filtered with a syringe-driven 0.22 µm filter (Millipore Sigma, cat. # SLGV033RB). dSurf droplet oil (HFE7500 +2% dSurf, Fluigent, cat. # DR-RE-SU-12, inlet 1 on diagram) was used as provided for the fluorinated oil shell. For the inner solution (PBS pH 7.4 + 1% Tween 20, inlets 2 and 3 on diagram), 1 g of Tween 20 was measured in a graduated cylinder. For the outer solution (PBS pH 7.4 + 1% Tween 20 + 2% Pluronic F68, inlet 4 on diagram), 1 g of Tween 20 and 2 g of Pluronic F68/Kolliphor P 188 (Millipore Sigma, cat. # K4894) was measured in a graduated cylinder. For each, the volume was brought up to 100 mL with PBS pH 7.4. A stirbar was added to each graduated cylinder, and the opening was sealed with Parafilm (Bemis, cat. # PM-996). The surfactants were dissolved by overturning and stirring, then the solutions were sterile filtered into fresh PETG storage bottles (Nalgene, cat. # 342020-0030).

For droplet generation, input solutions were loaded into disposable luer lock syringes: inner solution in two 1 mL syringes, dSurf oil in a 3 mL or 5 ml syringe, and outer solution in a 10 mL or 30 mL syringe (Becton Dickinson, cat. # 309628, 309657, 309646, 302995, 302832). Bubbles were removed from each solution by tapping and ejection, then each syringe was capped with a 27-gauge stainless steel dispensing needle (McMaster-Carr, cat. # 75165A688). The needles were inserted into the open end of polyethylene medical tubing (Scientific Commodities Inc., cat. # BB31695-PE/2). The syringes were installed into the pumps, and the tubing was cut sufficiently long to reach the PDMS device on the stage (see Supplementary Figure 1). The device was centered in the microscope field of view and the free ends of the tubing were inserted directly into the ports of the PDMS device after trimming the tubing to reduce slack. The tubing was removed, and the right-hand side of the device was plasma treated to render it hydrophilic. The ports for inner and oil phases were covered with Scotch tape, and the PDMS device array was placed in a plasma cleaner (Harrick Plasma, cat. # PDC-001) with a dry scroll pump (Agilent cat. # IDP3B01), a Type 0536 TC vacuum gauge (Agilent cat. # L6141303), and a vacuum gauge monitor (Harrick Plasma, cat. # PDC-VCG). The chamber was vacated and the three way valve was opened to ambient air with the regulator valve that maintained pressure of 400 mbar. Plasma treatment was performed under “high” setting (30 watts) for 10-12 minutes. After deactivating plasma coils, the chamber was re-pressurized to ambient pressure and the PDMS device array was removed.

Fluid solutions were ejected sufficient to fill the tubing with solution and eliminate air bubbles. An outlet line was connected to the device and the free end was inserted into a fresh 2 mL tube (outlet 5 on diagram) The device was recentered on the microscope, and the outer solution line was connected to the device and the outer solution was flowed through the port for 30 seconds at a flow rate of 4000-5000 µL/. The oil input was set to a flow rate of 600 µl/hr and the tubing was connected to the port. Once the oil was visible in the second first flow focuser the flow rate was cut back to 400 µL/hr and adjusted to between 300-400 µl/hr. Finally, the two inner inputs were set to flow rates of 200 µL/hr each and the lines were connected to the device. When the inner solution was visible in the first flow focuser, each flow rate was decreased to 50-150 µl/hr. Flow rates were periodically adjusted to maintain stable droplet generation or alter the droplet geometry.

For our first attempt at co-encapsulation of octanol and PBS in a droplet picoreactor, the input for one inner solution was replaced with pure 1-octanol.

*Generating triple emulsions with pre-emulsified 1-octanol/aqueous inner solutions*

Octanol and PBS solutions were used as previously generated with the addition of the lipophilic fluorophore Nile Red to label the octanol layer. 2 µL of 200 mM Nile Red in DMSO was mixed with 1 mL of 1-octanol + 5% (w/v) Tween 80 and with 1 mL of 1-octanol + 5% Span 80 for a final concentration of 400 µM. The labeled octanol was pre-emulsified prior to encapsulation in droplets. A 1:1 emulsion of 300 µL PBS pH 7.4 + 5% (w/v) Tween 20 with 300 µL 1-octanol + 5% Span 80 produces a stable, homogeneous emulsion that did not separate or cream on the timescale of droplet generation.

Triple emulsion droplets were generated as described above for single emulsions with the modified inner solutions. PBS + 5% (w/v) Tween 20 was used as inner solution 1 (inlet 2 on diagram) and pre-emulsfied PBS pH 7.4 + 5% (w/v) Tween 20 with 1-octanol + 5% (w/v) Span 80 + 20 µM Nile Red was used as inner solution 2 (inlet 3 on diagram). For a 3:1 emulsion, the cream was allowed to settle in a downward pointing syringe, and the aqueous layer was ejected. During droplet generation, the concentration of octanol varied as a function of flow rate as observed by the optical transmission in the input channel (see Figure 3). For a 1:1 octanol/aqueous emulsion, fluctuations in octanol concentration were not observable by eye under a consistent flow rate. Droplets were kept at room temperature unless otherwise noted.

Plasmid construction and miniprep

Plasmids for GFP expression (SMT000) were a gift from Kara Brower and Akshay Maheshwari. These plasmids were deposited in Addgene under accession number 216849. Plasmids were transformed into DH5alpha E. coli (New England Biolabs, cat. # C2987I) and plated on LB-Agar plates (Fisher Scientific, AC611895000) with 30 µg/mL chloramphenicol (Sigma-Aldrich, cat. # C0378). Single colonies were used to inoculate cultures in 5 mL of LB (Fisher Scientific, cat. # BP1426) + 30 µg/mL chloramphenicol in 14 mL culture tubes (Fisher Scientific, cat. # 14-956-9C). After 18 hours of incubation at 37˚C and 250 rpm shaking, cultures were pelleted by centrifuging for at 3000 rpm for10 minutes at 4˚C in a swinging bucket centrifuge (Eppendorf, 5810 R). The supernatant was discarded and the pellets were stored at -20˚C until they were miniprepped (Qiagen) and eluted into nuclease-free water (Promega, cat. # P1193). Stock plasmid concentrations ranged from 200-300 ng/µL. Chloramphenicol was kept as 1000x 30 mg/mL stocks in pure ethanol and stored at -20˚C until use.

Expressing protein in the presence of hydrocarbon solvents and surfactants

GFP was expressed in the presence of water-immiscible hydrocaron solvents and water-soluble surfactants as a read-out of PURE (New England Biolabs, cat. # E6800) activity under each condition. An expression reaction consisted of 4 µL PURE Part A, 3 µL PURE Part B, 0.2 µL recombinant RNAse inhibitor (Promega, cat. # N2515), 0.4 µL of SMT000 plasmid DNA in nuclease-free water, 2.4 µL of nuclease free water in PCR strip tubes (USA Scientific, cat. # 1402-4700). Reactions were incubated at 37˚C for 2 hours. For testing hydrocarbon solvent compatibility, 10 µL of each solvent was pipetted on top of the PURE reaction. The reactions were overturned several times before incubation. For testing surfactant compatibility with Benzalkonium Chloride, CHAPS, Sarkosyl, Triton X-100, Tween 20, and Tween 80, solutions of 25% (w/v) surfactant were prepared by adding 0.25 g of surfactant to a 1.5 mL tube (USA Scientific, cat. # 1415-2600) and bringing the volume up to 1 mL with nuclease free water. Serial dilutions were prepared by mixing 500 µL of nuclease free water with 500 µL of nuclease free water with surfactant. The PURE reaction mixture was then modified by replacing 2.4 µL of nuclease free water with 0.4 µL of nuclease free water plus 2 µL of surfactant in nuclease free water for a final concentration of 0.625-5% surfactant in the reaction. For each reaction, protein expression was detected by mixing 5 µL of the aqueous phase with 95 mL of PBS pH 7.4 in a 0.5 mL Eppendorf tube (Fisher Scientific, cat. # 13-698-790) and measuring GFP fluorescence intensity on a spectrophotometer fluorimeter (DeNovix, DS-11 FX+) with 470 nm excitation 514-567 nm emission detection.

Generating triple emulsions for expressing protein in biphasic droplet picoreactors with 1-octanol/aqueous emulsion cores

Triple emulsions for expressing GFP in biphasic droplet picoreactors were generated as described above for double emulsions with the following buffer modifications to introduce *in vitro* transcription translation reagents into the droplets and to eliminate inhibitory concentrations of phosphate. An outer solution (inlet 4 on diagram) of 394 µM HEPES + 1% (w/v) Tween 20 + 2% (w/v) Pluronic F68 pH 7.4 was prepared by adding 7 g of HEPES (Sigma Life Sciences, cat. # H3375), 1 g of Tween 20, and 2 g of Pluronic F68/Kolliphor P 188 to a graduated cylinder and bringing the volume up to 90 mL with nuclease-free water. After dissolving the salts and surfactants, the solution was brought to pH 7.4 with 1 M NaOH. The volume was brought up to 100 mL with nuclease-free water. Inner solution 1 (inlet 2 on diagram) was prepared as an emulsification of 1-octanol and stock plasmid solution. 80 µL of SMT000 plasmid in nuclease-free water was mixed with 20 µL of nuclease-free water + 25% (w/v) Tween 20 in nuclease-water then emulsified with 100 µL 1-octanol + 5% (w/v) Span 80 + 20 µM Nile Red by vortexing at 3000 rpm for 5 minutes. Inner solution 2 (inlet 3 on diagram) containing PURE components was prepared by mixing 100 µL PURE Part A, 75 µL PURE Part B, and 5 µL recombinant RNAse inhibitor.

Imaging droplets and emulsions

Images of droplets were imaged on an inverted light microscope (Nikon, Eclipse Ti). Samples consisting of any of single, double, triple emulsions were resuspended in solution by overturning the sample tube, and 10 µL of solution was pipetted onto a Countess chamber slide (ThermoFisher Scientific, cat. # C10228). The light microscope was controlled with a distribution of the MicroManager software package. Images were taken with a combination of brightfield (Semrock, BRFLD-A-NTE-ZERO), green fluorescence (Chroma, 96226, Ex.: AT480/30x, Dichroic mirror: AT505DC, Em.: AT535/40m), and red-orange fluorescence (Chroma, 49004, Ex.: ET545/25x, Dichroic mirror: T565lpxr, Em.: ET605/70m) filter cubes. Exposure times were 10 ms for bright field, 500 ms for green fluorescence (GFP), and 50 ms for red-orange fluorescence (Nile Red). Illumination was provided by a SOLA SE light engine through a 3mm liquid light guide (Lumencor, cat # SOLA SE 5-LCR-SA). A 10x objective lens was used for all images (Nikon, cat# MRD00100). Images were captured on an sCMOS camera (Andor, Zyla 4.2 Plus, VSC-06278). All images were recorded at 1x1 binning.

Fine single emulsions of 1-octanol in aqueous solutions were analyzed manually. Droplet diameters were measured with ImageJ and exported to a .csv file. Radii were calculated and plotted as histograms with a custom python script.

Droplet picoreactor images were analyzed with a custom python script using the PIL and OpenCV libraries. Scripts and data are available at https://osf.io/gbq5r/. Briefly, corresponding brightfield and fluorescence images are provided as input. Brightfield images are subject to a binary threshold filter and vignette masking before individual droplets are identified with the HoughCircle method from the OpenCV library. Individual particles are masked out twice from the red fluorescence image, once for the particle at full radius (full_image) and once with reduced radius (center_image). The average pixel intensity is calculated for the full_ and center_images (full_image_intensity and center_image_intensity, respectively). Triple emulsions are detected as particles where center_image_intensity - full_image_intensity is above a user-set threshold. Parameters for the HoughCircle method were kept consistent for all images. Triple emulsion threshold values for the red fluorescence images were kept consistent between images from the same day to reflect the observed decrease in the fluorescence signal with long incubation times. Binary thresholding and vignette masking parameters were manually modified for each image to ensure accurate droplet detection. Intensity from the green channel was integrated over the droplet to quantify GFP fluorescence signal.

FACS

Droplet sorting was performed following an optimized calibration workflow on a FACSAria II cell sorter (BD). Laser delays were set with 32 µm AccuCount Ultra Rainbow calibration beads (Spherotech, cat. # ACURFP2.5-300-1). The forward scatter threshold was set to ½ of the of the mean FSC-H, the blue laser was set to a delay of zero, the window extension was set to 0 µs, and the number of displayed events was set to 100. As needed, detector voltages (blue laser (488 nm) with B525 detector (525/50 nm), and green laser (532 nm) with G660 detector (660/20 nm)) were modulated bring the signal within the linear range of the detector (<10^5^). The delays were then manually changed with a step size of 1 µs until the detector readout from each laser was maximized. After optimizing the delay value, the delay window was reset to 2 µs, and the forward scatter threshold reduced to 5,000. Area scaling was then set such that the area and height measurements were equivalent, ensuring that the area, which was used in the assay was at least as large as the height. FSC and SSC voltages were adjusted using the 32 µm AccuCount beads to bring the beads above threshold and on-scale.

Double emulsions were resuspended in their outer solution without dilution and 50-100 µL was pipetted into a 1.2 mL polypropylene microtiter tube (Fisher Scientific, cat. # [02-681-376](https://www.fishersci.com/shop/products/fisherbrand-polypropylene-microtiter-tubes-10/02681376)). The samples were well-suspended before placed the tube into the adapter and loading it onto the cell sorter. Samples were agitated during flow with 300 rpm rotation. Forward and side scatter detector voltages were modulated as needed to keep the main population above threshold and on-scale for the SSC-A vs. FSC-A plot. The sorter was setup with the 130uM nozzle and the break-off into single droplets was stabilized using adjustment of the frequency and amplitude and maintained at a gap of 12, per usual manufacturer recommendations.

Sorting was performed in purity mode. Droplet sorting delays were manually calibrated by sorting double emulsions onto a glass slide. At each delay step of 0.03 µs, 50 events detected by a scatter gate in the SSC-A vs. FSC-A plot were sorted onto a glass slide, and the number of recovered emulsions within the drops was counted under optical microscopy. Multiple steps were taken in each direction to establish a gradient. Steps were then taken in the positive direction until the number of recovered decreased. Additional measurements were then taken near the maximum, and the delay with the highest recovery rate was selected. 3-5 measurements of recovery rate were made at the maximum. Typical sort recovery was 75-80%. Fluorescence detector voltages were optimized with triple emulsions containing GFP from in-droplet IVTT and 1-octanol + 5% (w/v) Span 80 + 20 µM Nile Red to center the signal for positive samples between 10^3^-10^5^. GFP was detected with the blue laser (488nm) excitation /detector with 525/50 nm filter, and Nile Red was detected with the green laser (532nm) excitation/detector with 660/20 nm filter. After calibration, all parameters (delays and voltages) were kept consistent for all samples.

For analysis and sorting, samples were loaded as described and analyzed with manually set polygon gates for scattering events in the SSC-A vs. FSC-A plot and consecutive singleton gates in FSC-A vs. FSC-H and SSC-A vs. SSC-H plots. 10,000 events were recorded for each sample. Event rates ranged between 100-800 Hz during sorting, and the sample pressure was modulated to keep the event rate close to 200 Hz. If the event rate fell below 50 Hz, fresh sample with well-resuspended droplets was added to the library tube. A polygon gate for high GFP (>10^3^) was created on the GFP vs. SSC-A plot, and droplets within this gate were sorted into a 1.5 mL Eppendorf tube containing ~200 µL of outer buffer solution.

**References**

(1)	PubChem. *1-Butanol*. https://pubchem.ncbi.nlm.nih.gov/compound/263 (accessed 2024-01-18).

(2)	PubChem. *1-Hexanol*. https://pubchem.ncbi.nlm.nih.gov/compound/8103 (accessed 2024-01-18).

(3)	PubChem. *1-Octanol*. https://pubchem.ncbi.nlm.nih.gov/compound/957 (accessed 2023-09-18).

(4)	PubChem. *2-Octanone*. https://pubchem.ncbi.nlm.nih.gov/compound/8093 (accessed 2024-01-18).

(5)	PubChem. *Hexyl acetate*. https://pubchem.ncbi.nlm.nih.gov/compound/8908 (accessed 2024-01-18).

(6)	PubChem. *Octane*. https://pubchem.ncbi.nlm.nih.gov/compound/356 (accessed 2024-01-18).

(7)	Davies, J. T. A Quantitative Kinetic Theory of Emulsion Type. I. Physical Chemistry of the Emulsifying Agent. In *Proceedings of the International Congress of Surface Activity*; Gas/Liquid and Liquid/Liquid Interface; 1957; pp 426–438.

(8)	Reddy, S. R.; Fogler, H. S. Emulsion Stability: Determination from Turbidity. *Journal of Colloid and Interface Science* **1981**, *79* (1), 101–104. https://doi.org/10.1016/0021-9797(81)90052-7.

(9)	Brower, K. K.; Khariton, M.; Suzuki, P. H.; Still, C.; Kim, G.; Calhoun, S. G. K.; Qi, L. S.; Wang, B.; Fordyce, P. M. Double Emulsion Picoreactors for High-Throughput Single-Cell Encapsulation and Phenotyping via FACS. *Anal. Chem.* **2020**, *92* (19), 13262–13270. https://doi.org/10.1021/acs.analchem.0c02499.

(10)	Brower, K. K.; Carswell-Crumpton, C.; Klemm, S.; Cruz, B.; Kim, G.; Calhoun, S. G. K.; Nichols, L.; Fordyce, P. M. Double Emulsion Flow Cytometry with High-Throughput Single Droplet Isolation and Nucleic Acid Recovery. *Lab Chip* **2020**, *20* (12), 2062–2074. https://doi.org/10.1039/D0LC00261E.
